# Regulation of Vacuole Morphology by PIEZO Channels in Spreading Earth Moss

**DOI:** 10.1101/2020.08.27.269282

**Authors:** Ivan Radin, Ryan A. Richardson, Ethan R. Weiner, Carlisle S. Bascom, Magdalena Bezanilla, Elizabeth S. Haswell

## Abstract

The perception of mechanical force is a fundamental property of most, if not all cells. PIEZO channels are plasma membrane-embedded mechanosensitive calcium channels that play diverse and essential roles in mechanobiological processes in animals^1,2^. PIEZO channel homologs are found in plants^3,4^, but their role(s) in the green lineage are almost completely unknown. Plants and animals diverged approximately 1.5 billion years ago, independently evolved multicellularity, and have vastly different cellular mechanics^5^. Here, we investigate PIEZO channel function in the moss *Physcomitrium patens*, a representative of one of the first land plant lineages. *Pp*PIEZO1 and *Pp*PIEZO2 were redundantly required for normal growth, size, and shape of tip-growing caulonema cells. Both were localized to vacuolar membranes and facilitated the release of calcium into the cytosol in response to hypoosmotic shock. Loss-of-function (*ΔPppiezo1/2*) and gain-of-function (*PpPIEZO2-R2508K* and *-R2508H*) mutants revealed a role for moss PIEZO homologs in regulating vacuole morphology. Our work here shows that plant and animal PIEZO homologs have diverged in both subcellular localization and in function, likely co-opted to serve different needs in each lineage. The plant homologs of PIEZO channels thus provide a compelling lens through which to study plant mechanobiology and the evolution of mechanoperceptive strategies in multicellular eukaryotes.

---

The ability to sense and respond to mechanical forces like gravity, touch, or cell swelling is an ancient property essential for normal cellular function^6^. Externally or internally derived forces can lead to increased lateral membrane tension, activating mechanosensitive (MS) channels and leading to the flow of ions down their electrochemical gradients^7–9^. Members of the PIEZO family of MS ion channels form large trimeric complexes embedded in the plasma membrane that conduct calcium^1–3^. In animals, PIEZO channels are required for the perception of light touch, shear stress, and compressive force, proprioception, brain development, red blood cell volume control, and nociception (reviewed in^1,2^).

PIEZO channel homologs are found throughout eukaryotic genomes^3,4,10^ but have been little studied outside animals. AtPIEZO1, a PIEZO homolog in the genome of the model flowering plant *Arabidopsis thaliana* is required to control the systemic spread of viruses^4^, but the mechanism is not known. How PIEZO channels might function in the green lineage (here defined to include green algae and land plants) is a particularly compelling question given the biomechanics of the plant cell: a large intracellular vacuole that presses the cytoplasm and plasma membrane against a sturdy yet flexible cell wall with up to a thousand-fold higher osmotic pressure than animal cells. Here, we report the first study of PIEZO channel function in the model plant *Physcomitrium* (formerly *Physcomitrella*) *patens*. This moss species has emerged as an exciting model system due to its phylogenetic position as an early land plant; its utility for studying development, tip-growth, water stress responses, and ease of imaging and genetic manipulations^11^.

## Phylogenetic analysis of plant and animal PIEZO homologs

To generate an updated phylogenetic tree of eukaryotic PIEZO homologs, protein sequences were selected from public databases based on multiple criteria: homology to mouse mPiezo1 and mPiezo2, large size (> 1800 amino acids), > 11 transmembrane helices, and to represent the diversity of eukaryotic lineages (Supplementary Table 1). Both Maximum Likelihood (Fig. 1a, Supplementary Fig. 1) and Neighbor-Joining (Supplementary Fig. 3) trees based on the conserved C-terminal domain (Extended Data Fig. 1b) show that modern PIEZO proteins descended from a single ancestor and that plant and animal homologs diverged very early, providing evolutionary space for specialization and diversification of function. In addition, they support a model of repeated duplication and loss events of PIEZO homologs in multiple eukaryotic lineages. Maximum likelihood trees based on full protein sequences further support duplications of the ancestral *PIEZO* gene in one genus of green algae; in mosses; in basal eudicots; and in core eudicots (Extended Data Fig. 1a, Supplementary Fig. 2). In each case, we named the subfamilies PIEZO1 and PIEZO2. In core eudicots, this ancestral duplication was followed by at least seven subsequent independent PIEZO1 duplication events. At the same time, Eudicot PIEZO2 was lost in at least 11 lineages. As a result, extant core eudicots have between 1 and 3 *PIEZO* homologs.

**Figure 1.**
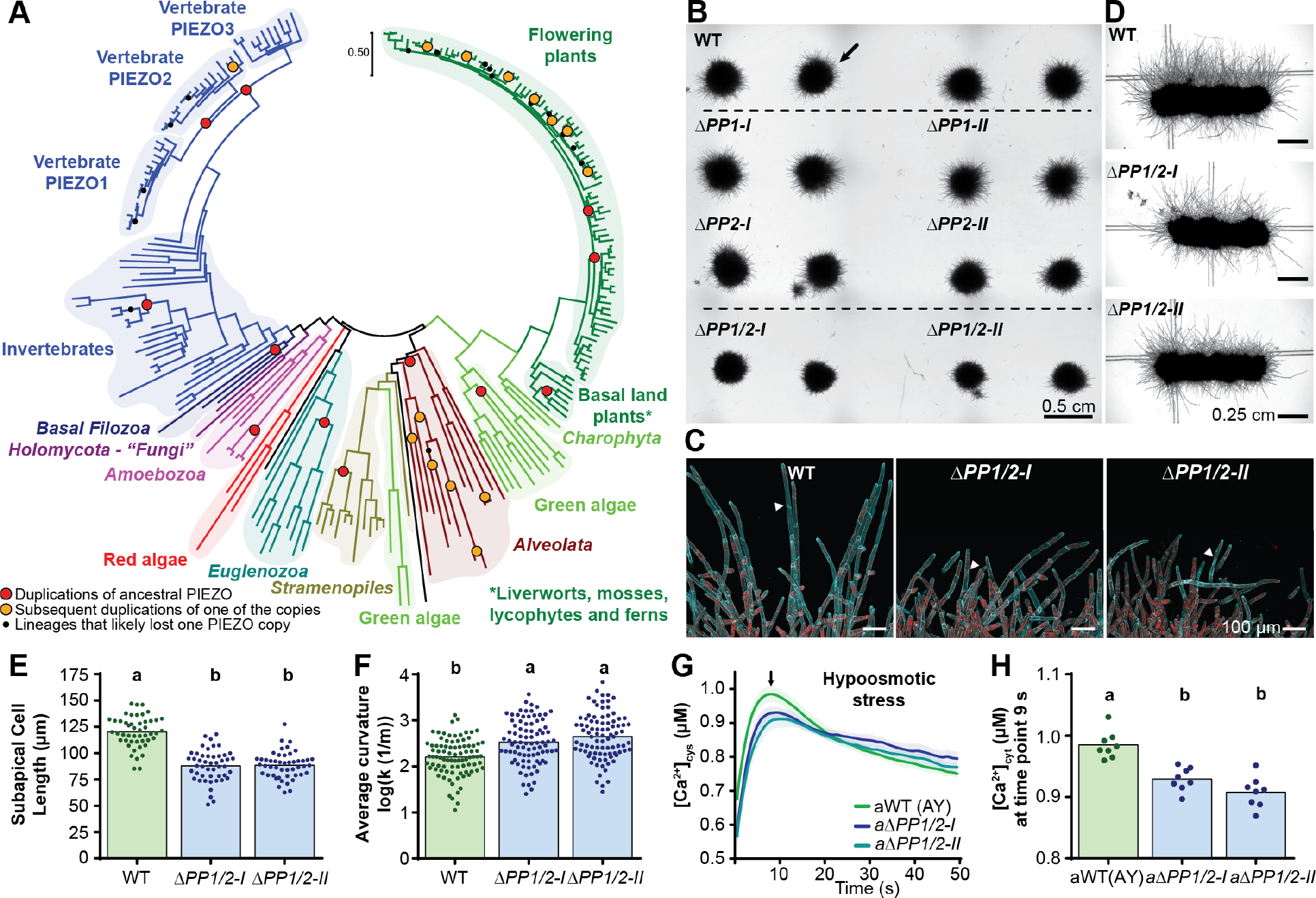
Moss PIEZO homologs are required for normal cell growth and cytosolic calcium transients. **a.** Maximum likelihood phylogenetic tree of 235 PIEZO homologs. **b.** Moss from fragmented protonema, grown for 7 days on cellophaned media. Black arrow, spreading caulonema filaments. **c.** Deconvolved Z-maximum intensity projections of cells at the plant edge. Cyan, calcofluor-white; red, chlorophyll autofluorescence. Arrowheads, oblique cross walls. **d.** Plants grown in the dark. **e.** Subapical caulonemal cell length. **f.** Curvature of caulonemal filaments. **g.** Average [Ca^2+^]_cyt_±SD (shaded area) in response to hypoosmotic shock. **h.** Quantification of [Ca^2+^]_cyt_ 9 s after shock in g. Statistics, one-way ANOVA with post-hoc Tukey test, p<0.05.

## Moss PIEZO homologs are redundantly required for caulonemal tip growth

We identified two PIEZO homologs encoded in the genome of *P. patens, Pp*PIEZO1 (encoded by Pp3c9_13300V3.1) and *Pp*PIEZO2 (encoded by Pp3c3_17170V3.1). They share 57% full-length protein sequence identity to each other, 20% identity to both mouse mPiezo1 and mPiezo2 and 41% to Arabidopsis PIEZO1. Single (*ΔPP1-I, ΔPP1-II, ΔPP2-I* and *ΔPP2-II;* for ***P**hyscomitrium* **P**IEZO) and double (*ΔPP1/2-I* and *ΔPP1/2-II*) mutant lines were generated using CRISPR/Cas9 gene editing (Extended Data Fig. 1c, Supplementary Table 2). When cultured on standard media with cellophane, *ΔPP1* and *ΔPP2* single mutant plants were similar in size to the WT, but double *ΔPP1/2* mutants were significantly smaller (Extended Data Fig. 1d). The most striking difference was in the filaments that spread away from the edge of the main plant, which were much shorter and less abundant (Fig. 1b, black arrow). Staining with a cell wall dye (Calcofluor-white) showed that those filaments were mostly comprised of caulonemal cells (Fig. 1c, note the oblique cross walls, white arrowheads). Caulonemal cells are one of the two cell types that form moss juvenile (protonema) tissue and are tasked with plant spreading^12^. In the dark, where only caulonemal cells grow, *ΔPP1/2* mutants exhibited fewer and shorter filaments in comparison to WT (Fig. 1d). *ΔPP1/2* caulonemal cells grew 30% more slowly than WT (Extended Data Fig. 1e, Supplementary Fig. 4). Furthermore, *ΔPP1/2* subapical caulonemal cells were approximately 25% shorter than WT (Fig. 1e) with the same cell width (Extended Data Fig. 1f). *ΔPP1/2* filaments were also significantly curvier (Fig. 1f). A third independently generated *ΔPP1/2* mutant line (Supplementary Table 2) showed similar growth defects (Extended Data Fig. 1g, h).

## *Pp*PIEZO1 and *Pp*PIEZO2 contribute to cytosolic calcium transients induced by hypoosmotic shock

Animal PIEZO channels allow for the influx of the cytosolic calcium ([Ca^2+^]_cyt_) that is observed in response to hypo-osmotic shock and membrane stretch in a range of cultured cells^13^. To determine if *Pp*PIEZO1 and *Pp*PIEZO2 have a similar function, we generated double mutants in moss lines expressing a cytosolic version of the calcium sensor aequorin-YFP (AY)^14^, *aΔPP1/2-I* and *aΔPP1/2-II* (Supplementary Table 2) and tested their response to hypo-osmotic shock (Extended Data Fig. 2a). WT plants expressing AY (*aWT*) showed a dramatic increase in cytosolic calcium concentration in response to hypoosmotic, but not iso-osmotic, treatment (Fig. 1g, Extended Data Fig. 2b-d). These [Ca^2+^]_cyt_ responses were dampened in *aΔPP1/2-I* and *aΔPP1/2-II* cells. At its maximum, 9 s after the shock, cytosolic calcium concentrations were significantly lower (~7.5%) in *aΔPP1/2-I* and *aΔPP1/2-II* lines than in *aWT* (Fig. 1h), indicating that *Pp*PIEZO double mutants released less calcium into their cytoplasm in response to hypo-osmotic shock compared with *aWT*.

## *Pp*PIEZO1 and *Pp*PIEZO2 localize to the vacuolar membrane

We expected that *P. patens* PEIZO homologs would localize to the plasma membrane, like their animal counterparts^3,16^. However, *P. patens* protoplasts transiently expressing codon-optimized o*Pp*PIEZO1 or o*Pp*PIEZO2 tagged with monomeric enhanced GFP (hereafter referred to as mGFP) from the maize *UBQ* promoter exhibited mGFP signal between the cytoplasm (marked with cytoplasmic mCherry) and the large central vacuole (void of signal) (Fig. 2a, left and middle panels, Extended Data Fig. 3a), closely resembling the vacuolar marker Vam3-mGFP (Fig. 2a, right panel). To validate this result, we used homologous recombination to add the coding sequence of *mGFP* to the 3’ ends of the *PpPIEZO1* and *PpPIEZO2* native loci (Extended Data Fig. 1c) to make *gPpPIEZO1-mGFP* and *gPpPIEZO2-mGFP* knock-in lines (Supplementary Table 2). Immunoblots indicated that full-length fusion proteins were present in all lines (Fig. 2b), and they grew normally (Extended Data Fig. 3b). gPpPIEZO1/2-mGFP localized to the vacuolar membrane, in subapical and apical caulonemal cells (Fig. 2c, Extended Data Fig. 3c) and chloronemal cells (Supplementary Fig. 5). Chloronemal cells are the second cell type of protonemal tissue and the main site of photosynthesis^12^. Finally, stable, constitutive expression of mGFP-fused, codon-optimized PIEZO homologs from moss (*oPpPIEZO1* and *oPpPIEZO2*) and from the dicot *Arabidopsis thaliana* (*oAtPIEZO1*) produced a localization pattern consistent with vacuolar targeting (Fig. 2d, Extended Data Fig. 3d). Multiple cellular compartments contribute to hypo-osmotic shock-induced [Ca^2+^]_cyt_ transients^15^, and the vacuole is likely to contribute but not dominate cytoplasmic calcium transients^19,20^. Thus, similar to previously characterized vacuolar MS cation channels from plants (TPK^17^) and yeast (TRPY1^18^), PpPIEZO1/2 channels likely release Ca^2+^ from vacuolar stores in response to mechanical stimuli.

**Figure 2.**
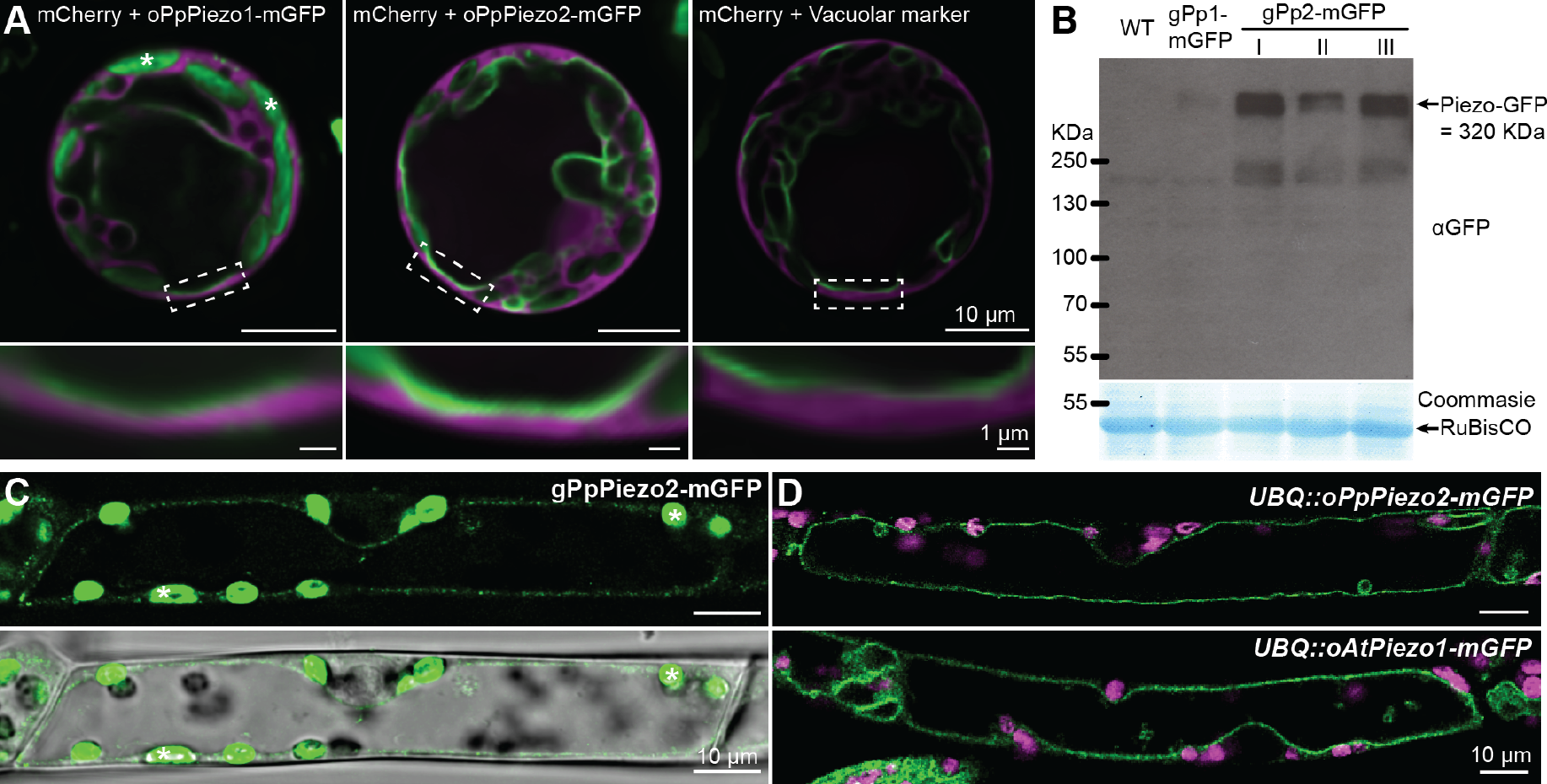
PpPIEZO proteins localize to the vacuolar membrane. **a.** Deconvolved single focal plane confocal image of WT protoplasts transiently co-expressing cytosolic mCherry (magenta) and *UBQ::oPpPIEZO1-mGFP, UBQ::oPpPIEZO2-mGFP*, or vacuolar marker *Vam3-mGFP* (green). Dashed boxes are areas magnified in the bottom panels. **b.** Immunoblot with total protein extracts from WT and *gPpPIEZO-mGFP* knock-in lines. **c.** Top, deconvolved single focal plane confocal image of a subapical caulonemal cell. Bottom, brightfield overlay. Green, mGFP; asterisks, chlorophyll autofluorescence. **d.** Deconvolved single focal plane confocal image of subapical caulonemal cells. Green, mGFP; magenta, chlorophyll.

## *PpPIEZO1/2* are required for normal vacuolar morphology

Vacuoles occupy most of the volume of many plant cells, store water, ions, and metabolites and are an essential part of turgor pressure, protein degradation, and cytosolic pH maintenance^21^. Vacuole morphology changes dynamically during development, growth, guard cell opening, and in response to stresses^22^. Given the unexpected localization of moss PIEZO homologs, we investigated vacuolar morphology in caulonemal cells in WT and mutant lines using brightfield microscopy (Fig. 3a, b), staining with 2’,7’-Bis-(2-Carboxyethyl)-5-(and-6)-Carboxyfluorescein, Acetoxymethyl Ester (BCECF), which accumulates in the vacuolar lumen (Fig. 3c) and staining with MDY-64, which, among other compartments, also stains the vacuolar membrane (Fig. 3d). As described previously^23^, we observed that in the apical cells of WT moss caulonema, the vacuole changed dynamically with the age of the cell. In younger and shorter WT cells the vacuole was in a highly tubulated or fragmented form (Extended Data Fig. 4a). As the apical cell elongated, the vacuole in the region behind the nucleus combined into one large central vacuole, while the region in the apical portion of the cell remained tubule-like in 90% of WT cells (top images in Fig 3a-d). In contrast, we found that 50-60% of the apical cells in single *ΔPP2-I* and *ΔPP2-II* mutants and >95% of the cells from double *ΔPP1/2-I* and *ΔPp1/2-III* mutants contained only large, expanded vacuoles in the apical region (Fig. 3a-d, Extended Data Fig. 4b). Sometimes we observed intravacuolar bubble-like structures (Extended Data Fig. 4b). A third *ΔPP1/2-III* mutant line had the same vacuolar phenotype (Extended Data Fig. 4c, d). This vacuolar phenotype was independent of growth rate (Extended Data Fig. 4e). Other cell types like subapical caulonemal (Extended Data Fig. 4f) and all chloronemal cells (Supplementary Fig. 5) contain only large and expanded vacuoles. Codon-optimized *PpPIEZO1, PpPIEZO2* and *AtPIEZO1*, fused to *mGFP* and overexpressed from the *UBQ* promoter (Supplementary table 2) were able to rescue the vacuole phenotype of the *ΔPP1/2-II* mutant (Fig. 3e; Extended Data Fig. 5a, b), suggesting a functional conservation among plant PIEZO homologs. Finally, the vacuolar morphology of apical caulonemal cells from *gPpPIEZO1-mGFP* and *gPpPIEZO2-mGFP* lines was indistinguishable from the WT (Extended Data Fig. 5c, d), indicating, together with the complementation experiments (Fig. 3e, Extended Data Fig. 5a, b), that GFP tagging did not alter protein function.

**Figure 3.**
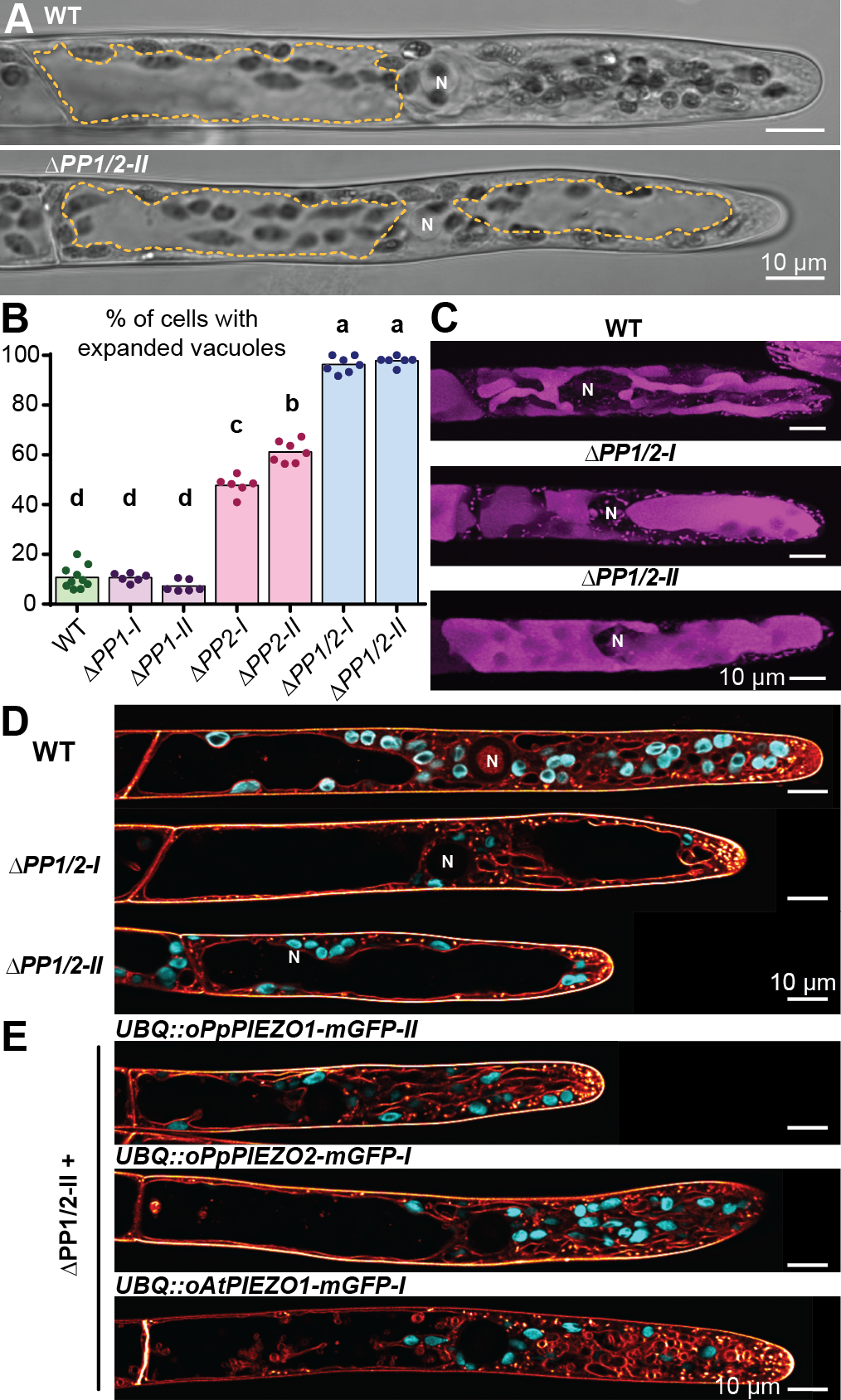
*PpPIEZO* double mutants have altered vacuolar morphology. **a.** Brightfield images of WT and *ΔPP1/2-II* apical caulonemal cells. Yellow lines, outline of expanded vacuoles. **b.** Percentage of cells with expanded nacu - oles in the tip region. Statistics as in Fig. 1. **c.** Cells stained with BCECF (Z-maximum intensity projection). **d.** Confocal single plane images of cells stained with MDY64 (orange). Cyan, chlorophyll autofluorescence. **e.** Confocal single plane images of *ΔPP1/2-II* cells expressing indicated constructs. Staining same as in **e.** All cells depicted are apical caulonemal cells. Fluorescence images were subjected to deconvolution.N, nucleus.

## Gain-of-function lesions in *Pp*PIEZO2 lead to complex intravacuolar membrane structures

Multiple gain-of-function lesions in the conserved C-terminal domains of animal PIEZOs are associated with disease^2^ (Fig. 4a, b). We introduced three such lesions, R2508K, R2508H, and E2548del, into the *PpPIEZO2* locus using CRISPR/Cas9 and oligodeoxynucleotide-assisted homology directed repair^24^ generating multiple independent lines for each lesion (Supplementary Table 2). Plants harboring the *Pp*PIEZO2-R2508K and -R2508H lesions, did not have any growth defects (Extended Data Fig. 6a, b), however, the vacuoles in the tip region of 60-70% of their caulonemal cells contained intravacuolar structures that were rarely observed in WT cells (Fig. 4c,-g, Extended Data Fig. 6c). Even among *PpPIEZO2-R2508K* and *-R2508H* mutant cells with tubule-like vacuolar morphology, 40-60% had internal bubble-like structures, compared to 20% of the WT (Fig. 4e, arrows). In mutant caulonemal cells with expanded vacuoles, we observed complex internal membrane structures (17-32% of total cells; Fig. 4f, arrows) and membrane lamination (5% and 15% of total cells in R2508K and R2508H mutants, respectively; Fig. 4g, number sign). In some cases, cytoplasm and chloroplasts were observed inside vacuolar membranes (Fig. 4f, arrow heads). Lamination was never observed in the WT. *Pp*PIEZO2 E2548del did not show these effects, instead producing a loss-of-function phenotype similar to *ΔPP2* mutants (Extended Data Fig. 7).

**Figure 4.**
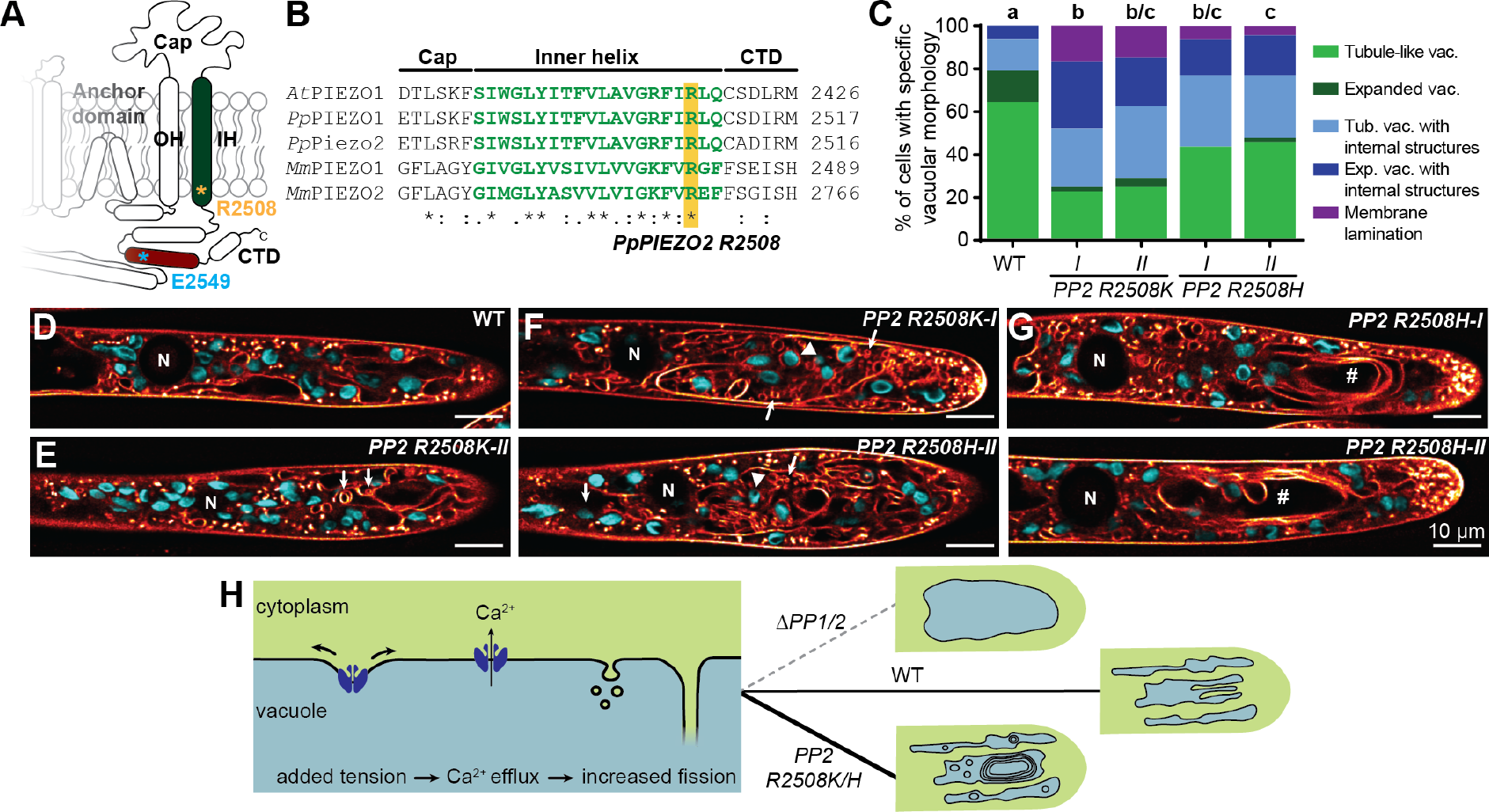
Vacuoles of *PpPIEZO2* gain-of-function mutants have complex internal membrane structures. **a.** Diagram of PIEZO pore module. **b.** Alignment of inner helix protein sequences **(*At, Arabidopsis; Pp***, moss; *Mm*, mouse). **c.** Quantitation of the vacuolar phenotypes shown in **d-g** in the indicated lines. Statistics: Fishers Exact Test. p<0.05 **d.** Tubule-like vacuoles **e.** Tubule-like vacuoles with internal structures (arrows). **f.** Expanded vacuoles with internal structures (arrows). **g.** Expanded vacuoles with membrane lamination (number signs). **d-g.** Deconvolved single focal plane confocal images of apical caulonemal cells stained with MDY64 (orange). Cyan, chlorophyll auto-fluorescence; N, nucleus; arrowheads, internalized chloroplasts. **h.** A model for PpPIEZO1/2 function in moss caulonemal cells.

## Summary

Taken together, these data show that moss PIEZO homologs regulate vacuolar morphology in apical caulonemal cells. Vacuole remodeling is linked to mechanosensing in plant cells^25^, and vacuolar invagination and fragmentation occur in response to high [Ca^2+^]_cyt_ in yeast^26^. We speculate that *Pp*PIEZOs release Ca^2+^ into the cytoplasm from vacuolar stores in response to the physical state of the vacuolar membrane, which in turn leads to an increase in the surface volume of vacuolar membranes (Fig. 4h). During WT caulonemal tip-growth, a tubule-like vacuole in the tip may allow these exploratory cells to grow more efficiently, and to better adapt to local changes in their environment. The work presented here also demonstrates that moss PIEZO homologs have diverged both in cellular localization and function from their animal counterparts and suggests that they have been co-opted to sense the mechanical status of the plant vacuolar membrane. Like animal PIEZO channels, plant PIEZOs still release calcium into the cytosol, just from vacuolar rather than extracellular stores. Perhaps this relocation to the vacuole reflects a higher freedom of movement of vacuolar membranes compared with the plasma membrane in plant cells, making them a preferred location to sense and respond to mechanical changes within the cell. Future work may reveal a multiplicity of ways in which PIEZO homologs have been adapted to the specific needs of the green lineage.

## Acknowledgements

This work was funded by HHMI-Simons Faculty Scholar Grant 55108530 to E. S. H. and NSF MCB-1330171 to M. B. We thank Shu-zon Wu for generating the Vam3-GFP vacuolar marker, and Seyed Ali Reza Mousavi and Ardem Patapoutian for codon-optimized *AtPIEZO1* and helpful discussions.

## Author contributions

I.R, R.R., E.W, and C.J.B. designed and performed experiments and analyzed data. I.R., R.R., M.B., and E.S.H analyzed data and wrote the paper.

---

**Extended Data Figure 1.**
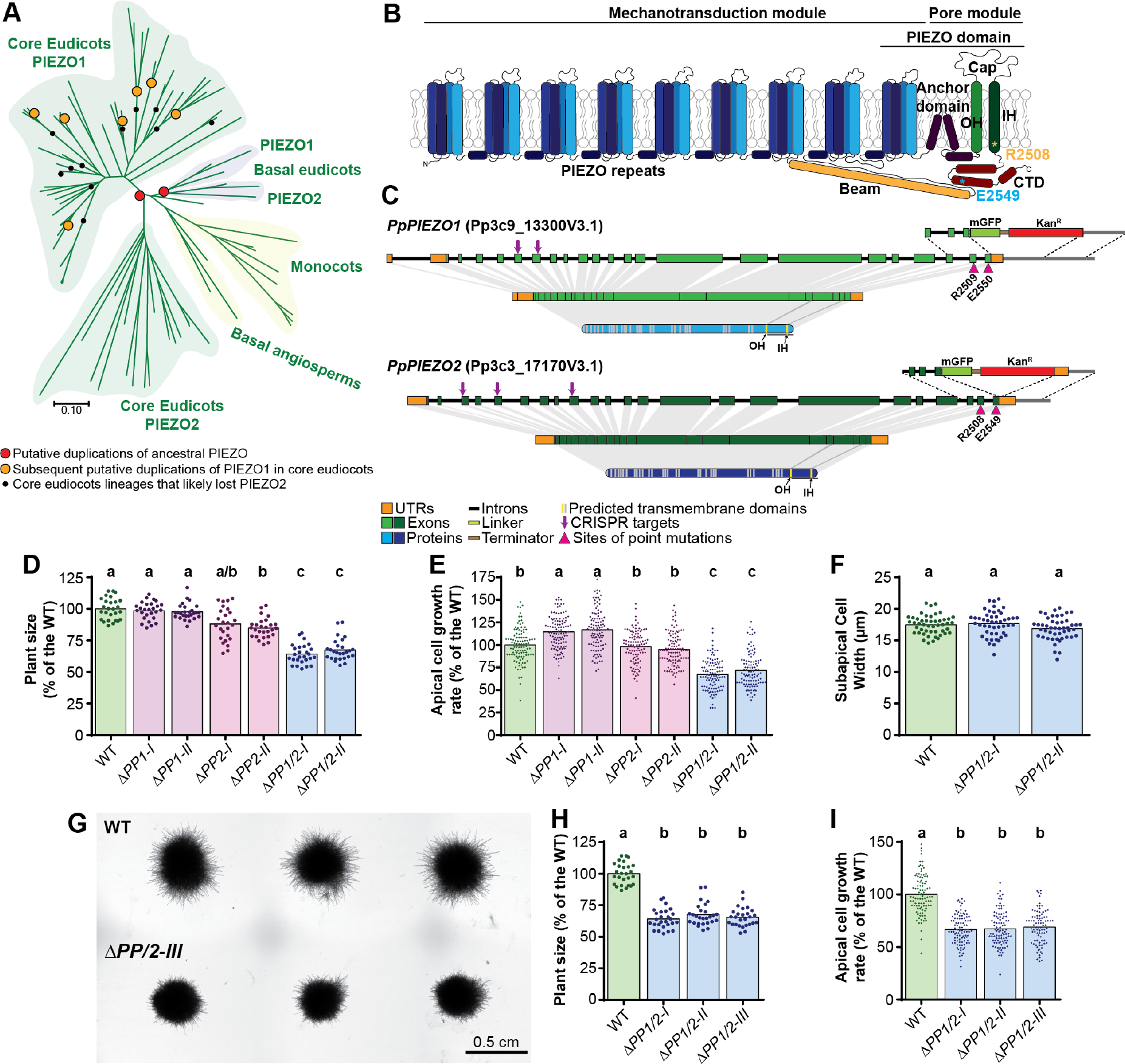
*Pp*PIEZO homologs in are required for normal growth in moss. **a.** Maximum Likelihood Unrooted Phylogenetic Tree of 83 PIEZO homologs from angiosperms. Full-length PIEZO protein sequences were used to generate the tree, which is drawn to scale with branch lengths corresponding to evolutionary distances expressed as amino acid substitutions per site. For details see Supplementary Fig. 2b. Accession numbers and details about PIEZO homolog sequences used are listed in Supplementary Table 1. **b.** Model of a typical PIEZO monomer based on CryoEM structures of mouse Piezo1 and Piezo2 channels. Asterisks mark the positions of residues mutated in *PpPIEZO2* gain-of-function alleles. OH, Outer helix; IH, inner pore-lining helix; CTD, C-terminal domain **c.** Schematics of the two moss *PIEZO* loci (drawn to scale) and the corresponding mRNA and protein products. The transmembrane domains drawn were predicted by TMHMM. Arrows mark the position of the guide RNA targets used to generate *PpPIEZO* knock-out lines. Also indicated are the integration cassettes with homology arms, monomeric GFP and Kanamycin resistance genes used to generate moss knock-in lines. Arrowheads mark the positions of point mutations generated in *PpPIEZO2*. **d.** Size (area) of moss started from fragmented protonema after 7 days of growth on cellophaned BCDAT. **e.** Average tip growth rate of caulonemal cells at the plant edge. **f.** Width of subapical caulonemal cells. **g.** WT and *ΔPP1/2-III* (a third independently generated *PpPIEZO1/2* double mutant) after 7 days of growth on cellophaned BCDAT media. **h.** Size (area) of moss started from fragmented protonema after 7 days of growth on cellophaned BCDAT plates. Data for WT, *ΔPP1/2-1* and *-II* is repeated from **d. i.** Average tip growth rate of caulonemal cells on the plant edge. Some data for WT, *ΔPP1/2-1* and *-II* are repeated from **e.** In **d, e, h**, and **i**, graphs are a combination of three replicates, each normalized to their respective WT. Statistics: Kruskal-Wallis test with Dunn’s multiple comparisons test. In **f**, graph is combination of three replicates. Statistics: one-way ANOVA with post-hoc Tukey test. p<0.05. Average values, standard deviations and N are given in Supplementary Table 3.

**Extended Data Figure 2.**
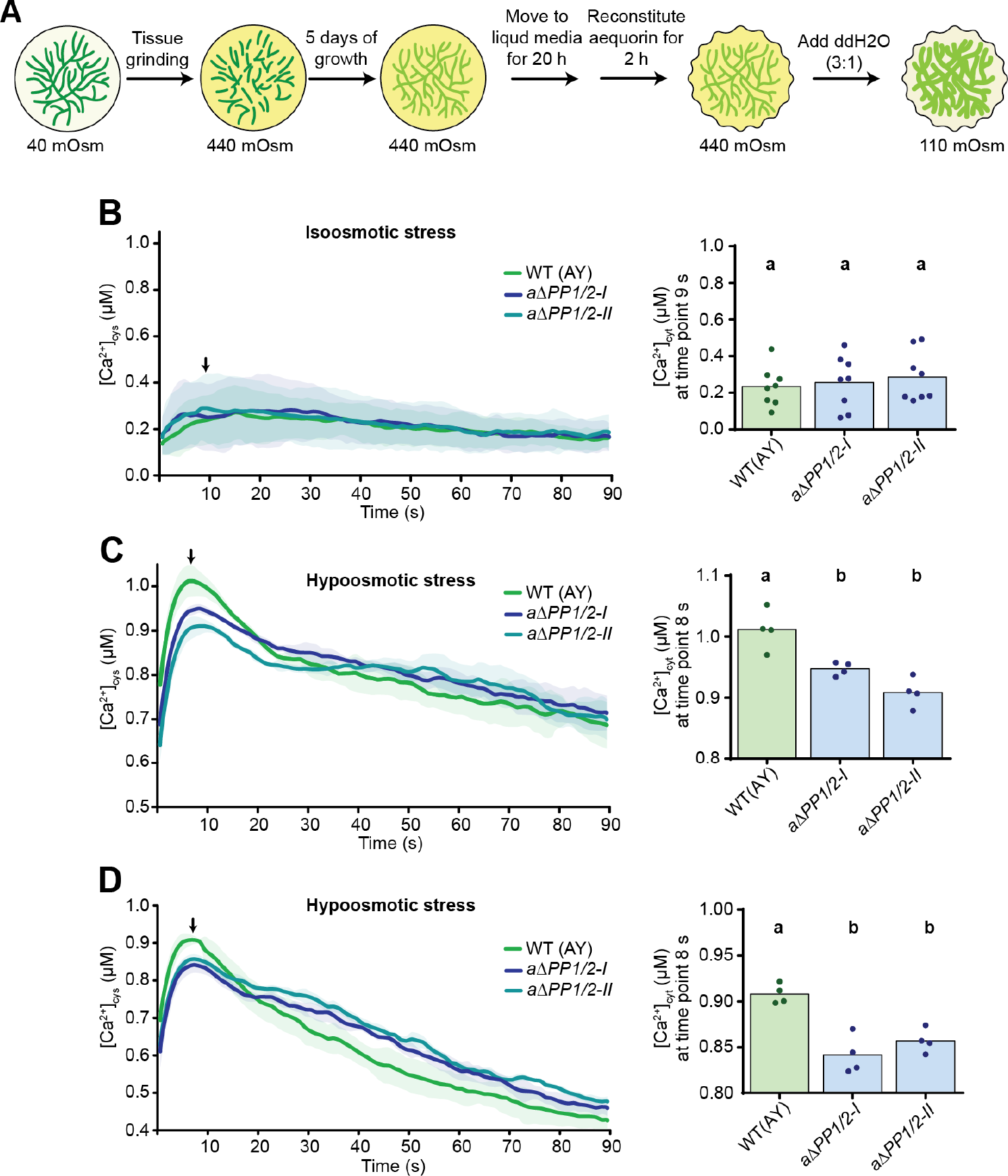
*Pp*PIEZO1 and 2 are necessary for full cytosolic calcium response to hypoosmotic shock. **a.** Schematic of the experimental setup. Numbers correspond to the calculated osmotic potential of moss media at each step. **b.** Change of cytosolic calcium concentration in response to isosmotic media. **c, d.** Two additional replicates of cytosolic calcium concentration in response to hypoosmotic shock. Recording of bioluminescence was started immediately after the addition of water or isosmotic media. Each line represents an average of 8 replicates for isosmotic treatment and 4 replicates for both hypoosmotic shock. Shaded areas, standard deviation; black arrows mark the peak of the response. Cytosolic calcium concentrations for this time point are depicted on the right-hand side of each graph. Statistics, one-way ANOVA with the post-hoc Tukey test. p<0.05. Average values, standard deviations and N are given in Supplementary Table 3.

**Extended Data Figure 3.**
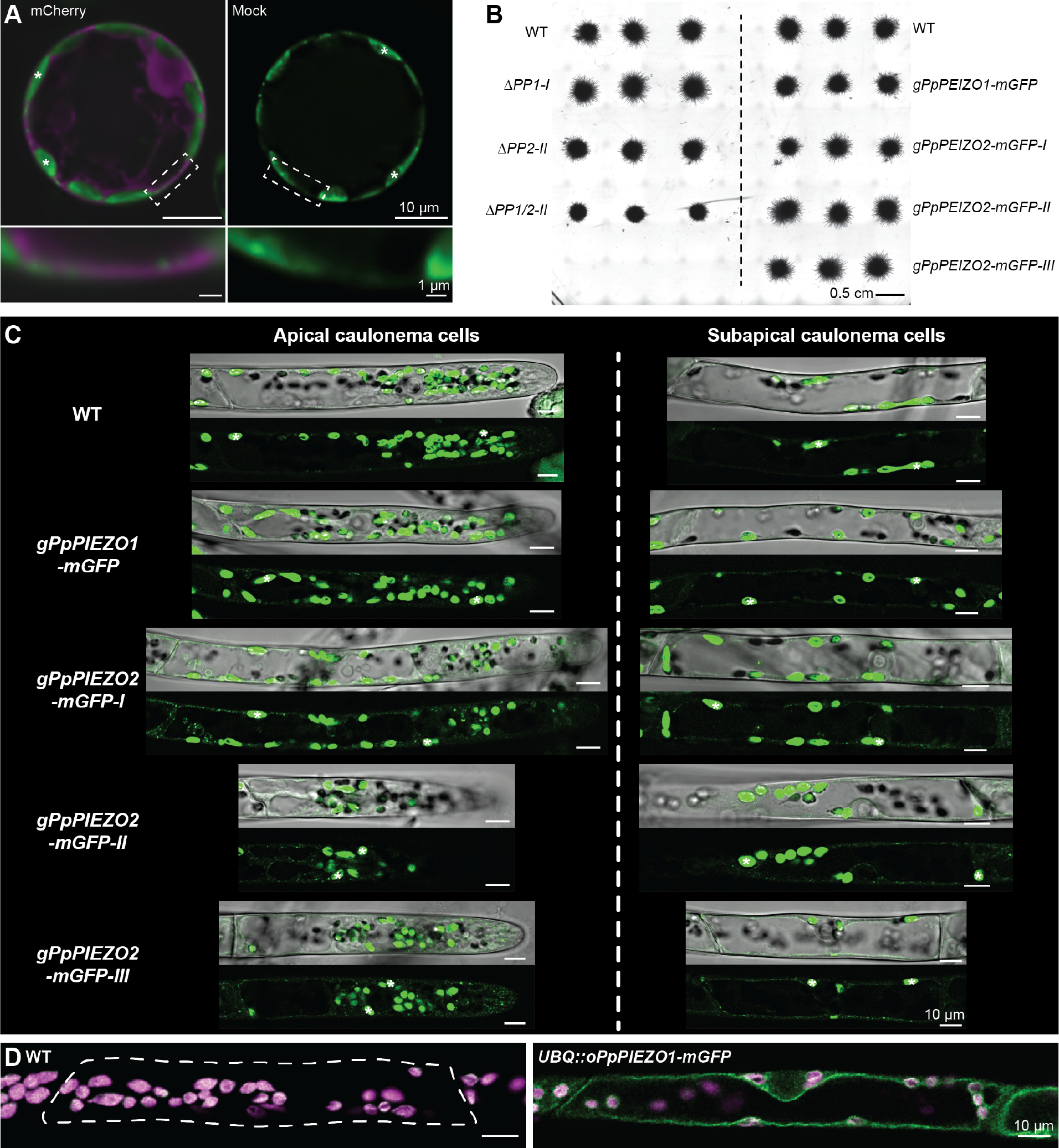
*Pp*PIEZO1 and *Pp*PIEZO2 localize to the vacuolar membrane regardless of expression level. **a.** Deconvolved confocal images of WT protoplasts 4 days after transformation with water or with a construct encoding cytosolic mCherry (magenta). Bottom panels are magnification of boxed areas in corresponding upper panel. **b.** Moss plants (started from fragmented protonema) of the indicated genotypes after 7 days of growth on cellophaned BCDAT media. **c.** Additional deconvolved single focal plane confocal images of *gPpPIEZO1-mGFP* and *gPpPIEZO2-mGFP* in apical and subapical caulonemal cells expressed from the native loci. Top panels, confocal image of the green channel; bottom panels, brightfield overlay. Asterisks, chlorophyll autofluorescence. **d.** Deconvolved single focal plane confocal image of subapical caulonemal cells from WT (left panel) and *UBQ::oPpPIEZO1-mGFP* moss (right panel). Green, mGFP; magenta, chlorophyll autofluorescence.

**Extended Data Figure 4.**
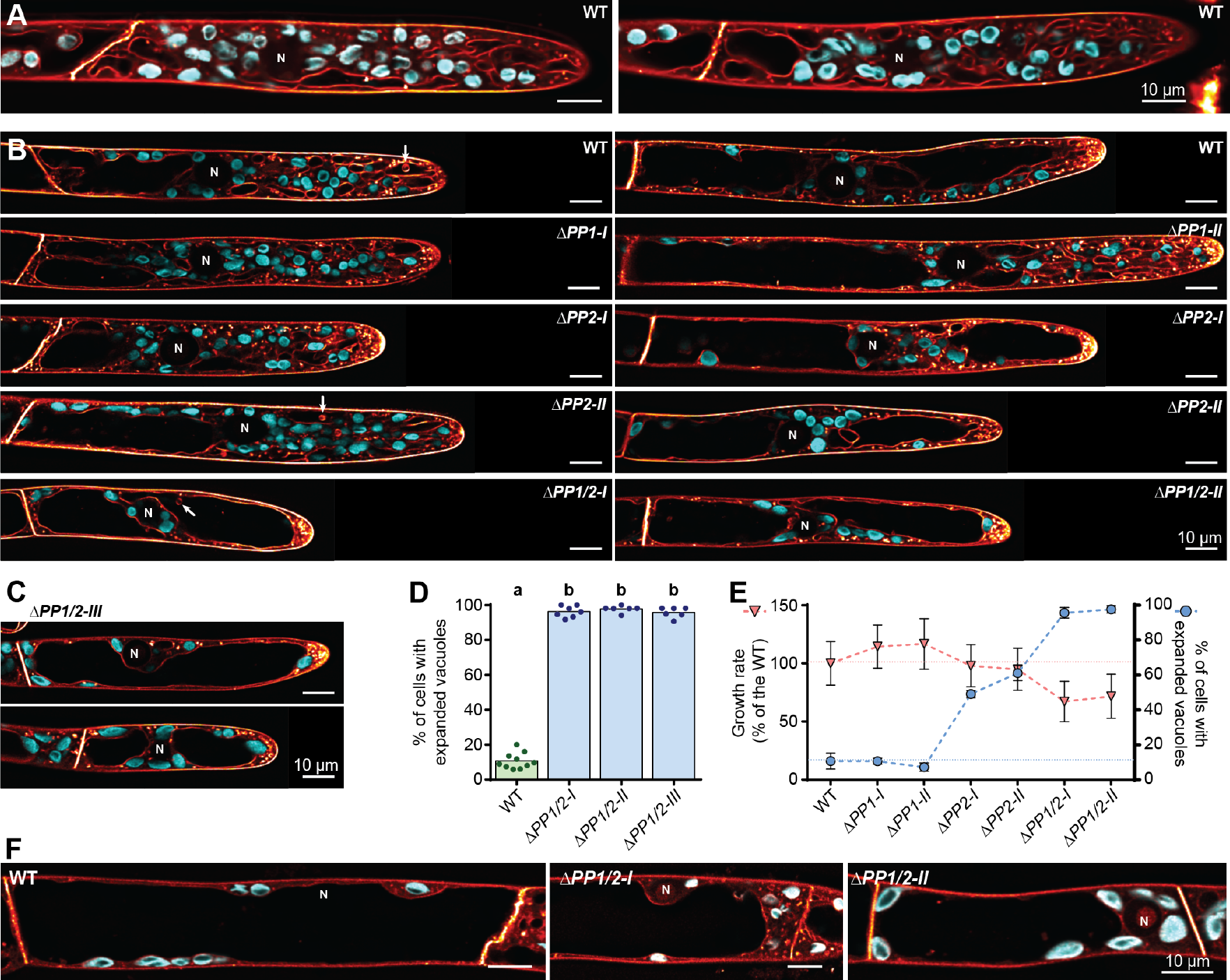
Change in the vacuolar morphology in piezo null mutants is not dependent on cell growth rate. **a.** Single slice deconvolved confocal images of vacuolar morphology in young, recently divided WT apical caulonemal cells stained with MDY64 (orange). Cyan, chlorophyll autofluorescence. Note tubule-like vacuolar morphology in the region between the cell base and the nucleus. **b.** Deconvolved single focal plane confocal images of vacuolar morphology in mature apical caulonemal cells from WT and *PpPIEZO* single and double mutants (same staining as in **a).** Note the expanded vacuoles in the region between cell base and nucleus. Arrows, intravacuolar membrane bubble-like structures. **c.** Examples of vacuolar morphology in *ΔPP1/2-III* moss. Same staining as in **a. d.** Percentage of cells with expanded vacuoles in the tip region for all three *ΔPP1/2* lines. Data for WT, *ΔPP1/2-1* and *-II* is repeated from Figure 3B. Statistics, one-way ANOVA with the post-hoc Tukey test. p<0.05. **e.** Average growth rate normalized to that of the WT (pink triangles, data from Extended Data Fig. 1e) and percentage of apical caulonemal cells with expanded vacuoles. for WT and PpPIEZO single and double mutant lines. Blue circles, data from Fig. 3b. Error bars are standard deviation, while dashed lines mark average values for the WT. Note that *ΔPP2* single mutants have growth rates similar to the WT, but ~50% of apical caulonemal cells have expanded vacuoles. **f.** Single slice deconvolved confocal images of vacuolar morphology in subapical caulonemal cells from WT and *ΔPP1/2* double mutants (same staining as in a). N, nucleus.

**Extended Data Figure 5.**
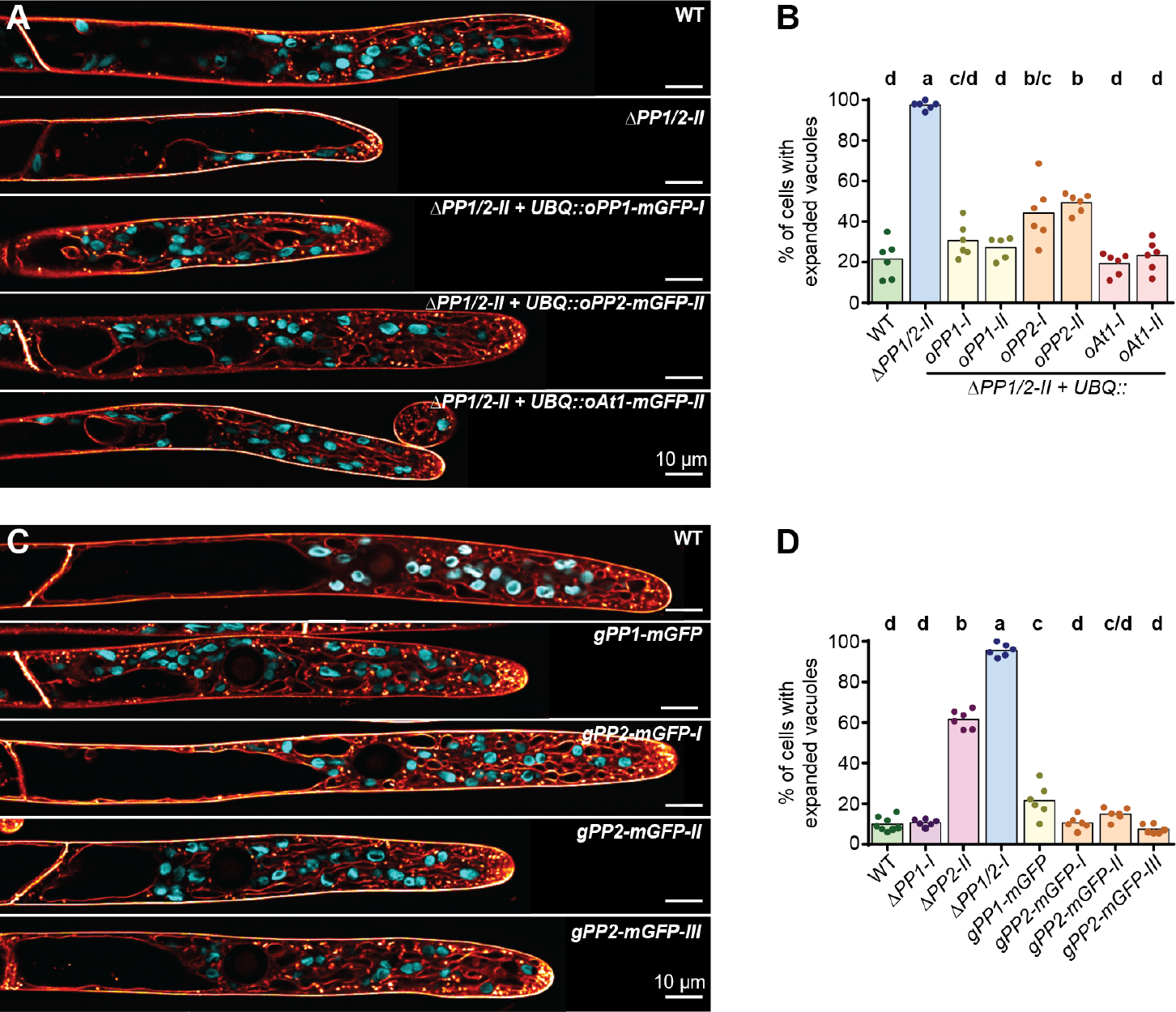
mGFP tagging does not alter the function of PpPIEZO1 and PpPIEZO2. **a, b.** Single slice deconvolved confocal images of vacuolar morphology in apical caulonemal cells, from indicated moss lines, stained with MDY64 (orange). Cyan, chlorophyll autofluorescence. **c, d.** Percentage of cells with expanded vacuoles in the tip region. Some data for WT, *ΔPP1-1, ΔPP2-II* and *ΔPP1/2-1*, in **d.** are repeated from Figure 3b. Statistics, one-way ANOVA with the post-hoc Tukey test. p<0.05. Average values, standard deviations and N are given in Supplementary Table 3.

**Extended Data Figure 6.**
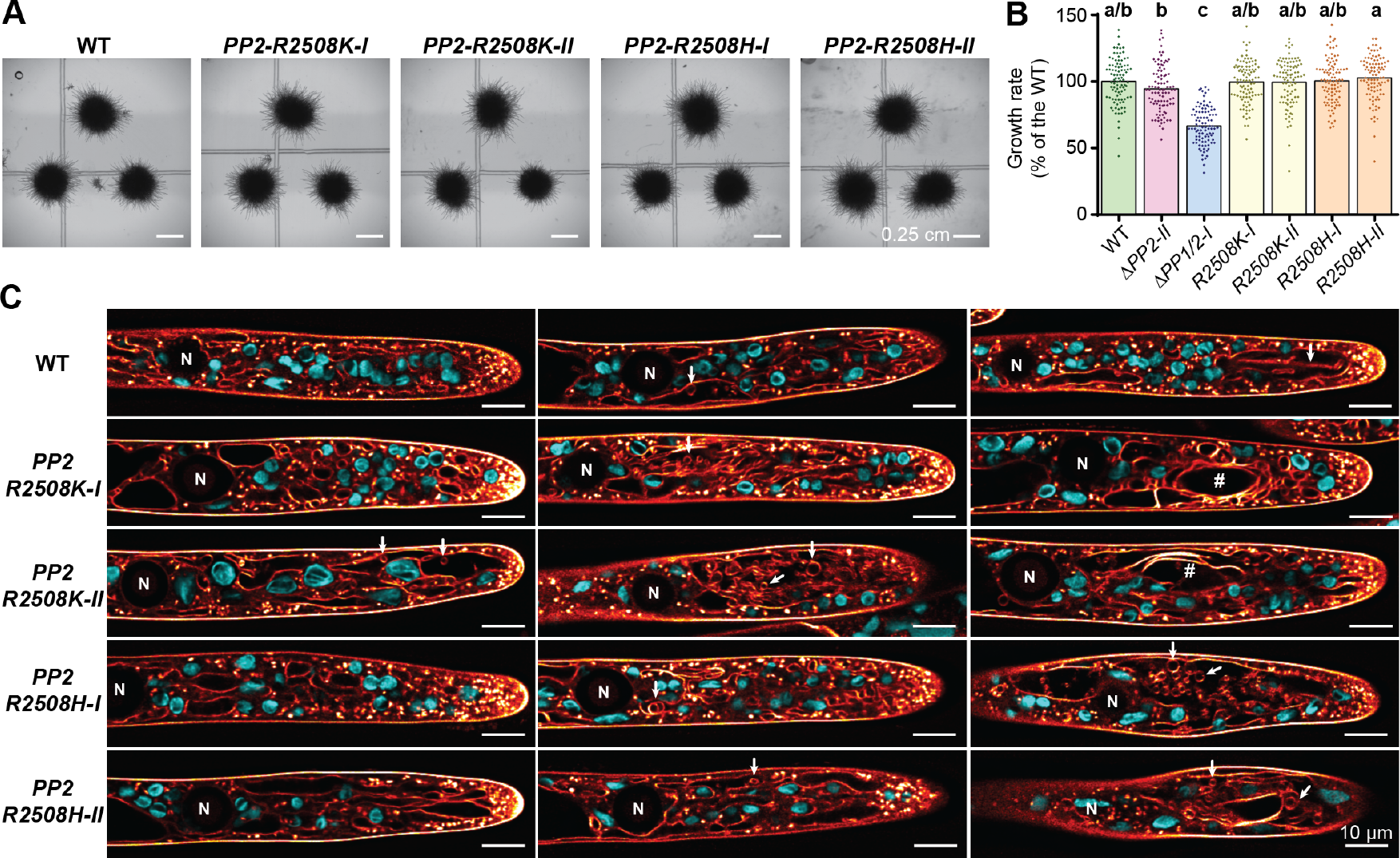
*PpPIEZO2 R2508K* and *R2508H* mutants have normal growth rate and altered vacuolar morphology. **a.** Moss plants (started from fragmented protonema) of the indicated genotypes after 6 days of growth on cellophaned BCDAT media. **b.** Average tip growth rate of caulonemal cells on the plant edge. Some data for *ΔPP2-II* and *ΔPP1/2-1* are repeated from Extended Data Fig. 1e. Graph is a combination of three replicates, each normalized to their respective WT. Statistics: Kruskal-Wallis test with Dunn’s multiple comparisons test. p<0.05. Average values, standard deviations and N are given in Supplementary Table 3. **c.** Single slice deconvolved confocal images of vacuolar morphology in apical caulonemal cells stained with MDY64 (orange). Cyan, chlorophyll autofluorescence). Examples of cells with different vacuolar morphologies are shown: tubule-like, expanded, tubule-like vacuoles with internal structures (arrows), expanded vacuoles with internal structure (arrows) and membrane lamination (number signs). Average values, standard deviations and N are given in Supplementary Table 3.

**Extended Data Figure 7.**
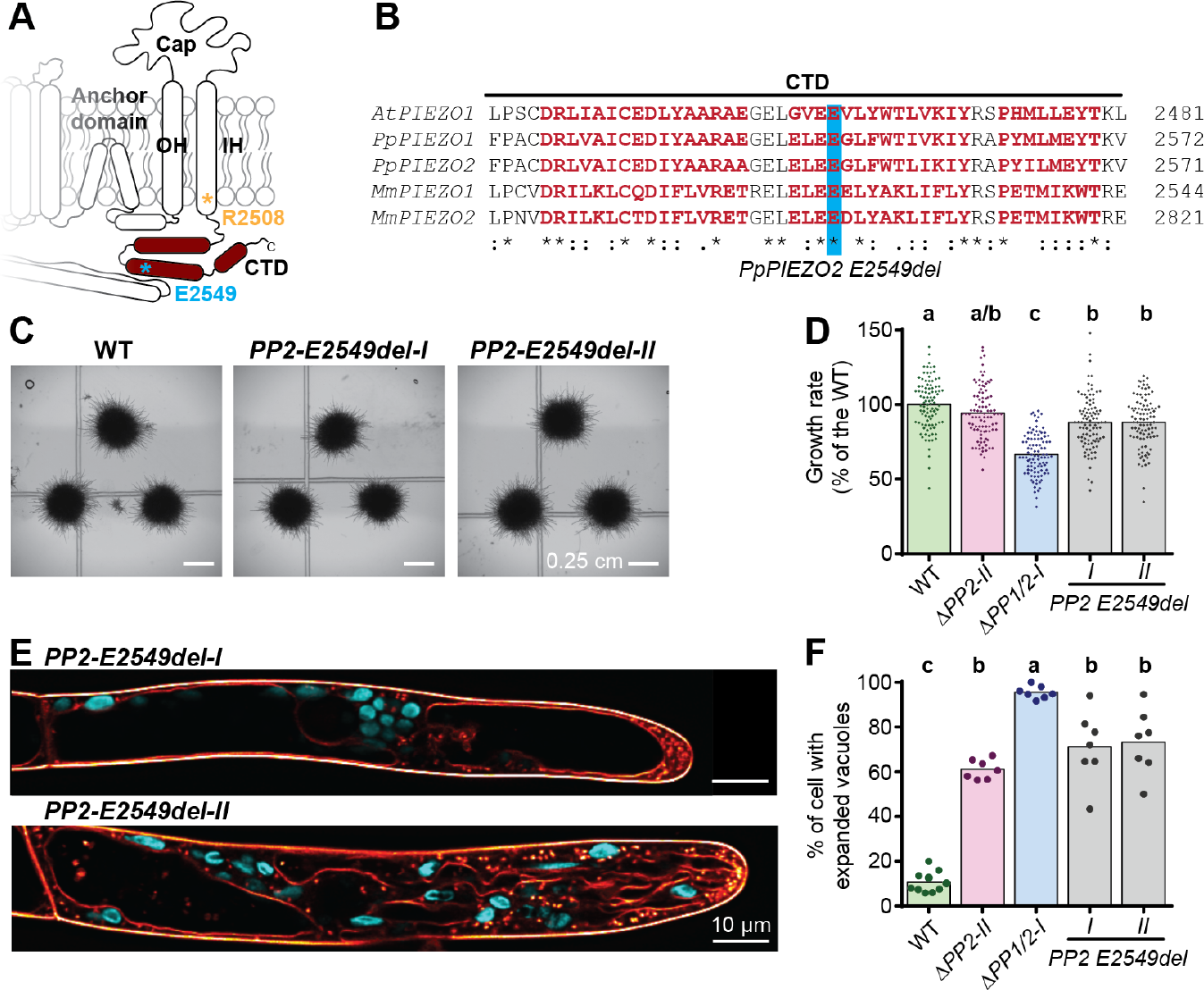
*PpPIEZO2 E2549del* produced loss-of-function phenotypes. **a.** Diagram of the PIEZO pore module. Asterisk, mutated residues. **b.** Alignment of C-terminal domain protein sequences from PIEZO homologs from *Arabidopsis thaliana* (*At*), *Physcomitrium patens* (*Pp*) and *Mus musculus* (*Mm*). **c.** Moss plants (started from fragmented protonema) of the indicated genotypes after 6 days of growth on cellophaned BCDAT media. **d.** Average tip growth rate of caulonemal cells on the plant edge. Some data for WT, *ΔPP2-II* and *ΔPP1/2-1* are repeated from Extended Data Fig. 1e. Graph is a combination of three replicates, each normalized to their respective WT. Statistics: Kruskal-Wallis test with Dunn’s multiple comparisons test. **e.** Examples of vacuolar morphology in apical caulonemal cells. MDY64, orange; cyan, chlorophyll autofluorescence. **f.** Percentage of cells with expanded vacuoles in the tip region. Data for *ΔPP2-II* and **ΔPP1/2-1** are repeated from Figure 3b. Statistics, one-way ANOVA with the post-hoc Tukey test. p<0.05. Average values, standard deviations and N are given in Supplementary Table 3.

## Methods

### Phylogenetic analysis

Full-length protein sequences of putative PIEZO homologs were identified by BLAST search of selected genomes using NCBI, Phytozome, and other databases. Sources for all protein sequences are given in Supplementary Table 1. Genomes for analysis were selected to represent as many eukaryotic lineages as possible. For those lineages where many genomes are available, only the genomes of representative species were analyzed. For flowering plants, we analyzed a larger number of genomes to help resolve the complex pattern of gene duplications and losses.

All homologs included in the analysis were identified by BLAST searches when either mouse mPiezo1 or mPiezo2 were used as query sequences. The only exceptions were homologs from *Mesotaenium endlicherianum*, which were identified by a direct search of predicted protein sequences using the conserved PFEW motif^1,2^. To further ensure that we included the best candidates, we utilized two well-known PIEZO protein characteristics: their large size (averaging around 2500 amino acids), and a very large number of predicted transmembrane domains with last two being separate from the others (Extended Data Fig. 1b). As such, all homologs included had at least 1800 amino acids and a large number of predicted transmembrane domains. The TMHMM Server v. 2.0^3^ was used to predict the transmembrane domains. The predicted sizes and number of transmembrane domains for each homolog are given in Supplementary Table 1. If multiple isoforms of PIEZO homologs were present, only one was used for analysis.

Phylogenetic analysis was performed with MEGA7^4^ software as described in^5^. For the tree depicted in Fig. 1a and Supplementary Figs, 1 and 3, 235 full-length protein sequences were aligned using MUSCLE^6^ with default settings. Parts of the alignments outside of the conserved PIEZO domain (T1973 to R2543 in mouse mPiezo1) were removed and the remaining PIEZO domain sequences were realigned with MUSCLE. That alignment, with average evolutionary divergence over all sequence pairs (p-distance model) of 0.636±0.010, was used to estimate the Maximum Likelihood tree in Fig. 1a and Supplementary Fig. 1 (LG+G+I model^7^; *+G* = 09913 with 5 categories; [+*I*] 0.30% of sites were allowed to be evolutionary invariable; only sites with 10% or fewer gaps in the alignment were included (502 sites); 500 bootstrap replications; log-likelihood = −119236.92) and the Neighbor-Joining tree^8^ in Supplementary Fig. 3 (with JTT matrix-based method^9^ and only sites with 10% or less gaps in the alignment = 502 sites in the final data set; 1000 bootstrap replications). For the tree in Extended Data Fig. 1a and Supplementary Fig. 2b, full-length sequences of 83 angiosperm (flowering plant) PIEZO homologs, with average evolutionary divergence (p-distance model) of 0.327±0.005, were aligned using MUSCLE and used to generate Maximum likelihood tree (JTT+G+I+F model^9^; *+G* = 09330 with 5 categories; [+*I*] 8.35% of sites were allowed to be evolutionary invariable; only sites with 10% or fewer gaps in the alignment were included (2307 sites); 500 bootstrap replications; log-likelihood - 135775.95). For the tree in Supplementary Fig. 2a, full-length sequences of 73 animal PIEZO homologs, with average evolutionary divergence (p-distance model) of 0.579±0.006, were aligned using MUSCLE and used to generate Maximum likelihood tree (LG+G+I^7^; *+G* = 0.8267 with 5 categories; [+*I*] 1.26% of sites were allowed to be evolutionary invariable; only sites with 10% or fewer gaps in the alignment were included (2104 sites); 500 bootstrap replications; log-likelihood −181869.63).

### Plant material and growth conditions

All plant lines used in this study have been generated in the wild-type *Physcomitrium* (formerly *Physcomitrella) patens* (Gransden strain) background. Moss was cultured under sterile conditions on BCDAT media (1.01 mM MgSO_4_, 1.84 mM KH_2_PO_4_, 10 mM KNO_3_, 5 mM Di-ammonium tartrate, 1 mM CaCl_2_, 45 μM FeSO_4_ and microelements (9.95 μM HBO_3_, 1.97 μM MnCl_2_, 0.232 μM AlK(SO_4_)_2_, 0.220 μM CuSO_4_, 0.235 μM KBr, 0.661 μM LiCl, 0.103 μM Na_2_MoO_4_, 0.231 μM CoCl_2_, 0.191 μM ZnSO_4_, 0.169 μM KI, 0.124 μM SnCl_2_, 0.248 μM NiCl_2_), pH = 6.5 (adjusted with KOH). For solid media, 0.7% agar (Sigma-Aldrich A9799) was added. Moss was cultured in a growth chamber with 80 μmol_photons_ m^−2^ s^−1^ of light under long-day conditions (16/8 h cycle with 24°C/21°C day and night, respectively). Moss was cultured as described in^10^. For maintenance of a weekly protonemal culture, 7-day-old tissue was ground with an Omni soft-tissue homogenizer and placed on solid BCDAT media overlaid with cellophane (325P cellulose 80 mm disc from A.A. Packaging Ltd).

### Protoplast isolation, regeneration, and transformation

Protoplasts were generated as previously described with some modification^11^. A 5 to 7-day-old protonemal tissue culture was scraped from solid plates and added to a 466 mM mannitol solution with 2% Driselase and Cellulase (21 unit/mL, Worthington CEL). The mixture was gently agitated for 1 h at room temperature (RT). Protoplasts were filtered through 100 μm stainless-steel mesh, spun down (200xg/ RT/ 5 min), and washed 3 times with 466 mM mannitol.

To regenerate plants, isolated protoplasts were spun down, resuspended in liquid plating media (1.03 MgSO_4_, 1.84 mM KH_2_PO_4_, 3.39 mM CaNO_3_, 45 μM FeSO_4_, 2.72 mM Di-ammonium tartrate, 10 mM CaCl_2_, 466 mM mannitol and microelements) and spread on cellophane covered PRMB plates (1.03 MgSO_4_, 1.84 mM KH_2_PO_4_, 3.39 mM CaNO_3_, 45 μM FeSO_4_, 2.72 mM Di-ammonium tartrate, 10 mM CaCl_2_, 330 mM mannitol, microelements and solidified with 0.8% agar). After 4 days of regeneration, cellophanes with young plants were moved to regular BCDAT plates.

Protoplast transformations were performed as previously described with some modifications^12^. Protoplasts were resuspended in 3M solution (500 mM mannitol, 15 mM MgSO_4_, 5.1 mM MES pH 5.6) to concentration of 2 million/mL. For each transformation, 300 μL of protoplasts was immediately mixed with plasmid DNA (~30 μg per plasmid, usually <10 μL) and 300 μL of filter-sterilized PEG solution (4 g of melted PEG 8000 mixed with 10 mL of 420 mM mannitol, 100 mM Ca(NO_3_)_2_ and 10 mM Tris pH 8.0 solution). The suspension was gently mixed by pipetting, incubated at RT for 10 min, heat-shocked for 3 min at 45°C, and incubated for another 10 min in a RT water bath. Subsequently, 10 mL of 466 mM mannitol solution was added to the transformation mixture, mixed and incubated for at least 30 min at RT. Finally, the protoplasts were spun down and resuspended in liquid plating media and either left for 4 days in that media for imaging (Fig. 2a, Extended Data Fig. 3a) or spread on top of cellophane-covered PRMB plates. After 4 days of protoplast regeneration on PRMB media, cellophanes with young plants were moved to BCDAT plates with appropriate antibiotics (15 μg/mL Hygromycin, 20 μg/mL G418 or 50 μg/mL Zeocin) for plant selection. For transient transformations, the selection was applied for 7 days, after which the plants were moved to BCDAT plates for subsequent genotyping and propagation. For stable transformations (genome integrations), plants were selected for 7 days, followed by a week of recovery on BCDAT plates and another week on antibiotic selection, after which plants were moved to BCDAT for subsequent genotyping.

### Cloning of constructs used in this study

#### pZeo-Cas9-PpPIEZO1

For expression of Cas9 and two guide RNAs (protospacers: 5’ – TAGGTGTAGCTCGGCCGTCA - 3’ and 5’ - CATCAGGTCAACTGCTGGGA - 3’ (Extended Data Fig. 1c), chosen with CRISPOR program^13^) targeting *PpPIEZO1* (Pp3c9_13300V3.1). Primers for guide RNA 1 (5’- CCATTAGGTGCTCGGCCGTCA -3’ and 5’-AAACTGACGGCCGAGCTACACCTA -3’) and guide RNA 2 (5’-CCATCATCAGGTCAACTGCTGGGA-3’ and 5’-AAACTCCCAGCAGTTGACCTGATG-3’) were annealed and ligated into pENTR-PpU6P-sgRNA-L1R5 and pENTR-PpU6P-sgRNA-L5L2 as described in^14^. Subsequently, both guide RNA expression constructs were combined into pZeo-Cas9-gate vector^14^, via LR+ recombination.

#### pZeo-Cas9-PpPIEZO2

For expression of Cas9 and two guide RNAs (protospacers: 5’ – CTCGTCCAAGGTCAGGACTA - 3’ and 5’ - CTGTCCAGCTGATAGCGGGA - 3’ (Extended Data Fig. 1c), chosen with CRISPOR program^13^) targeting *PpPIEZO2* (Pp3c3_17170V3.1). Primers for guide RNA 1 (5’-CCATCTCGTCCAAGGTCAGGACTA-3’ and 5’-AAACTAGTCCTGACCTTGGACGAG -3’) and guide RNA 2 (5’-CCATCTGTCCAGCTGATAGCGGGA-3’ and 5’-AAACTCCCGCTATCAGCTGGACAG-3’) were annealed and ligated into pENTR-PpU6P-sgRNA-L1R5 and pENTR-PpU6P-sgRNA-L5L2 as described^14^. Subsequently, both guide RNA expression constructs were inserted into the pZeo-Cas9-gate vector^14^, via LR+ recombination.

#### pMH-Cas9-PpPIEZO2

For expression of Cas9 and a guide RNA (protospacer: 5’ - CTGCGCATTTGCTTTGCTGT - 3’ (Extended Data Fig. 1c), chosen with CRISPOR program^13^) targeting *PpPIEZO2* (Pp3c3_17170V3.1). Primers for guide RNA (5’-CCATACAGCAAAGCAAATGCGCAG-3’ and 5’-AAACCTGCGCATTTGCTTTGCTGT-3’) were annealed and ligated into pENTR-PpU6P-sgRNA-L1L2 and as described in^14^. Subsequently, guide RNA expression construct was moved into pMH-Cas9-gate vector^14^, via LR recombination.

#### pTKUBI-AY

For expression of cytosolic Aequorin-YFP (AY)^15^ from the *UBQ* promoter. The AY coding sequence was amplified (For. 5’-CACCATGGTGAGCAAGGGCGAGG-3’ and Rev. 5’-TTAGGGGACAGCTCCACCGTAG-3’) and inserted into pENTR vector (D-TOPO reaction, LifeTech). Subsequently, the AY coding region was moved into expression vector pTKUBI-gate^16^, via LR reaction (LifeTech). Before moss transformation, the vector was linearized with PmeI.

#### pTHUBI-oPpPIEZO1-mGFP

For overexpression of human codon-optimized *PpPIEZO1* (*oPpPIEZO1*) C-terminally tagged with mGFP (monomeric enhanced GFP^17^). We used codon-optimized PIEZO CDSs (made by gene synthesis) because native plant PIEZO genes are unstable in *E. coli*^18^. o*PpPIEZO1* coding sequence was synthesized with appropriate attB sites and inserted into pDONR221-P1P5r. Using LR+ recombination *oPpPIEZO1* and mGFP (in pDONR-L5L2) were inserted into pTHUBI-gate expression vector (UBQ promoter)^19^.

#### pTHUBI-oPpPIEZO2-mGFP

Same as for *pTHUBI-oPpPIEZO1-mGFP* but with a codon-optimized version of *PpPIEZO2*.

#### pTHUBI-oAtPIEZO1-mGFP

For overexpression of human codon-optimized *AtPIEZO1* (from *Arabidopsis thaliana*) C-terminally tagged with mGFP. The *oAtPIEZO1* coding sequence was amplified (5’ – GGGGACAAGTTTGTACAAAAAAGCAGGCTTAATGGCTTCCTTTCTGGTGG – 3’ and 5’ – GGGGACAACTTTTGTATACAAAGTTGTGGCATCGTAATCCAGCTTTG – 3’) and inserted into pDONR221-P1P5r. Using LR+ recombination, *oPpPIEZO1* and mGFP (in pDONR-L5L2) were inserted into a pTHUBI-gate expression vector (UBQ promoter)^19^.

#### pTZUBI-oPpPIEZO1-mGFP

Same as for *pTHUBI-oPpPIEZO1-mGFP* but with the pTZUBI-gate^16^ backbone. This vector was used for the transient transformation of protoplasts in Fig. 2a.

#### pTZUBI-oPpPIEZO2-mGFP

Same as for *pTHUBI-oPpPIEZO2-mGFP* but with the using pTZUBI-gate^16^ backbone. This vector was used for the transient transformation of protoplasts in Fig. 2a.

#### pMK-Cas9-PP108

For expression of Cas9 and a guide RNA (protospacer: 5’ - GAATTGGTACCAGGCTGGGT - 3’, chosen with CRISPOR program^13^) targeting redundant PP108 (Pp3c20_980V3.1) locus. Primers for guide RNA (5’-CCATGAATTGGTACCAGGCTGGGT-3’ and 5’-AAACACCCAGCCTGGTACCAATTC-3’) were annealed and ligated into pENTR-PpU6P-sgRNA-L1L2 and as described in^14^. Subsequently, guide RNA expression construct was moved into pMK-Cas9-gate vector^14^, via LR recombination. This construct was co-transformed with pTHUBI vectors, to facilitate more efficient integration into the PP108 locus.

#### pGEM-gPP1-mGFP-Kan

For genomic tagging (knock-in) of *PpPIEZO1* with mGFP (Extended Data Fig. 1c). The *PpPIEZO1*, 5’ and 3’ homology arms were amplified (5’ arm: 5’-GGGGACAAGTTTGTACAAAAAAGCAGGCTATTTAAATTTGACGCTCAACAAGGAG-3’ / 5’ - GGGGACAACTTTTGTATACAAAGTTGTCTCCAACTTTTGTATACTCCATC-3’ and 3’ arm 5’ - GGGGACAACTTTGTATAATAAAGTTGCTATAGCATTTTTGAGGATTAAATG - 3’ / 5’ - GGGGACCACTTTGTACAAGAAAGCTGGGTAATTTAAATGGCTCTCAGGACTATCTTAC – 3’), and inserted (BP recombination reaction) into pDONR221-P1P5r and pDONR221-P3-P2, respectively. The two arms were combined with mGFP (in pDONR221-L5L4) and Kanamycin resistance cassette (in pDONR221-R4R3) into pGem-gate^17^ using LR+ recombination. Before moss transformation, the vector was linearized with SwaI.

#### pGEM-gPP2-mGFP-Kan

For genomic tagging (knock-in) of *PpPIEZO2* with mGFP (Extended Data Fig. 1c). The *PpPIEZO2* 5’ and 3’ homology arms were amplified (5’ arm: 5’-GGGGACAAGTTTGTACAAAAAAGCAGGCTCTCGAGAAATAGTACATGTATGGACTTGTG-3’ / 5’ – GGGGACAACTTTTGTATACAAAGTTGTCTCAACTTTTGTGTATTCCATAAG - 3’ and 3’ arm 5’-GGGGACAACTTTGTATAATAAAGTTGCTCCTCTTTGTTGCGGGAGAAG - 3’ / 5’ – GGGGACCACTTTGTACAAGAAAGCTGGGTACTCGAGAGATGCATTGGACTTTACTTTTG – 3’), and inserted (BP recombination reaction) into pDONR221-P1P5r and pDONR221-P3-P2, respectively. The two homology arms were combined with mGFP (in pDONR221-L5L4) and Kanamycin resistance cassette (in pDONR221-R4R3) into pGem-gate^17^ using LR+ recombination. Before moss transformation, the vector was linearized with XhoI.

#### pMH-Cas9_PP2_R2508

For expression of Cas9 and a guide RNA (protospacer: 5’ - TGGCAGTTGGGAGGTTTATT - 3’, chosen with CRISPOR program^13^) targeting *PpPIEZO2*. Primers for guide RNA (5’-CCATTGGCAGTTGGGAGGTTTATT-3’ and 5’-AAACAATAAACCTCCCAACTGCCA-3’) were annealed and ligated into pENTR-PpU6P-sgRNA-L1L2 and as described in^14^. Subsequently, guide RNA expression construct was moved into pMH-Cas9-gate vector^14^, via LR recombination. This construct was used together with annealed oligonucleotides (5’-TGGCAGTTGGGAGGTTTATTAAGTTGCAGTGCGCCGATATCCG – 3’ / 5’ – CGGATATCGGCGCACTGCAACTTAATAAACCTCCCAACTGCCA – 3’ for R2508K and 5’ – TGGCAGTGGGGAGGTTTATTCACCTGCAGTGCGCCGATATCCGT – 3’ / 5’ – ACGGATATCGGCGCACTGCAGGTGAATAAACCTCCCCACTGCCA for R2508H) to introduce point mutations in the genome via oligodeoxynucleotide-assisted homology-directed repair^20^.

#### pMH-Cas9_PP2_R2549

Same as for *pMH-Cas9_PP2_R2508*, but using different guide RNA (5’-AGCTGGGGAGTTGGAGCTAG – 3’, made with 5’-CCATAGCTGGGGAGTTGGAGCTAG – 3’ / 5’ – AAACCTAGCTCCAACTCCCCAGCT – 3’ annealed primer) and annealed oligonucleotides (5’ – GCAGCTGGGGAGTTGGAGCTCGAAGGTTTGTTCTGGACATTAAT – 3’ / 5’ – ATTAATGTCCAGAACAAACCTTCGAGCTCCAACTCCCCAGCTGC – 3’ for PP2 R2549del).

### Moss DNA isolation

For DNA isolation, a 2-3 mm clump of moss tissue (protonema or gametophores) was mechanically disrupted (using zirconia/silica beads and a mixer mill (30 swings per s for 5 min) and mixed with 400 μL of Shorty Buffer (200 mM Tris-Cl pH 9, 400 mM LiCl, 25 mM EDTA and 1% SDS). Cellular debris was spun down for 5 min at 21,000xg at RT and the supernatant was mixed 1:1 with isopropanol. After 10 min at −20°C, the solution was centrifuged for 10 min at 21,000xg at RT, the supernatant discarded, and the DNA pellet was resuspended in 300 μL of TE buffer. DNA was re-precipitated with 30 μL of 3M Sodium acetate and 800 μL of absolute ethanol. The solution was incubated at −20°C (2 h), spun down for 10 min at 21000xg, washed once with 70% ethanol, spun down again for 5 min at 21000xg. The supernatant was discarded, and the pellet was air-dried and resuspended in 400 μL of TE.

### Hypoosmotic shock and cytosolic calcium measurement

Moss AY lines grown on BCDAT plates were ground and spread on cellophaned BCDAT plates supplemented with 400 mM mannitol, cultured for 5 days, then transferred to liquid BCDAT + 400 mM mannitol and kept in the growth chamber for another 20 h. Subsequently, plants were transferred to the same media with 1 μM coelenterazine, incubated in dark for 2 h, washed three times in BCDAT + 400 mM mannitol media, evenly distributed in wells of opaque white 96-well plates (50 μL of media per well) and left to adjust in the dark for 30 min (Extended Data Fig. 2a). Ca^2+^-dependent aequorin bioluminescence was measured in a TECAN Pro200 plate reader with a 1 s integration time. Hypoosmotic stress was induced by an automatic injection of 150 μL of double-distilled water (~330 mOsm drop). For the isosmotic control, 150 μL of BCDAT + 400 mM mannitol media was injected. After iso- or hypo-osmotic treatment, 100 μL of 1 M CaCl_2_ in 10% ethanol was injected into each well to discharge all remaining aequorin. The cytosolic free calcium concentration was calculated with the following formula: [Ca^2+^]_*cyt*_ (μM) = 10^6 * 10^-(5.5593-0.332588log_10_(C/R)); where C corresponds to counts recorded per time interval (one second) and R to total remaining counts including discharge^21,22^.

### Moss growth assays

In order to test the growth of different moss lines, a small amount (area of ~2 mm^2^) of 7-day-old protonema tissue (from regular culture plates, see above), was shaped (using sterile tweezers) into a ball and placed on top of cellophane covered BCDAT plate. A similar amount of starting tissue was used for all moss lines tested. Under these conditions, the protonema tissue both grew in overall volume and on the edges produced caulonemal filaments^23^, which spread the plant (Fig. 1b, c, Extended Data Figs. 1g, 3b, 6a, 7a). Plates were sealed with breathable 3M surgical tape and cultured horizontally for 7 days under standard light conditions. Images of the growth assays were taken with a THUNDER Imager Model Organism Microscope (Leica) using a greyscale camera, 1x objective, and background illumination. Multi-tile imaging and subsequent stitching (25% overlap) was used to capture images of whole plates.

To determine the growth rates of the caulonemal cells on the plant edge (Expanded Data Figs. 1e, i; 6b; 7d), 7-day-old tissue was combined into 2 mm clumps and placed on 2×2 cm pieces of cellophane (3 clumps per cellophane piece). After 6 days of growth, plants were imaged with THUNDER Imager Model Organism Microscope (Leica) focusing on the plant edge. We made a 4 or 8 h time-lapses (15 min per frame; 1x objective, greyscale camera, and background illumination) for each individual plant (Supplementary Fig. 4), utilizing systems motorized stage. Plates were kept sealed during imaging.

For dark/gravity growth assay (Fig. 1d), the 7-day-old protonemal tissue was shaped into logs (~1 cm long and ~2mm wide) and placed on top of 2×2 cm pieces of cellophane (one log per cellophane), layered over BCDAT media supplemented with 0.2% (w/v) of glucose. Plates were sealed with breathable 3M surgical tape and cultured vertically for 48 h under light, followed by 10 days in the dark. Pictures were taken with the THUNDER Imager Model Organism Microscope (Leica) using a greyscale camera, 1x objective, and background illumination.

### Moss mounting, staining, and imaging

For imaging transformed protoplasts (Fig. 2A; Extended Data Figs. 3a), 10 μL of protoplast suspension was placed on the center on the slide with two small (22×22 mm) coverslips on either side as spacers. Over that, a larger (44 x 22 mm) coverslip was placed so it rested on the smaller ones. This insured that protoplasts would not be crushed by the top coverslip. The coverslips were fixed into position with drops of melted VALAP (1:1:1 mixture by weight of vaseline, paraffin, and lanolin).

In order to image caulonemal cells on the plant edge (Figs. 1c; 2c, d; 3a, c-e; 4d-g; Extended Data Figs. 3c, d; 4a-c, f; 5a, c; 6c, 7e), we grew protonemal tissue as described for growth assays. Clumps (~2 mm) of 7-day-old protonemal tissue were placed on cellophaned BCDAT plates and cultured for 6 days. For imaging, the whole plant was detached from the cellophane, by adding 5-10 μL of liquid BCDAT media and placed on a coverslip at the bottom of an empty 3.5 cm Petri dish. Liquid band-aid (CVS brand) was used to attach the center of the plant to the coverslips, but care was taken that the band-aid did not reach caulonemal cells on the edge; 0.5 – 1 μL was usually sufficient. After drying for 45-60s, the Petri dish was filled with 3 mL of liquid BCDAT media.

In some cases (Figs. 2c, d; 3a; Extended Data Figs. 3c, d), cells were imagined directly without any staining, while in others they were first stained. For **MDY64** staining (labels the vacuolar membrane, cell wall/plasma membrane, and vesicles) (Figs. 3d, e; 4d-g; Extended Figs. 4a-c, f; 5a, c, 6c, 7e), plants were first fixed to the coverslip as described above, then stained in BCDAT + 1 μM MDY-64 (Thermofisher Y7536) for 5 min in dark, washed twice with BCDAT for 1 min in light and finally mounted in 3 mL of the BCDAT media. For **BCECF** staining (accumulates in the vacuolar lumen) (Fig. 3c), plants were detached from cellophane, submerged into BCDAT + 10 μM BCECF, AM (2’,7’-Bis-(2-Carboxyethyl)-5-(and-6)-Carboxyfluorescein, Acetoxymethyl Ester; Thermofisher B1150) + 0.02% Pluronic F-127 for 1 hour in the dark, washed twice with BCDAT for 5 min in light and mounted to the coverslip as described above. For **Calcofluor-White** staining (labels cell walls) (Fig. 1c), plants were detached from cellophane, submerged into BCDAT + 0.1 mg/mL Calcofluor-White (Sigma-Aldrich F3543) for 5 min in the dark, washed twice with BCDAT for 2 min in light and mounted to the coverslip as described above.

Most of the confocal images (Figs. 1c; 2a, c; 3a, c-e; 4d-g; Extended Data Figs. 3a, c; 4a-c, f; 5a, c; 6c; 7e)) here made with an inverted Olympus FV3000 confocal microscope with UPLSAPO 60XW NA1.2 objective, Galvo scan unit, and High Sensitivity-Spectral Detectors, except imaged in Figs. 2d; Extended Data Fig 3d, which were made with an inverted Leica Sp8 Lightning Single Photon confocal microscope (HC PL APO CS2 63Xoil NA1.4 objective, Galvo scan unit, and High Sensitivity-Spectral Detectors. **mGFP** (Figs. 2; Extended Data Fig. 3a, c, d; Supplementary Fig.5) was excited by a 488 nm laser and detected in 500 - 550 nm range. **mCheery** (Figs. 2a; Extended Data Fig. 3a) was excited by a 561 nm laser and detected in the 570 - 610 nm range. **MDY64** (Figs. 3d, e; 4d-g; Extended Data Fig. 4a-c, f; 5a, c; 6c; 7e) was excited by a 445 nm laser and detected in the 480-540 nm range. **BCECF** (Fig. 3c) was excited by 445 and 448 nm lasers and in both cases detected in the 500-550 nm range. **Calcofluor-White** (Fig. 1c) was excited by a 405 nm laser and detected in the 415-460 nm range. **Chlorophyll autofluorescence** (Figs. 1c; 2a, d; 3d, e; 4d-g; Extended Data Fig. 3a, d; 4a-c, f; 5a, c; 6c; 7e; Supplementary Fig.5) was excited by 405, 445, 488, or 561 nm lasers and detected in the 660 - 700 nm range. **Bright-field** (Fig. 2c, 3a; Extended Data Fig. 3c; Supplementary Fig.5) was imaged using transmitted light detectors.

### Image processing and quantification

For analysis of plant size (Extended Data Fig. 1d, i) (defined as the area occupied by the plan, including the spreading caulonema filaments), stitched images were exported as TIFF and analyzed in Fiji^24^. After the threshold (max entropy) was set, the individual plants were identified, and their area measured with Analyse particles command.

To measure caulonema tip growth speed (Extended Data Fig. 1e, h; 6b; 7d) the time-lapse videos made with Leica THUNDER images (see above; Supplementary Fig. 4) were exported as TIFF files and processed in Fiji. To measure the growth speed, a segmented line was drawn through the center of the growing caulonema filaments and used to generate a kymogram (see examples in Supplementary Fig. 4). The average tip growth speed was calculated from the distance the tip traversed over a specific time period.

To calculate an average filament curvature (Fig. 1f), we used the first frame of the time-lapse video made for tip growth rate measurement (Supplementary Fig. 4). A segmented line was drawn through the center of the caulonema filaments and used to fit a spline. The curvature radii along the draw spline were calculated with a script from Olivier Burri (Ecole Polytechnique Fédérale de Lausanne; https://gist.github.com/lacan/42f4abe856f697e664d1062c200fd21f). The curvature radii (in meters) were averaged for each filament and its reciprocal (curvature – k) was calculated. The obtained curvature values were converted to logarithms to obtain Gaussian distribution.

To determine the percentage of cells with expanded vacuoles in the tip regions of caulonemal cells (Fig. 3b; Extended Data Fig. 4d; 5b, d, 7f), moss was imaged using bright-field on Olympus FV3000 confocal microscope (UPLSAPO 60XW NA1.2 objective) and cells were manually classified as either having tips filled with cell material (Fig. 3a top panel) or have large and expanded vacuole (Fig. 3a bottom panel). For each plant, approximately 50 apical caulonemal cells were evaluated and the percentage of cells with expanded vacuoles was counted as one data point.

Subapical caulonemal cell length and width (Fig. 1e; Extended Data Fig. 1f) was quantified in Fiji^24^ using Maximum Z projections of Calcofluor-White stained moss images (Fig.1c). Cell length and width were measured by drawing a line along or perpendicular to the longitudinal axis of the cell, respectively. To estimate the cell width, 3 perpendicular lines were drawn in different parts of the cell and their lengths averaged.

Confocal microscopy images in Figs. 1c; 2a, c; 3c-e; 4d-g; Extended Data Fig. 3a, c; 4a-c, f; 5a, c; 6c; 7e; Supplementary Fig.5 were subjected to deconvolution (Olympus CellSens software, Advanced Maximum Likelihood Algorithm with 3 or 5 iterations). Color (LUT, lookup table) and display intensity range of individual channels were adjusted to facilitate clarity. The images that are meant to be compared directly (e.g. WT vs. knock-in lines in Extended Data Fig. c or images in Fig. 4e-g) were adjusted to the same intensity range. The intensity of the chlorophyll autofluorescence channel was adjusted independently from the main channel. After adjustments, the deconvolved images were captured with Microsoft Snipping Tool, saved as Portable Network Graphic (PNG) files and imported into Adobe Illustrator. Confocal microscopy images in Figs. 2d and Extended Data Fig. 3d were deconvolved using Leica LAS X software (an adaptive strategy with 5 iterations), adjusted as described above, and exported as PNG files. The following lookup tables (LUTs) were used: Calcofluor-White – Cyan; mGFP – Green; mCherry – Magenta; BCECF (445 nm excitation channel) – Magenta; MDY64 – Fall (a range LUT (from black to white over orange/yellow shades) from Olympus FV31S-SW software); Chlorophyll Autofluorescence – Red, Magenta or Cyan. No images used for data quantification were subjected to deconvolution.

To determine the percentage of apical caulonemal cells with different vacuolar morphologies in the tip region (Fig. 4b), deconvolved Z-stack images were manually inspected and cells were classified into one of the 5 vacuolar morphology categories. To minimize the influence of cell growth stage and other factors on vacuolar morphology variation, only mature apical caulonema cells with the expanded vacuoles in the region behind the nucleus and with overall similar cell morphology (e.g. WT cells in Fig.3a, d) were included in the quantification shown in Fig. 4b.

### Statistical analysis

All statistical analysis was performed in GraphPad Prism software, except Fisher’s Exact test which was performed in IBM’s SPSS software. Specific tests used are indicated in the figure legends. In all tests, p<0.05 was used as a cutoff for significance.

### Western blots

9-day old moss plants regenerated from protoplasts were collected, excess water was removed with tissue paper, and they were ground to a fine powder in a mortar and pestle with liquid nitrogen. The powder was transferred to an Eppendorf tube, mixed 1:20 (mg:μL) with 2X sample buffer (100 mM TRIS pH6.8, 4% (w/v) SDS, 0.2% (w/v) Bromophenol Blue, 20% Glycerol (v/v), 1x protein inhibitor cocktail and 2% (v/v) mM beta-mercaptoethanol). The mixture was then vortexed, freeze-thawed in liquid nitrogen three times, incubated at 80°C for 15 min, and spun down at 21000xg for 1 min. Proteins were separated on an SDS-PAGE gel (8% running and 4% stacking gel, 1.5 mm thickness), and transferred to a PVDF membrane (BioRad) for 16 h at 4°C using a wet transfer system from BioRad (transfer buffer: 20 mM Tris, 150 mM Glycine, 20% (v/v) Methanol and 0.05% (w/v) SDS). For detection, the blot was blocked in 5% milk in TBS-T for 8h, incubated overnight at 4°C with anti-GFP (Takara 632380, 1:10000), washed in TBS-T, and then incubated with anti-mouse-HRP (Millipore-Sigma F0257 1:5000) for 1h at RT. The blot was washed in TBS-T, then detected with SuperSignal™ West Femto substrate. After detection, proteins on the PVDF membrane were stained with Coomassie for 5 min in 50% (v/v) methanol, 7% (v/v) acetic acid and 0.1% (w/v) Coomassie Blue R; destained for 5 min in 50% (v/v) methanol and 7% (v/v) acetic acid, rinsed in 90% (v/v) methanol and 10% (v/v) acetic acid, and finally washed in water^25^.

## References

1. Bagriantsev, S. N., Gracheva, E. O. & Gallagher, P. G. Piezo proteins: Regulators of mechanosensation and other cellular processes. J. Biol. Chem. 289, 31673–31681 (2014).

2. Wu, J., Lewis, A. H. & Grandl, J. Touch, Tension, and Transduction – The Function and Regulation of Piezo Ion Channels. Trends Biochem. Sci. 42, 57–71 (2017).

3. Coste, B. et al. Piezo1 and Piezo2 are essential components of distinct mechanically activated cation channels. Science 330, 55–60 (2010).

4. Zhang, Z. et al. Genetic analysis of a Piezo-like protein suppressing systemic movement of plant viruses in Arabidopsis thaliana. Sci. Rep. 9, 3187 (2019).

5. Durand-Smet, P. et al. A comparative mechanical analysis of plant and animal cells reveals convergence across kingdoms. Biophys. J. 107, 2237–2244 (2014).

6. Anishkin, A., Loukin, S. H., Teng, J. & Kung, C. Feeling the hidden mechanical forces in lipid bilayer is an original sense. Proc. Natl. Acad. Sci. U. S. A. 111, 7898–7905 (2014).

7. Ranaade, S. S., Syeda, R. & Patapoutian, A. Mechanically Activated Ion Channels. Neuron 87, 1162–1179 (2015).

8. Basu, D. & Haswell, E. S. Plant mechanosensitive ion channels: an ocean of possibilities. Curr. Opin. Plant Biol. 40, 43–48 (2017).

9. Cox, C. D., Bavi, N. & Martinac, B. Bacterial Mechanosensors. Annu. Rev. Physiol. 80, 71–93 (2018).

10. Prole, D. L. & Taylor, C. W. Identification and Analysis of Putative Homologues of Mechanosensitive Channels in Pathogenic Protozoa. PLoS One 8, (2013).

11. Rensing, S. A., Goffinet, B., Meyberg, R., Wu, S & Bezanilla, M. The Moss Physcomitrium (Physcomitrella) patens : A Model Organism for Non-Seed Plants. Plant Cell 1997, (2020).

12. Kofuji, R. & Hasebe, M. Eight types of stem cells in the life cycle of the moss Physcomitrella patens. Curr. Opin. Plant Biol. 17, 13–21 (2014).

13. Pietra, A. Della, Mikhailov, N. & Giniatullin, R. The emerging role of mechanosensitive piezo channels in migraine pain. Int. J. Mol. Sci. 21, (2020).

14. Mehlmer, N. et al. A toolset of aequorin expression vectors for in planta studies of subcellular calcium concentrations in Arabidopsis thaliana. J. Exp. Bot. 63, 1751–1761 (2012).

15. Tuteja, N. & Mahajan, S. Calcium signaling network in plants: An overview. Plant Signal. Behav. 2, 79–85 (2007).

16. Gudipaty, S. A. et al. Mechanical stretch triggers rapid epithelial cell division through Piezo1. Nature 543, 118–121 (2017).

17. Maathuis, F. J. M. Vacuolar two-pore K+ channels act as vacuolar osmosensors. New Phytol. 191, 84–91 (2011).

18. Zhou, X. L. et al. The transient receptor potential channel on the yeast vacuole is mechanosensitive. Proc. Natl. Acad. Sci. U. S. A. 100, 7105–7110 (2003).

19. Schönknecht, G. Calcium signals from the vacuole. Plants 2, 589–614 (2013).

20. Peiter, E. The plant vacuole: Emitter and receiver of calcium signals. Cell Calcium 50, 120–128 (2011).

21. Tan, X. et al. A Review of Plant Vacuoles: Formation, Located Proteins, and Functions. Plants 8 (2019).

22. Cui, Y., Zhao, Q., Hu, S. & Jiang, L. Vacuole Biogenesis in Plants: How Many Vacuoles, How Many Models? Trends Plant Sci. 25, 538–548 (2020).

23. Jensen, L. C. W. Division, Growth, and Branch Formation in Protonema of the Moss Physcomitrium turbinatum: Studies of Sequential Cytological Changes in Living Cells. Protoplasma 107, 301–317 (1981).

24. Yi, P. & Goshima, G. Transient cotransformation of CRISPR/Cas9 and oligonucleotide templates enables efficient editing of target loci in Physcomitrella patens. Plant Biotechnol. J. 18, 599–601 (2020).

25. Dünser, K. et al. Extracellular matrix sensing by FERONIA and Leucine-Rich Repeat Extensins controls vacuolar expansion during cellular elongation in Arabidopsis thaliana. EMBO J. 38, (2019).

26. Kellermayer, R., Aiello, D. P., Miseta, A. & Bedwell, D. M. Extracellular Ca2+ sensing contributes to excess Ca2+ accumulation and vacuolar fragmentation in a pmr1Δ mutant of S. cerevisiae. J. Cell Sci. 116, 1637–1646 (2003).

## Methods References

2. Prole, D. L. & Taylor, C. W. Identification and Analysis of Putative Homologues of Mechanosensitive Channels in Pathogenic Protozoa. PLoS One 8, (2013).

3. Möller, S., Croning, M. D. R. & Apweiler, R. Evaluation of methods for the prediction of membrane spanning regions. Bioinformatics 17, 646–653 (2001).

4. Kumar, S., Stecher, G. & Tamura, K. MEGA7: Molecular Evolutionary Genetics Analysis Version 7.0 for Bigger Datasets. Mol. Biol. Evol. 33, 1870–1874 (2016).

5. Hall, G. B. Phylogenetic Trees Made Easy: A How-To Manual. 5^th^ edition. (Sinauer Associates, 2017).

6. Edgar, R. C. MUSCLE: multiple sequence alignment with high accuracy and high throughput. Nucleic Acids Res. 32, 1792–1797 (2004).

7. Le, S. Q. & Gascuel, O. An improved general amino acid replacement matrix. Mol. Biol. Evol. 25, 1307–1320 (2008).

8. Saitou, N. & Nei, M. The neighbor-joining method: a new method for reconstructing phylogenetic trees. Mol. Biol. Evol. 4, 406–425 (1987).

9. Jones, D. T., Taylor, W. R. & Thornton, J. M. The rapid generation of mutation data matrices from protein sequences. Bioinformatics 8, 275–282 (1992).

10. Cove, D. J. et al. Culturing the moss *Physcomitrella patens*. Cold Spring Harb. Protoc. 4 (2009).

11. Cove, D. J. et al. Isolation and regeneration of protoplasts of the moss *<u>Physcomitrella patens</u>*. Cold Spring Harb. Protoc. 4 (2009).

12. Cove, D. J. et al. Transformation of the moss *Physcomitrella patens* using direct DNA uptake by protoplasts. Cold Spring Harb. Protoc. 4 (2009).

13. Haeussler, M. et al. Evaluation of off-target and on-target scoring algorithms and integration into the guide RNA selection tool CRISPOR. Genome Biol. 17 (2016).

14. Mallett, D. R., Chang, M., Cheng, X. & Bezanilla, M. Efficient and modular CRISPR-Cas9 vector system for *Physcomitrella patens*. Plant Direct 3, 1–15 (2019).

15. Mehlmer, N. et al. A toolset of aequorin expression vectors for in planta studies of subcellular calcium concentrations in *Arabidopsis thaliana*. J. Exp. Bot. 63, 1751–1761 (2012).

16. Wu, S. Z. & Bezanilla, M. Myosin VIII associates with microtubule ends and together with actin plays a role in guiding plant cell division. Elife 3, e03498 (2014).

17. Vidali, L. et al. Rapid formin-mediated actin-filament elongation is essential for polarized plant cell growth. Proc. Natl. Acad. Sci. U. S. A. 106, 13341–13346 (2009).

18. Zhang, Z. et al. Genetic analysis of a Piezo-like protein suppressing systemic movement of plant viruses in *Arabidopsis thaliana*. Sci. Rep. 9, 3187 (2019).

19. Vidali, L., Augustine, R. C., Kleinman, K. P. & Bezanilla, M. Profilin Is Essential for Tip Growth in the Moss *Physcomitrella patens*. Plant Cell 19, 3705–3722 (2007).

20. Yi, P. & Goshima, G. Transient cotransformation of CRISPR/Cas9 and oligonucleotide templates enables efficient editing of target loci in *Physcomitrella patens*. Plant Biotechnol. J. 18, 599–601 (2020).

21. Plieth, C. Aequorin as a reporter gene. Methods Mol. Biol. 323, 307–327 (2006).

22. Knight, H., Trewavas, A. J. & Knight, M. R. Cold Calcium Signaling in *Arabidopsis* Involves Two Cellular Pools and a Change in Calcium Signature after Acclimation. Plant Cell 8, 489–503 (1996).

23. Thelander, M., Olsson, T. & Ronne, H. Effect of the energy supply on filamentous growth and development in *Physcomitrella patens*. J. Exp. Bot. 56, 653–662 (2005).

24. Schindelin, J. et al. Fiji: an open-source platform for biological-image analysis. Nat. Methods 9, 676–682 (2012).

25. Welinder, C. & Ekblad, L. Coomassie staining as loading control in Western blot analysis. J. Proteome Res. 10, 1416–1419 (2011).

