## Supplemental Files for "Regulation of Vacuole Morphology by PIEZO Channels in Spreading Earth Moss"

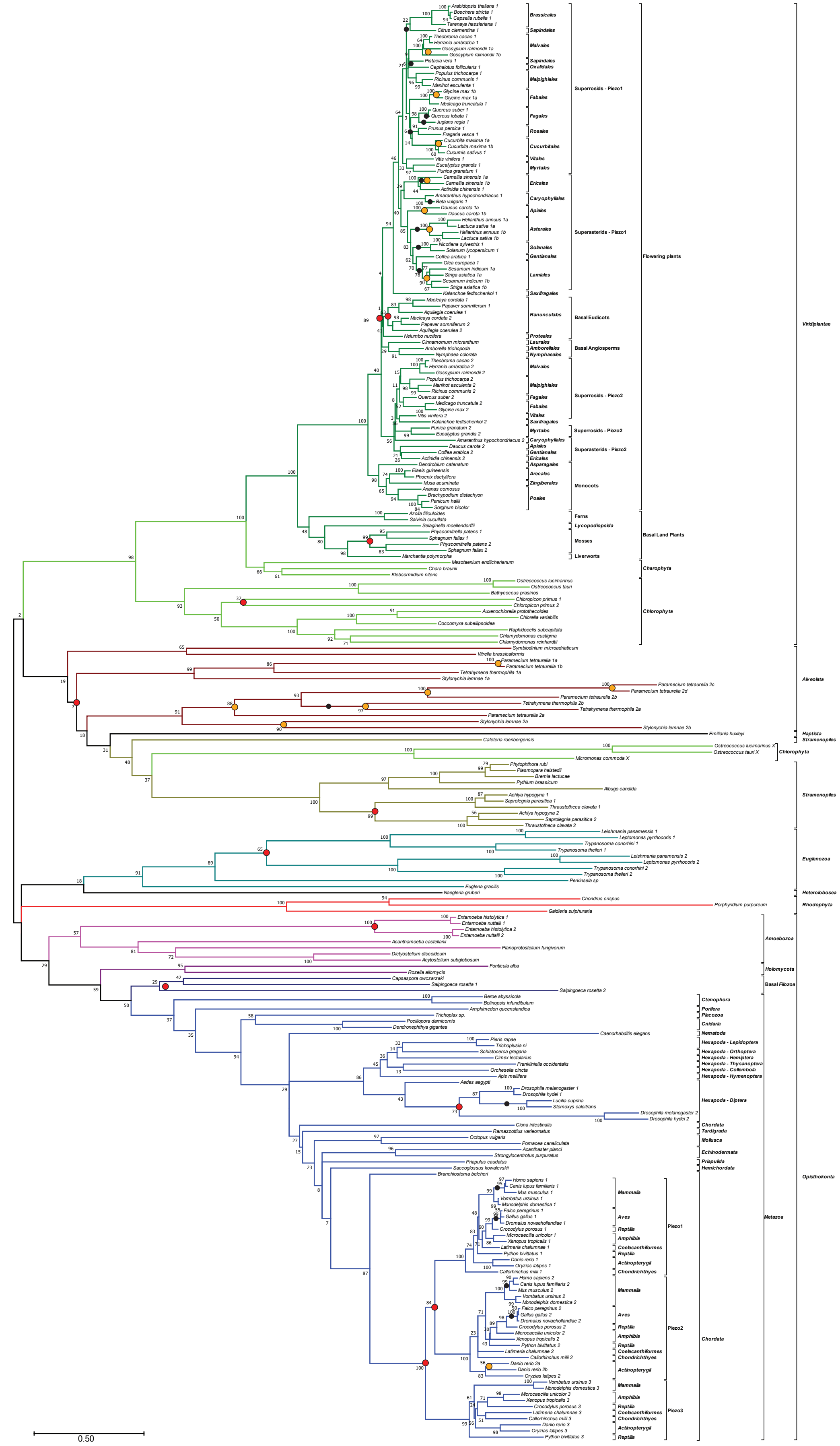


**Supplementary Figure 1. Maximum Likelihood Unrooted Phylogenetic Tree of PIEZO Homologs.**
This is a more detailed version of the tree presented in Fig. 1a. The protein sequences of PIEZO domains from 235 homologs were used to generate a Maximum Likelihood Tree. Protein sequences of conserved PIEZO domain (Extended Data Fig. 1b) were aligned using MUSCLE (default settings in MEGA7). The tree is drawn to scale with branch lengths corresponding to evolutionary distances expressed as amino acid substitutions per site. The reliability of each node was tested with 500 bootstrap replications and the resulting values are depicted next to each node. Genes are labeled with species names followed by a number (denoting homologs that putatively arose through gene duplication of an ancestral *PIEZO*, red circles) and a letter (denoting homologs that putatively arose from subsequent duplications of one of the *PIEZO* copies, yellow circles). Black circles mark putative losses of one homolog in that lineage. Homologs marked with X most likely arose through a horizontal gene transfer event. Accession numbers and additional details about the sequences used are listed in Supplementary Table 1.

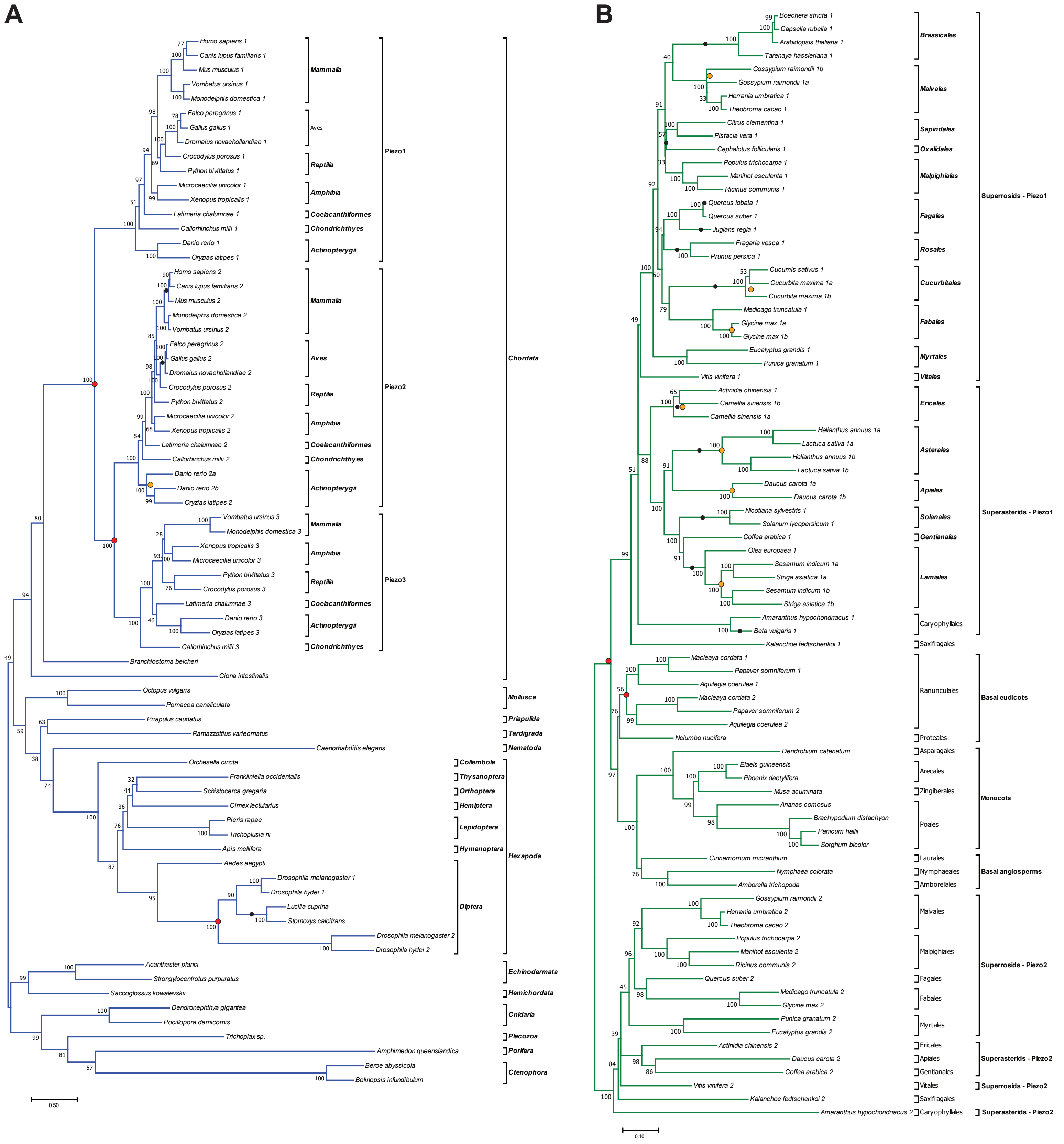


**Supplementary Figure 2. Maximum Likelihood Unrooted Phylogenetic Trees of Animal and Angiosperm PIEZO homologs.** Maximum Likelihood Trees of 73 PIEZO animal homologs (**a**) and 83 PIEZO homologs from angiosperms/flowering plants (**b**). Full-length protein sequences of PIEZO homologs were aligned using MUSCLE (default settings in MEGA7). Both trees were drawn to scale with branch lengths corresponding to evolutionary distances expressed as amino acid substitutions per site. The reliability of each node was tested with 500 bootstrap replications and the resulting values are depicted next to each node. Genes are labelled with species names followed by a number (denoting homologs that putatively arose through gene duplication of an ancestral *PIEZO*, red circles) and/or a letter (denoting homologs that putatively arose from subsequent duplications of one of the *PIEZO* copies, yellow circles). Black circles mark putative losses of one homolog in that lineage. Homologs marked with X most likely arose through a horizontal gene transfer event. Accession numbers and additional details about the sequences used are listed in Supplementary Table 1.

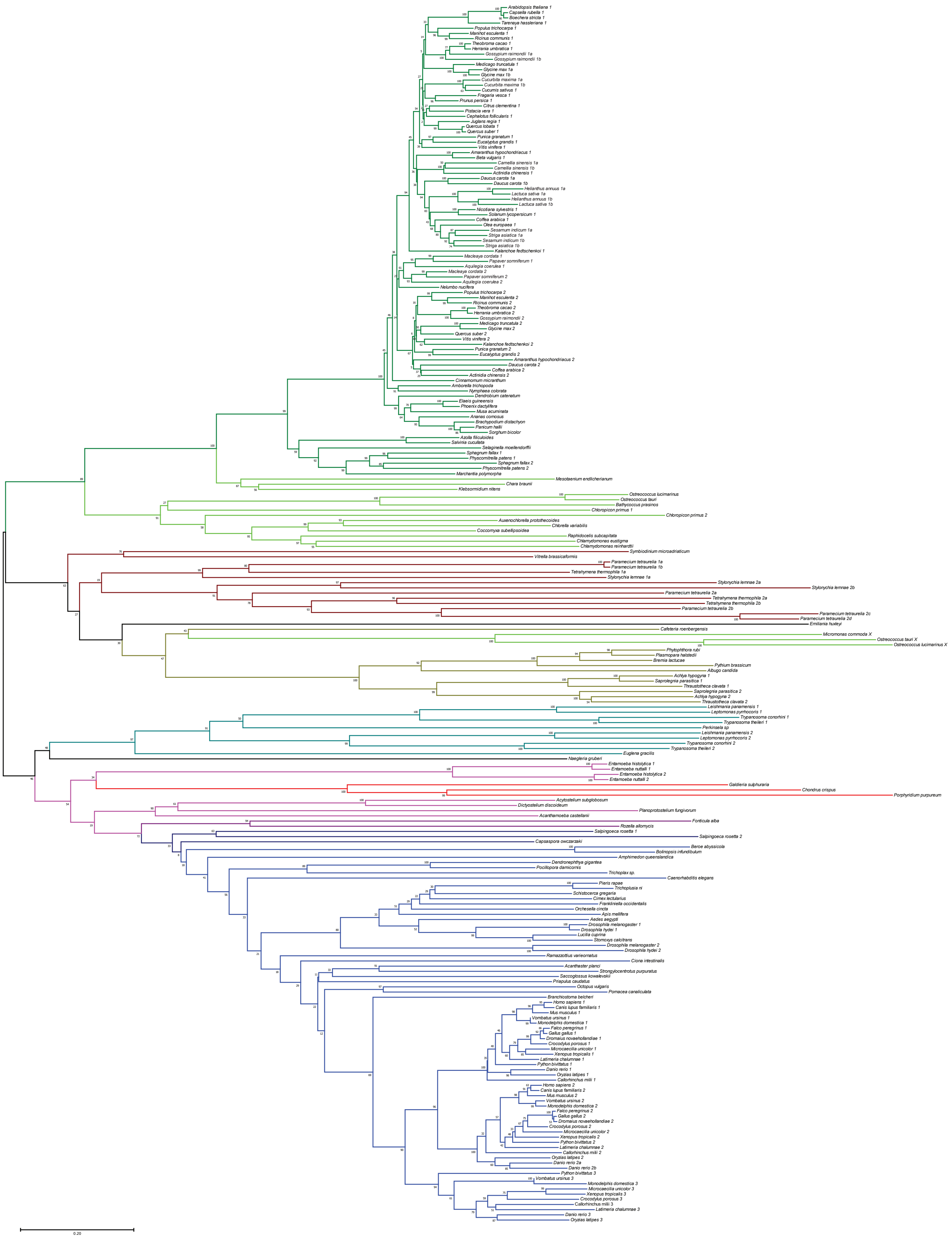

**Supplementary Figure 3. Neighbor-Joining Unrooted Phylogenetic Tree of PIEZO Homologs.**
The protein sequences of PIEZO domains from 235 homologs were aligned using MUSCLE (default settings in MEGA7) and used to generate a Neighbor-Joining Tree. The evolutionary distances were calculated with the Jones-Taylor-Thornton (JTT) model (in MEGA7) with site coverage cut-off of 90% (502 positions in final calculations). The tree is drawn to scale with branch lengths corresponding to evolutionary distances expressed as amino acid substitutions per site. The reliability of each node was tested with 1000 bootstrap replications and the resulting values are depicted next to each node.

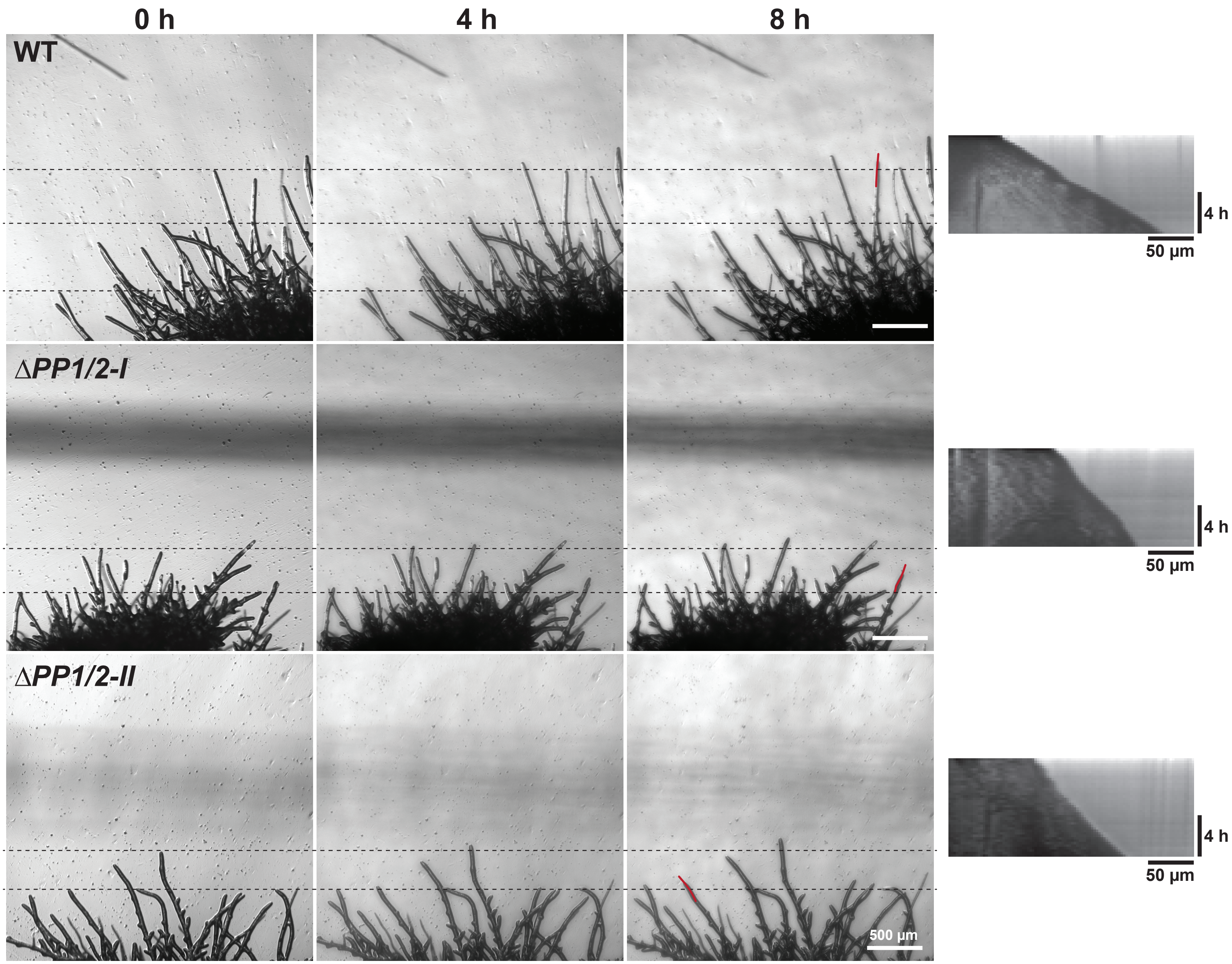


**Supplementary Figure 4. Moss growth rate assay.**
Still images from time-lapse videos of WT and Δ*PP1/2* mutants (started from fragmented protonemata) grown for 5 days on the surface of cellophaned BCDAT media. Red lines are examples of those used to generate kymographs in Fiji, shown in the right panels. Kymographs were used to calculate average growth rates by dividing the distance that the tip traversed with time.

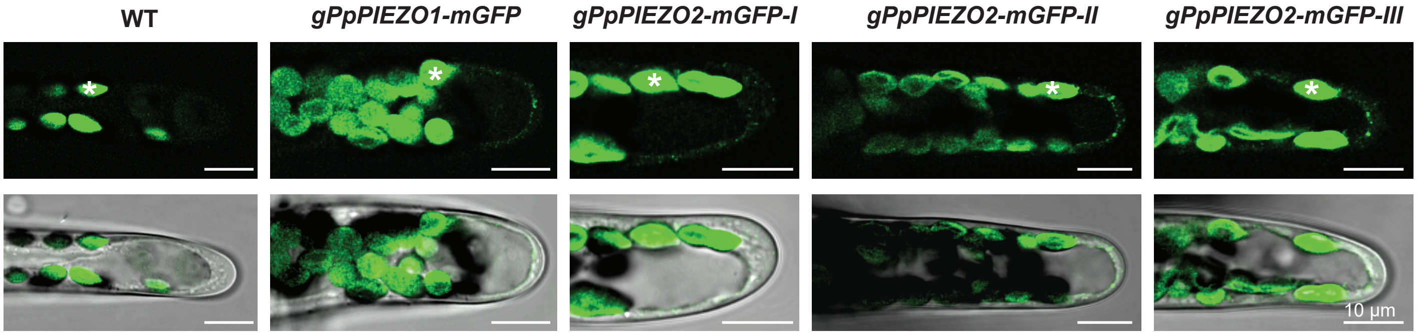


**Supplementary Figure 5. *Pp*PIEZO1 and *Pp*PIEZO2 localize to the vacuolar membrane in chloronemal cells.**
Top, deconvolved single focal plane confocal images of apical chloronemal cells from 9-day-old plants regenerated from protoplasts. Green, mGFP; asterisks, chlorophyll autofluorescence. Bottom, brightfield overlay.
