## Supplemental Table 2 for "Regulation of Vacuole Morphology by PIEZO Channels in Spreading Earth Moss"

**Supplementary Table 2. List of moss lines used in this study**

| Name | Backgro und | Vector used | Method of modification | Target locus | Observed mutation on gDNA level | Observed mutation on mRNA level | Effect |
|---|---|---|---|---|---|---|---|
| ***ΔPP1-I*** | WT | *pZeo-Cas9-PpPIEZO1* | Two CRISPR targets + NHEJ* | *PpPIEZO1* | 495 bp deletion and 3 bp insertion between two CRISPR sites | 232 bp deletion and 3 bp insertion | Frameshift and premature stop codon 31 bp after the mutations |
| ***ΔPP1-II*** | WT | *pZeo-Cas9-PpPIEZO1* | Two CRISPR targets + NHEJ* | *PpPIEZO1* | 714 bp deletion and 2 bp insertion between two CRISPR sites | 279 bp deletion, 2 bp insertion and 67 bp of 7th intron retained | Overall in-frame deletion but retained intron section contains an in-frame stop codon |
| ***ΔPPp2-I*** | WT | *pZeo-Cas9-PpPIEZO2* | Two CRISPR targets + NHEJ* | *PpPIEZO2* | 827 bp deletion between two CRISPR sites; 522 bp of which was reinserted in the opposite orientation | 321 bp deletion and 522 bp insertion (the inverted region) | Overall in-frame deletion but first 3 bp of the inverted region are in-frame stop codon |
| ***ΔPP2-II*** | WT | *pZeo-Cas9-PpPIEZO2* | Two CRISPR targets + NHEJ* | *PpPIEZO2* | 834 bp deletion between two CRISPR sites | 328 bp deletion | Frameshift and premature stop codon 5 bp after the mutations |
| ***ΔPP1/2-I*** | *ΔPP2-I* | *pZeo-Cas9-PpPIEZO1* | Two CRISPR targets + NHEJ* | *PpPIEZO1* | 490 bp deletion between two CRISPR sites | 227 bp deletion | Frameshift and premature stop codon 68 bp after the mutation |

| | | | | | | | |
|---|---|---|---|---|---|---|---|
| ***ΔPP1/2-II*** | *ΔPP2-II* | *pZeo-Cas9-PpPIEZO1* | Two CRISPR targets + NHEJ* | *PpPIEZO1* | 7 bp and 370 bp deletions at the first and second CRISPR site, respectively | 7 bp deletion in the 5th exon, 43 bp of 6th intron retained and 150 bp deletion in the 6th exon | Premature stop codon 56 bp after the first mutation (7bp del) |
| ***ΔPP1/2-III*** | *ΔPP1-II* | *pMH-Cas9-PpPIEZO2* | One CRISPR target + NHEJ* | *PpPIEZO2* | 13 bp insertion at the CRIPSR site | 13 bp insertion | Frameshift and premature stop codon 53 bp after the insertion |
| **AY** | WT | *pTKUBI-AY* | Homologous recombination-based insertion | *ARPC2 Pp1s249_67V6.1* | Insertion of expression cassette into a redundant locus | N/A | N/A |
| ***aΔPp2-I*** | AY | *pZeo-Cas9-PpPIEZO2* | Two CRISPR targets + NHEJ* | *PpPIEZO2* | 314 bp and 509 bp deletions at the first and second CRISPR site, respectively | 147 and 170 bp deletions | Frameshift and premature stop codon 93 bp after the mutations |
| ***aΔPp2-II*** | AY | *pZeo-Cas9-PpPIEZO2* | Two CRISPR targets + NHEJ* | *PpPIEZO2* | 21 bp insertion at the first site and 345 bp + 12 bp insertion at the second CRISPR site | 21 bp insertion at the 1st sire, 49 bp of the 4th intron retained, 12 bp insertion and 55 bp deletion at the 2nd site | Stop codon in the retained intron section |
| ***aΔPp1/2-I*** | *aΔPp2-I* | *pZeo-Cas9-PpPIEZO1* | Two CRISPR targets + NHEJ* | *PpPIEZO1* | 5 bp insertion at the first site and 3 bp deletion and 38 bp insertion at the 2nd site | Same for gDNA | Frameshift and premature stop codon 58 bp after 1st mutations |
| ***aΔPp1/2-II*** | *aΔPp2-II* | *pZeo-Cas9-PpPIEZO1* | Two CRISPR targets + NHEJ* | *PpPIEZO1* | 493 bp deletion and 115 bp insertion between two CRISPR sites | 230 bp deletion and 115 bp insertion | Premature stop codon 39 bp into the insertion |

| | | | | | | | |
|---|---|---|---|---|---|---|---|
| ***gPpPIEZO1-mGFP*** | WT | *pGEM-gPP1-mGFP-Kan* | Homologous recombination-based insertion | *PpPIEZO1* | Insertion of *mGFP* CDS and Kan$^R$ cassette. | *PpPIEZO1* mRNA is fused to *mGFP* CDS | PpPIEZO1 with C-terminal mGFP tag |
| ***gPpPIEZO2-mGFP-I*** | WT | *pGEM-gPP2-mGFP-Kan* | Homologous recombination-based insertion | *PpPIEZO2* | Insertion of *mGFP* CDS and Kan$^R$ cassette. | *PpPIEZO2* mRNA is fused to *mGFP* CDS | PpPIEZO2 with C-terminal mGFP tag |
| ***gPpPIEZO2-mGFP-II*** | WT | *pGEM-gPP2-mGFP-Kan* | Homologous recombination-based insertion | *PpPIEZO2* | Insertion of *mGFP* CDS and Kan$^R$ cassette. | *PpPIEZO2* mRNA is fused to *mGFP* CDS | PpPIEZO2 with C-terminal mGFP tag |
| ***gPpPIEZO2-mGFP-III*** | WT | *pGEM-gPP2-mGFP-Kan* | Homologous recombination-based insertion | *PpPIEZO2* | Insertion of *mGFP* CDS and Kan$^R$ cassette. | *PpPIEZO2* mRNA is fused to *mGFP* CDS | PpPIEZO2 with C-terminal mGFP tag |
| ***ΔPp1/2-II + UBQ::oPpPIEZO1-mGFP-I*** | *ΔPp1/2-II* | *pTHUBI-oPpPIEZO1-mGFP + pMK-Cas9-PP108* | CRISPR + homology-directed repair | *PP108* (Pp3c20_980V3.1) | Insertion of expression cassette into a redundant locus | N/A | N/A |
| ***ΔPp1/2-II + UBQ::oPpPIEZO1-mGFP-II*** | *ΔPp1/2-II* | *pTHUBI-oPpPIEZO1-mGFP + pMK-Cas9-PP108* | CRISPR + homology-directed repair | *PP108* (Pp3c20_980V3.1) | Insertion of expression cassette into a redundant locus | N/A | N/A |
| ***ΔPp1/2-II + UBQ::oPpPIEZO2-mGFP-I*** | *ΔPp1/2-II* | *pTHUBI-oPpPIEZO2-mGFP + pMK-Cas9-PP108* | CRISPR + homology-directed repair | *PP108* (Pp3c20_980V3.1) | Insertion of expression cassette into a redundant locus | N/A | N/A |
| ***ΔPp1/2-II + UBQ::oPpPIEZO2-mGFP-II*** | *ΔPp1/2-II* | *pTHUBI-oPpPIEZO2-mGFP + pMK-Cas9-PP108* | CRISPR + homology-directed repair | *PP108* (Pp3c20_980V3.1) | Insertion of expression cassette into a redundant locus | N/A | N/A |
| ***ΔPp1/2-II + UBQ::oAtPIEZO1-mGFP-I*** | *ΔPp1/2-II* | *pTHUBI-oAtPIEZO1-mGFP + pMK-Cas9-PP108* | CRISPR + homology-directed repair | *PP108* (Pp3c20_980V3.1) | Insertion of expression cassette into a redundant locus | N/A | N/A |

| | | | | | | | |
|---|---|---|---|---|---|---|---|
| ***ΔPp1/2-II + UBQ::oAtPIEZO1-mGFP-II*** | *ΔPp1/2-II* | *pTHUBI-oAtPIEZO1-mGFP + pMK-Cas9-PP108* | CRISPR + homology-directed repair | *PP108* (Pp3c20_980V3.1) | Insertion of expression cassette into a redundant locus | N/A | N/A |
| ***PpPIEZO2 R2508K-I*** | WT | *pMH-Cas9_PP2_R2508* | CRISPR + ONAHDR** | *PpPIEZO2* | TTTATTCGGTTGCAG → TTTATTAAGTTGCAG | Same gDNA | FIRLQ → FIKLQ |
| ***PpPIEZO2 R2508K-II*** | WT | *pMH-Cas9_PP2_R2508* | CRISPR + ONAHDR** | *PpPIEZO2* | TTTATTCGGTTGCAG → TTTATTAAGTTGCAG | Same gDNA | FIRLQ → FIKLQ |
| ***PpPIEZO2 R2508H-I*** | WT | *pMH-Cas9_PP2_R2508* | CRISPR + ONAHDR** | *PpPIEZO2* | TTTATTCGGTTGCAG → TTTATTCACCTGCAG | Same gDNA | FIRLQ → FIHLQ |
| ***PpPIEZO2 R2508H-II*** | WT | *pMH-Cas9_PP2_R2508* | CRISPR + ONAHDR** | *PpPIEZO2* | TTTATTCGGTTGCAG → TTTATTCACCTGCAG | Same gDNA | FIRLQ → FIHLQ |
| ***PpPIEZO2 E2549del-I*** | WT | *pMH-Cas9_PP2_R2549* | CRISPR + ONAHDR** | *PpPIEZO2* | GAGCTAGAGGAGGGT → GAGCTCGAA---GGT | Same gDNA | ELEEG → ELE_G |
| ***PpPIEZO2 E2549del-II*** | WT | *pMH-Cas9_PP2_R2549* | CRISPR + ONAHDR** | *PpPIEZO2* | GAGCTAGAGGAGGGT → GAGCTCGAA---GGT | Same gDNA | ELEEG → ELE_G |

* Non-homologous end joining

**Oligodeoxynucleotide-assisted homology-directed repair
