## Supplemental Table 1 for "Regulation of Vacuole Morphology by PIEZO Channels in Spreading Earth Moss"

| Species | Higher Clasification | Common name of the species or the group | Protein name | Predicted Size (aa) | # of predicted TMs* | Sequence identifier | Source of the Sequence | Link |
|---|---|---|---|---|---|---|---|---|
| **Red algae** | | | | | | | | |
| *Chondrus crispus* | *Rhodophyta › Florideophyceae › Gigartinales* | Red algae - Irish moss | CcPIEZO | 2929 | 37 | XP_005712062.1 | NCBI | https://www.ncbi.nlm.nih.gov/protein/XP_005712062.1 |
| *Galdieria sulphuraria* | *Rhodophyta › Bangiophyceae › Cyanidiales* | Red algae | GsPIEZO | 2823 | 41 | XP_005703632.1 | NCBI | https://www.ncbi.nlm.nih.gov/protein/XP_005703632.1 |
| *Porphyridium purpureum* | *Rhodophyta › Bangiophyceae › Porphyridiales* | Red algae | PpPIEZO | 2789 | 40 | KAA8498933.1 | NCBI | https://www.ncbi.nlm.nih.gov/protein/KAA8498933.1 |
| ***Chlorophyta*** | | | | | | | | |
| *Auxenochlorella protothecoides* | *Viridiplantae › Chlorophyta › Trebouxiophyceae › Chlorellales* | Green algae | ApPIEZO | 2188 | 28 | XP_011395811.1 | NCBI | https://www.ncbi.nlm.nih.gov/protein/XP_011395811.1 |
| *Bathycoccus prasinos* | *Viridiplantae › Chlorophyta › Mamiellophyceae › Mamiellales* | Green algae | BpPIEZO | 2615 | 24 | XP_007513197.1 | NCBI | https://www.ncbi.nlm.nih.gov/protein/XP_007513197.1 |
| *Chlamydomonas eustigma* | *Viridiplantae › Chlorophyta › Chlorophyceae › Chlamydomonadales* | Green algae | CePIEZO | 3992 | 21 | GAX82130.1 | NCBI | https://www.ncbi.nlm.nih.gov/protein/GAX82130.1 |
| *Chlamydomonas reinhardtii* | *Viridiplantae › Chlorophyta › Chlorophyceae › Chlamydomonadales* | Green algae | CrPIEZO | 4638 | 27 | PNW79141.1 | NCBI | https://www.ncbi.nlm.nih.gov/protein/PNW79141.1 |
| *Chlorella variabilis* | *Viridiplantae › Chlorophyta › Trebouxiophyceae › Chlorellales* | Green algae | CvPIEZO | 2946 | 28 | XP_005847868.1 | NCBI | https://www.ncbi.nlm.nih.gov/protein/XP_005847868.1 |
| *Chloropicon primus* | *Viridiplantae › Chlorophyta › Chloropicophyceae › Chloropicales* | Green algae | CpPIEZO1 | 2443 | 26 | QDZ18668.1 | NCBI | https://www.ncbi.nlm.nih.gov/protein/QDZ18668.1 |
| *Chloropicon primus* | *Viridiplantae › Chlorophyta › Chloropicophyceae › Chloropicales* | Green algae | CpPIEZO2 | 2392 | 26 | QDZ17963.1 | NCBI | https://www.ncbi.nlm.nih.gov/protein/QDZ17963.1 |
| *Coccomyxa subellipsoidea* | *Viridiplantae › Chlorophyta › Trebouxiophyceae › Elliptochloris clade* | Green algae | CsPIEZO | 2782 | 26 | XP_005645135.1 | NCBI | https://www.ncbi.nlm.nih.gov/protein/XP_005645135.1 |
| *Micromonas commoda* | *Viridiplantae › Chlorophyta › Mamiellophyceae › Mamiellales* | Green algae | McPIEZOX | 4067 | 30 | XP_002499460.1 | NCBI | https://www.ncbi.nlm.nih.gov/protein/XP_002499460.1 |
| *Ostreococcus lucimarinus* | *Viridiplantae › Chlorophyta › Mamiellophyceae › Mamiellales* | Green algae | OlPIEZO | 2372 | 24 | XP_001418409.1 | NCBI | https://www.ncbi.nlm.nih.gov/protein/XP_001418409.1 |
| *Ostreococcus lucimarinus* | *Viridiplantae › Chlorophyta › Mamiellophyceae › Mamiellales* | Green algae | OlPIEZOX | 2525 | 23 | XP_001418109.1 | NCBI | https://www.ncbi.nlm.nih.gov/protein/XP_001418109.1 |
| *Ostreococcus tauri* | *Viridiplantae › Chlorophyta › Mamiellophyceae › Mamiellales* | Green algae | OtPIEZO | 2430 | 23 | XP_022839181.1 | NCBI | https://www.ncbi.nlm.nih.gov/protein/XP_022839181.1 |
| *Ostreococcus tauri* | *Viridiplantae › Chlorophyta › Mamiellophyceae › Mamiellales* | Green algae | OtPIEZOX | 2452 | 25 | XP_022839111.1 | NCBI | https://www.ncbi.nlm.nih.gov/protein/XP_022839111.1 |
| *Raphidocelis subcapitata* | *Viridiplantae › Chlorophyta › Chlorophyceae › Sphaeropleales* | Green algae | RsPIEZO | 3290 | 22 | GBF91850.1 | NCBI | https://www.ncbi.nlm.nih.gov/protein/GBF91850.1 |
| ***Charophyta*** | | | | | | | | |
| *Chara braunii* | *Viridiplantae › Streptophyta › Charophyceae › Charales* | | CbPIEZO | 2416 | 18 | GBG87247.1 | NCBI | https://www.ncbi.nlm.nih.gov/protein/GBG87247.1 |
| *Klebsormidium nitens* | *Viridiplantae › Streptophyta › Klebsormidiophyceae › Klebsormidiales* | | KnPIEZO | 2528 | 32 | GAQ83025.1 | NCBI | https://www.ncbi.nlm.nih.gov/protein/GAQ83025.1 |
| *Mesotaenium endlicherianum* | *Viridiplantae › Streptophyta › Zygnemophyceae › Zygnematales* | | MePIEZO | 3237 | 28 | ** | | https://figshare.com/articles/Genomes_of_subaerial_Zygnemat **Identified by direct search of predicted protein sequences using conserved |
| **Other land plants** | | | | | | | | |
| *Azolla filiculoides* | *Viridiplantae › Tracheophyta › Polypodiopsida › Salviniales* | Mosquito fern | AfPIEZO | 2449 | 32 | Azfi_s0043.g027082 | Fernbase | https://www.fernbase.org/tools/blast/match/show?blast_db_id=36;id=Azfi_s0043.g027082;hilite_coords=6-2449 |
| *Marchantia polymorpha* | *Viridiplantae › Marchantiophyta › Marchantiopsida › Marchantiales* | Common liverwort | MpPIEZO | 2626 | 27 | Mapoly0023s0171.1 | Phytozome | https://phytozome.jgi.doe.gov/pz/portal.html#!gene?search=1&crown=1&detail=1&method=0&searchText=transcriptid:33014158 |
| *Physcomitrium patens* | *Viridiplantae › Bryophyta › Bryopsida › Funariales* | Spreading earthmoss | PpPIEZO1 | 2573 | 32 | Pp3c9_13300V3.1 | Phytozome | https://phytozome.jgi.doe.gov/pz/portal.html#!gene?method=0&crown=1&search=1&detail=1&searchText=transcriptid:32914300 |
| *Physcomitrium patens* | *Viridiplantae › Bryophyta › Bryopsida › Funariales* | Spreading earthmoss | PpPIEZO2 | 2572 | 33 | Pp3c3_17170V3.1 | Phytozome | https://phytozome.jgi.doe.gov/pz/portal.html#!gene?method=0&crown=1&search=1&detail=1&searchText=transcriptid:32944813 |
| *Salvinia cucullata* | *Viridiplantae › Tracheophyta › Polypodiopsida › Salviniales* | Watermoss (Fern) | ScPIEZO | 2395 | 32 | Sacu_v1.1_s0023.g008885 | Fernbase | https://www.fernbase.org/tools/blast/match/show?blast_db_id=40;id=Sacu_v1.1_s0023.g008885;hilite_coords=15-2392 |
| *Selaginella moellendorffii* | *Viridiplantae › Tracheophyta › Lycopodiopsida › Selaginellales* | | SmPIEZO | 3075 | 28 | 404967 | Phytozome | https://phytozome.jgi.doe.gov/pz/portal.html#!gene?search=1&crown=1&detail=1&method=0&searchText=transcriptid:15414154 |
| *Sphagnum fallax* | *Viridiplantae › Bryophyta › Sphagnopsida › Sphagnales* | Flat-topped bogmoss | SfPIEZO1 | 2626 | 31 | Sphfalx0090s0046.1 | Phytozome | https://phytozome.jgi.doe.gov/pz/portal.html#!gene?search=1&crown=1&detail=1&method=0&searchText=transcriptid:32629378 |
| *Sphagnum fallax* | *Viridiplantae › Bryophyta › Sphagnopsida › Sphagnales* | Flat-topped bogmoss | SfPIEZO2 | 2568 | 33 | Sphfalx0044s0102.1 | Phytozome | https://phytozome.jgi.doe.gov/pz/portal.html#!gene?search=1&crown=1&detail=1&method=0&searchText=transcriptid:32619892 |
| **Basal angiosperms** | | | | | | | | |
| *Amborella trichopoda* | *Viridiplantae › Tracheophyta › Magnoliopsida › Amborellales* | | AtPIEZO | 2485 | 32 | evm_27.model.AmTr_v1.0_scaffold00025.159 | Phytozome | https://phytozome.jgi.doe.gov/pz/portal.html#!gene?search=1&crown=1&detail=1&method=0&searchText=transcriptid:31566310 |
| *Cinnamomum micranthum* | *Viridiplantae › Tracheophyta › Magnoliopsida › Laurales* | Cinnamon | CmPIEZO | 2497 | 33 | RWR76622.1 | NCBI | https://www.ncbi.nlm.nih.gov/protein/RWR76622.1?report=genbank&log$=prottop&blast_rank=1&RID=2A5XVRZ401R |
| *Nymphaea colorata* | *Viridiplantae › Tracheophyta › Magnoliopsida › Nymphaeales* | Type of water lily | NcPIEZO | 2496 | 29 | XP_031501189.1 | NCBI | https://www.ncbi.nlm.nih.gov/protein/XP_031501189.1 |
| **Monocots** | | | | | | | | |
| *Ananas comosus* | *Viridiplantae › Tracheophyta › Magnoliopsida › Poales* | Pineapple | AcPIEZO | 2251 | 27 | Aco019669.1 | Phytozome | https://phytozome.jgi.doe.gov/pz/portal.html#!gene?method=0&crown=1&search=1&detail=1&searchText=transcriptid:33052821 |
| *Brachypodium distachyon* | *Viridiplantae › Tracheophyta › Magnoliopsida › Poales* | Purple false brome | BdPIEZO | 2503 | 33 | Bradi2g13251.1 | Phytozome | https://phytozome.jgi.doe.gov/pz/portal.html#!gene?method=0&crown=1&search=1&detail=1&searchText=transcriptid:32772825 |
| *Dendrobium catenatum* | *Viridiplantae › Tracheophyta › Magnoliopsida › Asparagales* | Type of orchid | DcPIEZO | 2493 | 30 | XP_020701627.1 | NCBI | https://www.ncbi.nlm.nih.gov/protein/XP_020701627.1 |
| *Elaeis guineensis* | *Viridiplantae › Tracheophyta › Magnoliopsida › Arecales* | African oil palm | EgPIEZO | 2504 | 29 | XP_010914337.1 | NCBI | https://www.ncbi.nlm.nih.gov/protein/XP_010914337.1 |
| *Musa acuminata* | *Viridiplantae › Tracheophyta › Magnoliopsida › Zingiberales* | Species of banana | MaPIEZO | 2502 | 31 | XP_009387181.1 | NCBI | https://www.ncbi.nlm.nih.gov/protein/XP_009387181.1 |
| *Panicum hallii* | *Viridiplantae › Tracheophyta › Magnoliopsida › Poales* | Hall's panicgrass | PhPIEZO | 2501 | 31 | Pahal.H01333.1 | Phytozome | https://phytozome.jgi.doe.gov/pz/portal.html#!gene?method=0&crown=1&search=1&detail=1&searchText=transcriptid:32498593 |
| *Phoenix dactylifera* | *Viridiplantae › Tracheophyta › Magnoliopsida › Arecales* | Date palm | PdPIEZO | 2407 | 26 | XP_026663953.1 | NCBI | https://www.ncbi.nlm.nih.gov/protein/XP_026663953.1 |
| *Sorghum bicolor* | *Viridiplantae › Tracheophyta › Magnoliopsida › Poales* | Sorghum | SbPIEZO | 2579 | 30 | Sobic.003G170700.5 | Phytozome | https://phytozome.jgi.doe.gov/pz/portal.html#!gene?method=0&crown=1&search=1&detail=1&searchText=transcriptid:37912540 |
| ***Basal eudicots*** | | | | | | | | |
| *Aquilegia coerulea* | *Viridiplantae › Tracheophyta › Magnoliopsida › Ranunculales* | Colorado blue columbine | AcPIEZO1 | 2486 | 34 | PIA39355.1 | NCBI | https://www.ncbi.nlm.nih.gov/protein/PIA39355.1 |
| *Aquilegia coerulea* | *Viridiplantae › Tracheophyta › Magnoliopsida › Ranunculales* | Colorado blue columbine | AcPIEZO2 | 2406 | 29 | PIA44723.1 | NCBI | https://www.ncbi.nlm.nih.gov/protein/PIA44723.1 |
| *Macleaya cordata* | *Viridiplantae › Tracheophyta › Magnoliopsida › Ranunculales* | Five-seeded plume-poppy | McPIEZO1 | 2263 | 25 | OVA11397.1 | NCBI | https://www.ncbi.nlm.nih.gov/protein/OVA11397.1 |
| *Macleaya cordata* | *Viridiplantae › Tracheophyta › Magnoliopsida › Ranunculales* | Five-seeded plume-poppy | McPIEZO2 | 2408 | 32 | OVA04417.1 | NCBI | https://www.ncbi.nlm.nih.gov/protein/OVA04417.1 |
| *Nelumbo nucifera* | *Viridiplantae › Tracheophyta › Magnoliopsida › Proteales* | Indian lotus | NnPIEZO | 2472 | 33 | XP_010242543.1 | NCBI | https://www.ncbi.nlm.nih.gov/protein/XP_010242543.1 |
| *Papaver somniferum* | *Viridiplantae › Tracheophyta › Magnoliopsida › Ranunculales* | Opium poppy | PsPIEZO1 | 2485 | 33 | XP_026456484.1 | NCBI | https://www.ncbi.nlm.nih.gov/protein/XP_026456484.1 |
| *Papaver somniferum* | *Viridiplantae › Tracheophyta › Magnoliopsida › Ranunculales* | Opium poppy | PsPIEZO2 | 2480 | 35 | XP_026450010.1 | NCBI | https://www.ncbi.nlm.nih.gov/protein/XP_026450010.1 |
| ***Superasterids*** | | | | | | | | |
| *Actinidia chinensis* | *Viridiplantae › Tracheophyta › Magnoliopsida › Ericales* | Golden kiwifruit | AcPIEZO1 | 2510 | 31 | PSR94793.1 | NCBI | https://www.ncbi.nlm.nih.gov/protein/PSR94793.1 |
| *Actinidia chinensis* | *Viridiplantae › Tracheophyta › Magnoliopsida › Ericales* | Golden kiwifruit | AcPIEZO2 | 2473 | 28 | PSS10163.1 | NCBI | https://www.ncbi.nlm.nih.gov/protein/PSS10163.1 |
| *Amaranthus hypochondriacus* | *Viridiplantae › Tracheophyta › Magnoliopsida ›* | Prince's-feather | AhPIEZO1 | 2474 | 32 | AHYPO_009829-RA | Phytozome | https://phytozome.jgi.doe.gov/pz/portal.html#!gene?method=0&crown=1&search=1&detail=1&searchText=transcriptid:32826545 |
| *Amaranthus hypochondriacus* | *Viridiplantae › Tracheophyta › Magnoliopsida › Caryophyllales* | Prince's-feather | AhPIEZO2 | 2163 | 23 | AHYPO_008786-RA | Phytozome | https://phytozome.jgi.doe.gov/pz/portal.html#!gene?method=0&crown=1&search=1&detail=1&searchText=transcriptid:32839695 |
| *Beta vulgaris* | *Viridiplantae › Tracheophyta › Magnoliopsida › Caryophyllales* | Beet | BvPIEZO1 | 2502 | 33 | XP_010696335.1 | NCBI | https://www.ncbi.nlm.nih.gov/protein/XP_010696335.1 |
| *Camellia sinensis* | *Viridiplantae › Tracheophyta › Magnoliopsida › Ericales* | Tea plant | CsPIEZO1a | 2399 | 27 | THG15462.1 | NCBI | https://www.ncbi.nlm.nih.gov/protein/THG15462.1 |
| *Camellia sinensis* | *Viridiplantae › Tracheophyta › Magnoliopsida › Ericales* | Tea plant | CsPIEZO1b | 2366 | 25 | THG04468.1 | NCBI | https://www.ncbi.nlm.nih.gov/protein/THG04468.1 |
| *Coffea arabica* | *Viridiplantae › Tracheophyta › Magnoliopsida › Gentianales* | Arabica coffee | CaPIEZO1 | 2475 | 31 | XP_027116044.1 | NCBI | https://www.ncbi.nlm.nih.gov/protein/XP_027116044.1 |
| *Coffea arabica* | *Viridiplantae › Tracheophyta › Magnoliopsida › Gentianales* | Arabica coffee | CaPIEZO2 | 2468 | 29 | XP_027114534.1 | NCBI | https://www.ncbi.nlm.nih.gov/protein/XP_027114534.1 |
| *Daucus carota* | *Viridiplantae › Tracheophyta › Magnoliopsida › Apiales* | Wild carrot | DcPIEZO1a | 2479 | 35 | XP_017241506.1 | NCBI | https://www.ncbi.nlm.nih.gov/protein/XP_017241506.1 |
| *Daucus carota* | *Viridiplantae › Tracheophyta › Magnoliopsida › Apiales* | Wild carrot | DcPIEZO1b | 2473 | 33 | XP_017252549.1 | NCBI | https://www.ncbi.nlm.nih.gov/protein/XP_017252549.1 |
| *Daucus carota* | *Viridiplantae › Tracheophyta › Magnoliopsida › Apiales* | Wild carrot | DcPIEZO2 | 2475 | 33 | XP_017255589.1 | NCBI | https://www.ncbi.nlm.nih.gov/protein/XP_017255589.1 |
| *Helianthus annuus* | *Viridiplantae › Tracheophyta › Magnoliopsida › Asterales* | Common sunflower | HaPIEZO1a | 2480 | 32 | XP_022010653.1 | NCBI | https://www.ncbi.nlm.nih.gov/protein/XP_022010653.1 |
| *Helianthus annuus* | *Viridiplantae › Tracheophyta › Magnoliopsida › Asterales* | Common sunflower | HaPIEZO1b | 2441 | 35 | XP_022035444.1 | NCBI | https://www.ncbi.nlm.nih.gov/protein/XP_022035444.1 |
| *Lactuca sativa* | *Viridiplantae › Tracheophyta › Magnoliopsida › Asterales* | Lactuca sativa | LsPIEZO1a | 2457 | 25 | XP_023738449.1 | NCBI | https://www.ncbi.nlm.nih.gov/protein/XP_023738449.1 |
| *Lactuca sativa* | *Viridiplantae › Tracheophyta › Magnoliopsida › Asterales* | Lactuca sativa | LsPIEZO1b | 2424 | 37 | XP_023749583.1 | NCBI | https://www.ncbi.nlm.nih.gov/protein/XP_023749583.1 |
| *Nicotiana sylvestris* | *Viridiplantae › Tracheophyta › Magnoliopsida › Solanales* | Woodland tobacco | NsPIEZO1 | 2473 | 31 | XP_009770604.1 | NCBI | https://www.ncbi.nlm.nih.gov/protein/XP_009770604.1 |
| *Olea europaea* | *Viridiplantae › Tracheophyta › Magnoliopsida › Lamiales* | Olive | OePIEZO1 | 2475 | 33 | XP_022847249.1 | NCBI | https://www.ncbi.nlm.nih.gov/protein/XP_022847249.1 |
| *Sesamum indicum* | *Viridiplantae › Tracheophyta › Magnoliopsida › Lamiales* | Sesame | SiPIEZO1a | 2480 | 33 | XP_020550018.1 | NCBI | https://www.ncbi.nlm.nih.gov/protein/XP_020550018.1 |
| *Sesamum indicum* | *Viridiplantae › Tracheophyta › Magnoliopsida › Lamiales* | Sesame | SiPIEZO1b | 2475 | 33 | XP_011082533.1 | NCBI | https://www.ncbi.nlm.nih.gov/protein/XP_011082533.1 |
| *Solanum lycopersicum* | *Viridiplantae › Tracheophyta › Magnoliopsida › Solanales* | Tomato | SlPIEZO1 | 2473 | 31 | XP_010326620.1 | NCBI | https://www.ncbi.nlm.nih.gov/protein/XP_010326620.1 |
| *Striga asiatica* | *Viridiplantae › Tracheophyta › Magnoliopsida › Lamiales* | Asiatic witchweed | SaPIEZO1a | 2468 | 30 | GER26520.1 | NCBI | https://www.ncbi.nlm.nih.gov/protein/GER26520.1 |
| *Striga asiatica* | *Viridiplantae › Tracheophyta › Magnoliopsida › Lamiales* | Asiatic witchweed | SaPIEZO1b | 2615 | 35 | GER32974.1 | NCBI | https://www.ncbi.nlm.nih.gov/protein/GER32974.1 |
| **Superrosids** | | | | | | | | |
| *Arabidopsis thaliana* | *Viridiplantae › Tracheophyta › Magnoliopsida › Brassicales* | Thale cress | AtPIEZO1 | 2485 | 31 | NP_001323909.1 | NCBI | https://www.ncbi.nlm.nih.gov/protein/NP_001323909.1 |
| *Boechera stricta* | *Viridiplantae › Tracheophyta › Magnoliopsida › Brassicales* | Drummond's rockcress | BcPIEZO1 | 2482 | 31 | Bostr.13208s0012.1 | Phytozome | https://phytozome.jgi.doe.gov/pz/portal.html#!gene?method=0&crown=1&search=1&detail=1&searchText=transcriptid:30674854 |
| *Capsella rubella* | *Viridiplantae › Tracheophyta › Magnoliopsida › Brassicales* | Pink shepherd's-purse | CePIEZO1 | 2485 | 31 | Carubv10022493m | Phytozome | https://phytozome.jgi.doe.gov/pz/portal.html#!gene?method=0&crown=1&search=1&detail=1&searchText=transcriptid:20901550 |
| *Cephalotus follicularis* | *Viridiplantae › Tracheophyta › Magnoliopsida › Oxalidales* | Albany pitcher plant | CfPIEZO1 | 2487 | 30 | GAV80465.1 | NCBI | https://www.ncbi.nlm.nih.gov/protein/GAV80465.1 |
| *Citrus clementina* | *Viridiplantae › Tracheophyta › Magnoliopsida › Sapindales* | Clementine | CcPIEZO1 | 2095 | 24 | Ciclev10018445m | Phytozome | https://phytozome.jgi.doe.gov/pz/portal.html#!gene?method=0&crown=1&search=1&detail=1&searchText=transcriptid:20807378 |
| *Cucumis sativus* | *Viridiplantae › Tracheophyta › Magnoliopsida › Cucurbitales* | Cucumber | CsPIEZO1 | 2330 | 30 | Cucsa.084850.1 | Phytozome | https://phytozome.jgi.doe.gov/pz/portal.html#!gene?method=0&crown=1&search=1&detail=1&searchText=transcriptid:16956604 |
| *Cucurbita maxima* | *Viridiplantae › Tracheophyta › Magnoliopsida › Cucurbitales* | Squash | CmPIEZO1a | 2475 | 33 | XP_022992489.1 | NCBI | https://www.ncbi.nlm.nih.gov/protein/XP_022992489.1 |
| *Cucurbita maxima* | *Viridiplantae › Tracheophyta › Magnoliopsida › Cucurbitales* | Squash | CmPIEZO1b | 2479 | 33 | XP_023004422.1 | NCBI | https://www.ncbi.nlm.nih.gov/protein/XP_023004422.1 |
| *Eucalyptus grandis* | *Viridiplantae › Tracheophyta › Magnoliopsida › Myrtales* | Flooded gum | EgPIEZO1 | 2481 | 33 | Eucgr.H01545.1 | Phytozome | https://phytozome.jgi.doe.gov/pz/portal.html#!gene?method=0&crown=1&search=1&detail=1&searchText=transcriptid:32043157 |
| *Eucalyptus grandis* | *Viridiplantae › Tracheophyta › Magnoliopsida › Myrtales* | Flooded gum | EgPIEZO2 | 2358 | 30 | Eucgr.B02004.1 | Phytozome | https://phytozome.jgi.doe.gov/pz/portal.html#!gene?method=0&crown=1&search=1&detail=1&searchText=transcriptid:32061515 |
| *Fragaria vesca* | *Viridiplantae › Tracheophyta › Magnoliopsida › Rosales* | Wild strawberry | FvPIEZO1 | 2185 | 22 | mrna27732.1-v1.0-hybrid | Phytozome | https://phytozome.jgi.doe.gov/pz/portal.html#!gene?method=0&crown=1&search=1&detail=1&searchText=transcriptid:27263409 |
| *Glycine max* | *Viridiplantae › Tracheophyta › Magnoliopsida › Fabales* | Soybean | GmPIEZO1a | 2483 | 31 | Glyma.10G023700.1 | Phytozome | https://phytozome.jgi.doe.gov/pz/portal.html#!gene?method=0&crown=1&search=1&detail=1&searchText=transcriptid:30476151 |
| *Glycine max* | *Viridiplantae › Tracheophyta › Magnoliopsida › Fabales* | Soybean | GmPIEZO1b | 2482 | 33 | Glyma.02G150200.1 | Phytozome | https://phytozome.jgi.doe.gov/pz/portal.html#!gene?method=0&crown=1&search=1&detail=1&searchText=transcriptid:30511069 |
| *Glycine max* | *Viridiplantae › Tracheophyta › Magnoliopsida › Fabales* | Soybean | GmPIEZO2 | 2138 | 23 | Glyma.19G238300.1 | Phytozome | https://phytozome.jgi.doe.gov/pz/portal.html#!gene?method=0&crown=1&search=1&detail=1&searchText=transcriptid:30511886 |
| *Gossypium raimondii* | *Viridiplantae › Tracheophyta › Magnoliopsida › Malvales* | Species of cotton | GrPIEZO1a | 2487 | 31 | Gorai.004G215100.1 | Phytozome | https://phytozome.jgi.doe.gov/pz/portal.html#!gene?method=0&crown=1&search=1&detail=1&searchText=transcriptid:26777611 |
| *Gossypium raimondii* | *Viridiplantae › Tracheophyta › Magnoliopsida › Malvales* | Species of cotton | GrPIEZO1b | 2476 | 31 | Gorai.007G133300.1 | Phytozome | https://phytozome.jgi.doe.gov/pz/portal.html#!gene?method=0&crown=1&search=1&detail=1&searchText=transcriptid:26781839 |
| *Gossypium raimondii* | *Viridiplantae › Tracheophyta › Magnoliopsida › Malvales* | Species of cotton | GrPIEZO2 | 2466 | 30 | Gorai.013G123400.1 | Phytozome | https://phytozome.jgi.doe.gov/pz/portal.html#!gene?method=0&crown=1&search=1&detail=1&searchText=transcriptid:26786547 |
| *Herrania umbratica* | *Viridiplantae › Tracheophyta › Magnoliopsida › Malvales* | | HuPIEZO1 | 2483 | 32 | XP_021280440.1 | NCBI | https://www.ncbi.nlm.nih.gov/protein/XP_021280440.1 |
| *Herrania umbratica* | *Viridiplantae › Tracheophyta › Magnoliopsida › Malvales* | | HuPIEZO2 | 2470 | 31 | XP_021283331.1 | NCBI | https://www.ncbi.nlm.nih.gov/protein/XP_021283331.1 |
| *Juglans regia* | *Viridiplantae › Tracheophyta › Magnoliopsida › Fagales* | English walnut | JrPIEZO1 | 2486 | 30 | XP_018847450.1 | NCBI | https://www.ncbi.nlm.nih.gov/protein/XP_018847450.1 |
| *Kalanchoe fedtschenkoi* | *Viridiplantae › Tracheophyta › Magnoliopsida › Saxifragales* | Widow's-thrill | KfPIEZO1 | 2478 | 31 | Kaladp0030s0103.1 | Phytozome | https://phytozome.jgi.doe.gov/pz/portal.html#!gene?method=0&crown=1&search=1&detail=1&searchText=transcriptid:35750950 |
| *Kalanchoe fedtschenkoi* | *Viridiplantae › Tracheophyta › Magnoliopsida › Saxifragales* | Widow's-thrill | KfPIEZO2 | 2451 | 31 | Kaladp0027s0078.1 | Phytozome | https://phytozome.jgi.doe.gov/pz/portal.html#!gene?method=0&crown=1&search=1&detail=1&searchText=transcriptid:35781550 |
| *Manihot esculenta* | *Viridiplantae › Tracheophyta › Magnoliopsida › Malpighiales* | Cassava | MePIEZO1 | 2485 | 32 | Manes.05G071700.1 | Phytozome | https://phytozome.jgi.doe.gov/pz/portal.html#!gene?method=0&crown=1&search=1&detail=1&searchText=transcriptid:32337353 |
| *Manihot esculenta* | *Viridiplantae › Tracheophyta › Magnoliopsida › Malpighiales* | Cassava | MePIEZO2 | 2499 | 29 | Manes.09G162700.1 | Phytozome | https://phytozome.jgi.doe.gov/pz/portal.html#!gene?method=0&crown=1&search=1&detail=1&searchText=transcriptid:32342087 |
| *Medicago truncatula* | *Viridiplantae › Tracheophyta › Magnoliopsida › Fabales* | Barrelclover | MtPIEZO1 | 2329 | 28 | Medtr1g089760.1 | Phytozome | https://phytozome.jgi.doe.gov/pz/portal.html#!gene?method=0&crown=1&search=1&detail=1&searchText=transcriptid:31099742 |
| *Medicago truncatula* | *Viridiplantae › Tracheophyta › Magnoliopsida › Fabales* | Barrelclover | MtPIEZO2 | 2462 | 30 | Medtr7g113930.1 | Phytozome | https://phytozome.jgi.doe.gov/pz/portal.html#!gene?method=0&crown=1&search=1&detail=1&searchText=transcriptid:31082655 |
| *Pistacia vera* | *Viridiplantae › Tracheophyta › Magnoliopsida › Sapindales* | Pistachio | PvPIEZO1 | 2447 | 33 | XP_031279982.1 | NCBI | https://www.ncbi.nlm.nih.gov/protein/XP_031279982.1 |
| *Populus trichocarpa* | *Viridiplantae › Tracheophyta › Magnoliopsida › Malpighiales* | Black cottonwood | PtPIEZO1 | 2482 | 31 | Potri.014G137200.1 | Phytozome | https://phytozome.jgi.doe.gov/pz/portal.html#!gene?method=0&crown=1&search=1&detail=1&searchText=transcriptid:27032594 |
| *Populus trichocarpa* | *Viridiplantae › Tracheophyta › Magnoliopsida › Malpighiales* | Black cottonwood | PtPIEZO2 | 2366 | 26 | Potri.005G037700.1 | Phytozome | https://phytozome.jgi.doe.gov/pz/portal.html#!gene?method=0&crown=1&search=1&detail=1&searchText=transcriptid:27027537 |
| *Prunus persica* | *Viridiplantae › Tracheophyta › Magnoliopsida › Rosales* | Peach | PpPIEZO1 | 2485 | 34 | Prupe.8G248000.1 | Phytozome | https://phytozome.jgi.doe.gov/pz/portal.html#!gene?method=0&crown=1&search=1&detail=1&searchText=transcriptid:32081222 |
| *Punica granatum* | *Viridiplantae › Tracheophyta › Magnoliopsida › Myrtales* | Pomegranate | PgPIEZO1 | 2440 | 32 | XP_031401099.1 | NCBI | https://www.ncbi.nlm.nih.gov/protein/XP_031401099.1 |
| *Punica granatum* | *Viridiplantae › Tracheophyta › Magnoliopsida › Myrtales* | Pomegranate | PgPIEZO2 | 2470 | 31 | XP_031403533.1 | NCBI | https://www.ncbi.nlm.nih.gov/protein/XP_031403533.1 |
| *Quercus lobata* | *Viridiplantae › Tracheophyta › Magnoliopsida › Fagales* | Valley oak | QlPIEZO1 | 2488 | 29 | XP_030947959.1 | NCBI | https://www.ncbi.nlm.nih.gov/protein/XP_030947959.1 |
| *Quercus suber* | *Viridiplantae › Tracheophyta › Magnoliopsida › Fagales* | Cork oak | QsPIEZO1 | 2488 | 29 | XP_023880259.1 | NCBI | https://www.ncbi.nlm.nih.gov/protein/XP_023880259.1 |
| *Quercus suber* | *Viridiplantae › Tracheophyta › Magnoliopsida › Fagales* | Cork oak | QsPIEZO2 | 2492 | 33 | XP_023908417.1 | NCBI | https://www.ncbi.nlm.nih.gov/protein/XP_023908417.1 |
| *Ricinus communis* | *Viridiplantae › Tracheophyta › Magnoliopsida › Malpighiales* | Ricinus | RcPIEZO1 | 2254 | 25 | 29703.m001534 | Phytozome | https://phytozome.jgi.doe.gov/pz/portal.html#!gene?method=0&crown=1&search=1&detail=1&searchText=transcriptid:16806230 |
| *Ricinus communis* | *Viridiplantae › Tracheophyta › Magnoliopsida › Malpighiales* | Ricinus | RcPIEZO2 | 2361 | 29 | 30128.m008722 | Phytozome | https://phytozome.jgi.doe.gov/pz/portal.html#!gene?method=0&crown=1&search=1&detail=1&searchText=transcriptid:16818311 |
| *Tarenaya hassleriana* | *Viridiplantae › Tracheophyta › Magnoliopsida › Brassicales* | Spider flower | ThPIEZO1 | 2473 | 29 | XP_010524005.1 | NCBI | https://www.ncbi.nlm.nih.gov/protein/XP_010524005.1 |
| *Theobroma cacao* | *Viridiplantae › Tracheophyta › Magnoliopsida › Malvales* | Cacao tree | TcPIEZO1 | 2501 | 32 | Thecc1EG004478t1 | Phytozome | https://phytozome.jgi.doe.gov/pz/portal.html#!gene?method=0&crown=1&search=1&detail=1&searchText=transcriptid:27437166 |
| *Theobroma cacao* | *Viridiplantae › Tracheophyta › Magnoliopsida › Malvales* | Cacao tree | TcPIEZO2 | 2463 | 30 | Thecc1EG026520t1 | Phytozome | https://phytozome.jgi.doe.gov/pz/portal.html#!gene?method=0&crown=1&search=1&detail=1&searchText=transcriptid:27424808 |
| *Vitis vinifera* | *Viridiplantae › Tracheophyta › Magnoliopsida › Vitales* | Common Grape Vine | VvPIEZO1 | 2409 | 29 | GSVIVT01011059001 | Phytozome | https://phytozome.jgi.doe.gov/pz/portal.html#!gene?method=0&crown=1&search=1&detail=1&searchText=transcriptid:17823040 |
| *Vitis vinifera* | *Viridiplantae › Tracheophyta › Magnoliopsida › Vitales* | Common Grape Vine | VvPIEZO2 | 2376 | 29 | GSVIVT01022038001 | Phytozome | https://phytozome.jgi.doe.gov/pz/portal.html#!gene?method=0&crown=1&search=1&detail=1&searchText=transcriptid:17830971 |
| ***Placozoa, Sponges, Cnidaria and Ctenophora*** | | | | | | | | |
| *Amphimedon queenslandica* | *Metazoa › Porifera › Haplosclerida* | Species of sponge | AqPIEZO | 2317 | 29 | XP_019849310.1 | NCBI | https://www.ncbi.nlm.nih.gov/protein/XP_019849310.1 |
| *Beroe abyssicola* | *Metazoa › Ctenophora › Beroida* | Cigar comb jelly | BaPIEZO | 2345 | 32 | AQX17753.1 | NCBI | https://www.ncbi.nlm.nih.gov/protein/AQX17753.1 |
| *Bolinopsis infundibulum* | *Metazoa › Ctenophora › Lobata* | Common northern comb jelly | BiPIEZO | 2466 | 36 | AQX17755.1 | NCBI | https://www.ncbi.nlm.nih.gov/protein/AQX17755.1 |
| *Dendronephthya gigantea* | *Metazoa › Cnidaria › Alcyonacea* | Species of soft coral | DgPIEZO | 2294 | 33 | XP_028392641.1 | NCBI | https://www.ncbi.nlm.nih.gov/protein/XP_028392641.1 |
| *Pocillopora damicornis* | *Metazoa › Cnidaria › Scleractinia* | Cauliflower coral | PdPIEZO | 2319 | 29 | XP_027057832.1 | NCBI | https://www.ncbi.nlm.nih.gov/protein/XP_027057832.1 |
| *Trichoplax sp.* | *Metazoa › Placozoa › Trichoplacidae* | | TsPIEZO | 2443 | 36 | RDD46920.1 | NCBI | https://www.ncbi.nlm.nih.gov/protein/RDD46920.1 |
| **Invertebrates** | | | | | | | | |
| *Caenorhabditis elegans* | *Metazoa › Nematoda › Rhabditida* | Nematode | CePIEZO | 2402 | 33 | NP_001293979.1 | NCBI | https://www.ncbi.nlm.nih.gov/protein/NP_001293979.1 |
| *Octopus vulgaris* | *Metazoa › Mollusca › Octopoda* | Common octopus | OvPIEZO | 2429 | 38 | XP_029647891.1 | NCBI | https://www.ncbi.nlm.nih.gov/protein/XP_029647891.1 |
| *Pomacea canaliculata* | *Metazoa › Mollusca › Gastropoda* | Golden apple snail | PcPIEZO | 2440 | 36 | XP_025076542.1 | NCBI | https://www.ncbi.nlm.nih.gov/protein/XP_025076542.1 |
| *Priapulus caudatus* | *Metazoa › Priapulida › Priapulimorpha* | Species of penis worms | PcPIEZO | 2300 | 32 | XP_014677335.1 | NCBI | https://www.ncbi.nlm.nih.gov/protein/XP_014677335.1 |
| *Ramazzottius varieornatus* | *Metazoa › Tardigrada › Parachela* | Specie of water bear | RvPIEZO | 2497 | 34 | GAU96232.1 | NCBI | https://www.ncbi.nlm.nih.gov/protein/GAU96232.1 |
| **Hexapoda** | | | | | | | | |
| *Aedes aegypti* | *Metazoa › Arthropoda › Diptera* | Yellow fever mosquito | AePIEZO | 3096 | 31 | XP_021703538.1 | NCBI | https://www.ncbi.nlm.nih.gov/protein/XP_021703538.1 |
| *Apis mellifera* | *Metazoa › Arthropoda › Hymenoptera* | Western honey bee | AmPIEZO | 2520 | 34 | XP_006560440.2 | NCBI | https://www.ncbi.nlm.nih.gov/protein/XP_006560440.2 |
| *Cimex lectularius* | *Metazoa › Arthropoda › Hemiptera* | Bed bugs | ClPIEZO | 2569 | 35 | XP_024082956.1 | NCBI | https://www.ncbi.nlm.nih.gov/protein/XP_024082956.1 |
| *Drosophila hydei* | *Metazoa › Arthropoda › Diptera* | Species of fruit fly | DhPIEZO1 | 2723 | 36 | XP_030079259.1 | NCBI | https://www.ncbi.nlm.nih.gov/protein/XP_030079259.1 |
| *Drosophila hydei* | *Metazoa › Arthropoda › Diptera* | Species of fruit fly | DhPIEZO1 | 2302 | 31 | XP_030081590.1 | NCBI | https://www.ncbi.nlm.nih.gov/protein/XP_030081590.1 |
| *Drosophila melanogaster* | *Metazoa › Arthropoda › Diptera* | Common fruit fly | DmPIEZO1 | 2533 | 39 | NP_001260228 | NCBI | https://www.ncbi.nlm.nih.gov/protein/NP_001260228 |
| *Drosophila melanogaster* | *Metazoa › Arthropoda › Diptera* | Common fruit fly | DmPIEZO2 | 2326 | 38 | NP_001303493.1 | NCBI | https://www.ncbi.nlm.nih.gov/protein/NP_001303493.1 |
| *Frankliniella occidentalis* | *Metazoa › Arthropoda › Thysanoptera* | Western flower thrips | FaPIEZO | 2755 | 31 | XP_026289758.1 | NCBI | https://www.ncbi.nlm.nih.gov/protein/XP_026289758.1 |
| *Lucilia cuprina* | *Metazoa › Arthropoda › Diptera* | Australian sheep blowfly | LcPIEZO | 2524 | 36 | XP_023298568.1 | NCBI | https://www.ncbi.nlm.nih.gov/protein/XP_023298568.1 |
| *Orchesella cincta* | *Metazoa › Arthropoda › Collembola* | Species of springtail | OcPIEZO | 2236 | 25 | ODM97774.1 | NCBI | https://www.ncbi.nlm.nih.gov/protein/ODM97774.1 |
| *Pieris rapae* | *Metazoa › Arthropoda › Lepidoptera* | Cabbage white buttfly | PrPIEZO | 2536 | 35 | XP_022128296.1 | NCBI | https://www.ncbi.nlm.nih.gov/protein/XP_022128296.1 |
| *Schistocerca gregaria* | *Metazoa › Arthropoda › Orthoptera* | Desert locust | ScPIEZO | 2528 | 31 | QGX90119.1 | NCBI | https://www.ncbi.nlm.nih.gov/protein/QGX90119.1 |
| *Stomoxys calcitrans* | *Metazoa › Arthropoda › Diptera* | Stable fly | ScPIEZO | 2677 | 36 | XP_013104293.1 | NCBI | https://www.ncbi.nlm.nih.gov/protein/XP_013104293.1 |
| *Trichoplusia ni* | *Metazoa › Arthropoda › Lepidoptera* | Cabbage looper | TnPIEZO | 2569 | 31 | XP_026726942.1 | NCBI | https://www.ncbi.nlm.nih.gov/protein/XP_026726942.1 |
| **Hemichordata, Echinodermata and Chordata** | | | | | | | | |
| *Acanthaster planci* | *Metazoa › Echinodermata › Valvatida* | Crown-of-thorns starfish | ApPIEZO | 2530 | 39 | XP_022080337.1 | NCBI | https://www.ncbi.nlm.nih.gov/protein/XP_022080337.1 |
| *Branchiostoma belcheri* | *Metazoa › Chordata › Amphioxiformes* | Species of lancelets | BbPIEZO | 2827 | 33 | XP_019626141.1 | NCBI | https://www.ncbi.nlm.nih.gov/protein/XP_019626141.1 |
| *Ciona intestinalis* | *Metazoa › Chordata › Enterogona* | Sea vase | CiPIEZO | 2342 | 31 | XP_026690886.1 | NCBI | https://www.ncbi.nlm.nih.gov/protein/XP_026690886.1 |
| *Saccoglossus kowalevskii* | *Metazoa › Hemichordata › Enteropneusta* | Species of acorn worms | SkPIEZO | 2411 | 35 | XP_006818270.1 | NCBI | https://www.ncbi.nlm.nih.gov/protein/XP_006818270.1 |
| *Strongylocentrotus purpuratus* | *Metazoa › Echinodermata › Echinoida* | Pacific purple sea urchin | SpPIEZO | 2467 | 35 | XP_030831611.1 | NCBI | https://www.ncbi.nlm.nih.gov/protein/XP_030831611.1 |
| **Vertebrates** | | | | | | | | |
| *Callorhinchus milii* | *Metazoa › Chordata › Chondrichthyes › Chimaeriformes* | Australian ghostshark | CmPIEZO1 | 2512 | 33 | XP_007887509.1 | NCBI | https://www.ncbi.nlm.nih.gov/protein/XP_007887509.1 |
| *Callorhinchus milii* | *Metazoa › Chordata › Chondrichthyes › Chimaeriformes* | Australian ghostshark | CmPIEZO2 | 2790 | 35 | XP_007895527.1 | NCBI | https://www.ncbi.nlm.nih.gov/protein/XP_007895527.1 |
| *Callorhinchus milii* | *Metazoa › Chordata › Chondrichthyes › Chimaeriformes* | Australian ghostshark | CmPIEZO3 | 2701 | 33 | XP_007883870.1 | NCBI | https://www.ncbi.nlm.nih.gov/protein/XP_007883870.1 |
| *Canis lupus familiaris* | *Metazoa › Chordata › Mammalia › Carnivora* | Dog | ClfPIEZO1 | 2533 | 33 | XP_022274796.1 | NCBI | https://www.ncbi.nlm.nih.gov/protein/XP_022274796.1 |
| *Canis lupus familiaris* | *Metazoa › Chordata › Mammalia › Carnivora* | Dog | ClfPIEZO2 | 2810 | 35 | XP_022277436.1 | NCBI | https://www.ncbi.nlm.nih.gov/protein/XP_022277436.1 |
| *Crocodylus porosus* | *Metazoa › Chordata › Reptilia › Crocodilia* | Saltwater crocodile | CpPIEZO1 | 2513 | 31 | XP_019390999.1 | NCBI | https://www.ncbi.nlm.nih.gov/protein/XP_019390999.1 |
| *Crocodylus porosus* | *Metazoa › Chordata › Reptilia › Crocodilia* | Saltwater crocodile | CpPIEZO2 | 2778 | 38 | XP_019401417.1 | NCBI | https://www.ncbi.nlm.nih.gov/protein/XP_019401417.1 |
| *Crocodylus porosus* | *Metazoa › Chordata › Reptilia › Crocodilia* | Saltwater crocodile | CpPIEZO3 | 2601 | 31 | XP_019403736.1 | NCBI | https://www.ncbi.nlm.nih.gov/protein/XP_019403736.1 |
| *Danio rerio* | *Metazoa › Chordata › Actinopterygii › Cypriniformes* | Zebrafish | DrPIEZO1 | 2538 | 30 | XP_696355.4 | NCBI | https://www.ncbi.nlm.nih.gov/protein/XP_696355.4 |
| *Danio rerio* | *Metazoa › Chordata › Actinopterygii › Cypriniformes* | Zebrafish | DrPIEZO2a | 2752 | 34 | XP_021325930.1 | NCBI | https://www.ncbi.nlm.nih.gov/protein/XP_021325930.1 |
| *Danio rerio* | *Metazoa › Chordata › Actinopterygii › Cypriniformes* | Zebrafish | DrPIEZO2b | 3005 | 34 | XP_021323945.1 | NCBI | https://www.ncbi.nlm.nih.gov/protein/XP_021323945.1 |
| *Danio rerio* | *Metazoa › Chordata › Actinopterygii › Cypriniformes* | Zebrafish | DrPIEZO3 | 2634 | 23 | XP_021334475.1 | NCBI | https://www.ncbi.nlm.nih.gov/protein/XP_021334475.1 |
| *Dromaius novaehollandiae* | *Metazoa › Chordata › Aves › Casuariiformes* | Emu | DnPIEZO1 | 2708 | 35 | XP_025965764.1 | NCBI | https://www.ncbi.nlm.nih.gov/protein/XP_025965764.1 |
| *Dromaius novaehollandiae* | *Metazoa › Chordata › Aves › Casuariiformes* | Emu | DnPIEZO2 | 2774 | 36 | XP_025950108.1 | NCBI | https://www.ncbi.nlm.nih.gov/protein/XP_025950108.1 |
| *Falco peregrinus* | *Metazoa › Chordata › Aves › Falconiformes* | Peregrine falcon | FpPIEZO1 | 2489 | 35 | XP_027639747.1 | NCBI | https://www.ncbi.nlm.nih.gov/protein/XP_027639747.1 |

| | | | | | | | | |
|---|---|---|---|---|---|---|---|---|
| *Falco peregrinus* | *Metazoa › Chordata › Aves › Falconiformes* | Peregrine falcon | **FpPIEZO2** | **2792** | **34** | XP_027647999.1 | NCBI | https://www.ncbi.nlm.nih.gov/protein/XP_027647999.1 |
| *Gallus gallus* | *Metazoa › Chordata › Aves › Galliformes* | Gallus gallus | **GgPIEZO1** | **2492** | **34** | XP_015148150.1 | NCBI | https://www.ncbi.nlm.nih.gov/protein/XP_015148150.1 |
| *Gallus gallus* | *Metazoa › Chordata › Aves › Galliformes* | Gallus gallus | **GgPIEZO2** | **2796** | **35** | XP_025003586.1 | NCBI | https://www.ncbi.nlm.nih.gov/protein/XP_025003586.1 |
| *Homo sapiens* | *Metazoa › Chordata › Mammalia › Primates* | Human | **HsPIEZO1** | **2521** | **31** | NP_001136336.2 | NCBI | https://www.ncbi.nlm.nih.gov/protein/NP_001136336.2/ |
| *Homo sapiens* | *Metazoa › Chordata › Mammalia › Primates* | Human | **HsPIEZO2** | **2752** | **35** | NP_071351.2 | NCBI | https://www.ncbi.nlm.nih.gov/protein/NP_071351.2 |
| *Latimeria chalumnae* | *Metazoa › Chordata › Coelacanthiformes* | West Indian Ocean coelacanth | **LcPIEZO1** | **2614** | **33** | XP_014347544.1 | NCBI | https://www.ncbi.nlm.nih.gov/protein/XP_014347544.1 |
| *Latimeria chalumnae* | *Metazoa › Chordata › Coelacanthiformes* | West Indian Ocean coelacanth | **LcPIEZO2** | **2596** | **35** | XP_014343966.1 | NCBI | https://www.ncbi.nlm.nih.gov/protein/XP_014343966.1 |
| *Latimeria chalumnae* | *Metazoa › Chordata › Coelacanthiformes* | West Indian Ocean coelacanth | **LcPIEZO3** | **2665** | **28** | XP_014342463.1 | NCBI | https://www.ncbi.nlm.nih.gov/protein/XP_014342463.1 |
| *Microcaecilia unicolor* | *Metazoa › Chordata › Amphibia › Gymnophiona* | Species of amphibian | **MuPIEZO1** | **2518** | **36** | XP_030060451.1 | NCBI | https://www.ncbi.nlm.nih.gov/protein/XP_030060451.1 |
| *Microcaecilia unicolor* | *Metazoa › Chordata › Amphibia › Gymnophiona* | Species of amphibian | **MuPIEZO2** | **2772** | **36** | XP_030042384.1 | NCBI | https://www.ncbi.nlm.nih.gov/protein/XP_030042384.1 |
| *Microcaecilia unicolor* | *Metazoa › Chordata › Amphibia › Gymnophiona* | Species of amphibian | **MuPIEZO3** | **2649** | **29** | XP_030069316.1 | NCBI | https://www.ncbi.nlm.nih.gov/protein/XP_030069316.1 |
| *Monodelphis domestica* | *Metazoa › Chordata › Mammalia › Didelphimorphia* | Gray short-tailed opossum | **MdPIEZO1** | **2538** | **33** | XP_007477361.1 | NCBI | https://www.ncbi.nlm.nih.gov/protein/XP_007477361.1 |
| *Monodelphis domestica* | *Metazoa › Chordata › Mammalia › Didelphimorphia* | Gray short-tailed opossum | **MdPIEZO2** | **2857** | **35** | XP_007487782.1 | NCBI | https://www.ncbi.nlm.nih.gov/protein/XP_007487782.1 |
| *Monodelphis domestica* | *Metazoa › Chordata › Mammalia › Didelphimorphia* | Gray short-tailed opossum | **MdPIEZO3** | **2111** | **23** | XP_016285057.1 | NCBI | https://www.ncbi.nlm.nih.gov/protein/XP_016285057.1 |
| *Mus musculus* | *Metazoa › Chordata › Mammalia › Rodentia* | House mouse | **MmPIEZO1** | **2546** | **30** | NP_001032375.1 | NCBI | https://www.ncbi.nlm.nih.gov/protein/NP_001032375.1 |
| *Mus musculus* | *Metazoa › Chordata › Mammalia › Rodentia* | House mouse | **MmPIEZO2** | **2824** | **34** | NP_001034574.4 | NCBI | https://www.ncbi.nlm.nih.gov/protein/NP_001034574 |
| *Oryzias latipes* | *Metazoa › Chordata › Actinopterygii › Beloniformes* | Japanese rice fish | **OlPIEZO1** | **2527** | **34** | XP_023819119 | NCBI | https://www.ncbi.nlm.nih.gov/protein/XP_023819119 |
| *Oryzias latipes* | *Metazoa › Chordata › Actinopterygii › Beloniformes* | Japanese rice fish | **OlPIEZO2** | **3001** | **35** | XP_023820679.1 | NCBI | https://www.ncbi.nlm.nih.gov/protein/XP_023820679.1 |
| *Oryzias latipes* | *Metazoa › Chordata › Actinopterygii › Beloniformes* | Japanese rice fish | **OlPIEZO3** | **2554** | **28** | XP_023812003.1 | NCBI | https://www.ncbi.nlm.nih.gov/protein/XP_023812003.1 |
| *Python bivittatus* | *Metazoa › Chordata › Reptilia › Squamata* | Burmese python | **PbPIEZO1** | **2317** | **30** | XP_025024610.1 | NCBI | https://www.ncbi.nlm.nih.gov/protein/XP_025024610.1 |
| *Python bivittatus* | *Metazoa › Chordata › Reptilia › Squamata* | Burmese python | **PbPIEZO2** | **2770** | **35** | XP_007433600.1 | NCBI | https://www.ncbi.nlm.nih.gov/protein/XP_007433600.1 |
| *Python bivittatus* | *Metazoa › Chordata › Reptilia › Squamata* | Burmese python | **PbPIEZO3** | **1914** | **20** | XP_025024760.1 | NCBI | https://www.ncbi.nlm.nih.gov/protein/XP_025024760.1 |
| *Vombatus ursinus* | *Metazoa › Chordata › Mammalia › Diprotodontia* | Common wombat | **VuPIEZO1** | **2521** | **36** | XP_027702284.1 | NCBI | https://www.ncbi.nlm.nih.gov/protein/XP_027702284.1 |
| *Vombatus ursinus* | *Metazoa › Chordata › Mammalia › Diprotodontia* | Common wombat | **VuPIEZO2** | **2807** | **35** | XP_027702878.1 | NCBI | https://www.ncbi.nlm.nih.gov/protein/XP_027702878.1 |
| *Vombatus ursinus* | *Metazoa › Chordata › Mammalia › Diprotodontia* | Common wombat | **VuPIEZO3** | **2581** | **25** | XP_027706364.1 | NCBI | https://www.ncbi.nlm.nih.gov/protein/XP_027706364.1 |
| *Xenopus tropicalis* | *Metazoa › Chordata › Amphibia › Anura* | Western clawed frog | **XtPIEZO1** | **2474** | **36** | XP_002933721.3 | NCBI | https://www.ncbi.nlm.nih.gov/protein/XP_002933721 |
| *Xenopus tropicalis* | *Metazoa › Chordata › Amphibia › Anura* | Western clawed frog | **XtPIEZO2** | **2822** | **35** | XP_012820813.2 | NCBI | https://www.ncbi.nlm.nih.gov/protein/XP_012820813 |
| *Xenopus tropicalis* | *Metazoa › Chordata › Amphibia › Anura* | Western clawed frog | **XtPIEZO3** | **2614** | **28** | XP_031750362.1 | NCBI | https://www.ncbi.nlm.nih.gov/protein/XP_031750362.1 |
| ***Opisthokonta*** | | | | | | | | |
| *Capsaspora owczarzaki* | *Opisthokonta › Holozoa › Ministeriida* | | **CoPIEZO** | **2964** | **33** | XP_004347991.1 | NCBI | https://www.ncbi.nlm.nih.gov/protein/XP_004347991.1 |
| *Fonticula alba* | *Opisthokonta › Holomycota › Fonticulida* | | **FaPIEZO** | **4978** | **32** | XP_009493826.1 | NCBI | https://www.ncbi.nlm.nih.gov/protein/XP_009493826.1 |
| *Rozella allomycis* | *Opisthokonta › Holomycota › Fungi › Rozellida* | | **RaPIEZO** | **2151** | **31** | EPZ31590.1 | NCBI | https://www.ncbi.nlm.nih.gov/protein/EPZ31590.1 |
| *Salpingoeca rosetta* | *Opisthokonta › Holozoa › Choanoflagellida* | | **SpPIEZO1** | **2610** | **29** | XP_004997271.1 | NCBI | https://www.ncbi.nlm.nih.gov/protein/XP_004997271.1 |
| *Salpingoeca rosetta* | *Opisthokonta › Holozoa › Choanoflagellida* | | **SpPIEZO2** | **4142** | **32** | XP_004990485.1 | NCBI | https://www.ncbi.nlm.nih.gov/protein/XP_004990485.1 |
| ***Amoebozoa*** | | | | | | | | |
| *Acanthamoeba castellanii* | *Amoebozoa › Centramoebida* | | **AcPIEZO** | **2644** | **37** | XP_004352964.1 | NCBI | https://www.ncbi.nlm.nih.gov/protein/XP_004352964.1 |
| *Acytostelium subglobosum* | *Amoebozoa › Dictyosteliida* | | **AsPIEZO** | **2484** | **35** | XP_012758690.1 | NCBI | https://www.ncbi.nlm.nih.gov/protein/XP_012758690.1 |
| *Dictyostelium discoideum* | *Amoebozoa › Dictyosteliida* | | **DdPIEZO** | **3080** | **35** | XP_640187.1 | NCBI | https://www.ncbi.nlm.nih.gov/protein/XP_640187.1 |
| *Entamoeba histolytica* | *Amoebozoa › Archamoebae* | | **EhPIEZO1** | **2166** | **31** | XP_649449.2 | NCBI | https://www.ncbi.nlm.nih.gov/protein/XP_649449.2 |
| *Entamoeba histolytica* | *Amoebozoa › Archamoebae* | | **EhPIEZO1** | **2401** | **34** | XP_655549.2 | NCBI | https://www.ncbi.nlm.nih.gov/protein/XP_655549.2 |
| *Entamoeba nuttalli* | *Amoebozoa › Archamoebae* | | **EnPIEZO1** | **2260** | **36** | XP_008855214.1 | NCBI | https://www.ncbi.nlm.nih.gov/protein/XP_008855214.1 |
| *Entamoeba nuttalli* | *Amoebozoa › Archamoebae* | | **EnPIEZO2** | **2427** | **36** | XP_008859622.1 | NCBI | https://www.ncbi.nlm.nih.gov/protein/XP_008859622.1 |
| *Planoprotostelium fungivorum* | *Amoebozoa › Variosea* | | **PfPIEZO** | **2547** | **39** | PRP78767.1 | NCBI | https://www.ncbi.nlm.nih.gov/protein/PRP78767.1 |
| ***Haptista*** | | | | | | | | |
| *Emiliania huxleyi* | *Haptista › Isochrysidales* | | **EhPIEZO** | **2614** | **17** | XP_005775450.1 | NCBI | https://www.ncbi.nlm.nih.gov/protein/XP_005775450.1 |
| ***Heterolobosea*** | | | | | | | | |
| *Naegleria gruberi* | *Heterolobosea › Schizopyrenida* | | **NgPIEZO** | **3262** | **39** | XP_002682879.1 | NCBI | https://www.ncbi.nlm.nih.gov/protein/XP_002682879.1 |
| ***Alveolata*** | | | | | | | | |
| *Paramecium tetraurelia* | *Alveolata › Ciliophora › Peniculida* | | **PtPIEZO1a** | **2544** | **38** | XP_001461300.1 | NCBI | https://www.ncbi.nlm.nih.gov/protein/XP_001461300.1 |
| *Paramecium tetraurelia* | *Alveolata › Ciliophora › Peniculida* | | **PtPIEZO1b** | **2552** | **36** | XP_001461126.1 | NCBI | https://www.ncbi.nlm.nih.gov/protein/XP_001461126.1 |
| *Paramecium tetraurelia* | *Alveolata › Ciliophora › Peniculida* | | **PtPIEZO2a** | **2338** | **28** | XP_001454572.1 | NCBI | https://www.ncbi.nlm.nih.gov/protein/XP_001454572.1 |
| *Paramecium tetraurelia* | *Alveolata › Ciliophora › Peniculida* | | **PtPIEZO2b** | **2470** | **31** | XP_001445884.1 | NCBI | https://www.ncbi.nlm.nih.gov/protein/XP_001445884.1 |
| *Paramecium tetraurelia* | *Alveolata › Ciliophora › Peniculida* | | **PtPIEZO2c** | **2744** | **35** | XP_001447671.1 | NCBI | https://www.ncbi.nlm.nih.gov/protein/XP_001447671.1 |
| *Paramecium tetraurelia* | *Alveolata › Ciliophora › Peniculida* | | **PtPIEZO2d** | **2765** | **38** | XP_001437510.1 | NCBI | https://www.ncbi.nlm.nih.gov/protein/XP_001437510.1 |
| *Stylonychia lemnae* | *Alveolata › Ciliophora› Sporadotrichida* | | **SlPIEZO1a** | **2547** | **30** | CDW72113.1 | NCBI | https://www.ncbi.nlm.nih.gov/protein/CDW72113.1 |
| *Stylonychia lemnae* | *Alveolata › Ciliophora› Sporadotrichida* | | **SlPIEZO2a** | **2429** | **32** | CDW91200.1 | NCBI | https://www.ncbi.nlm.nih.gov/protein/CDW91200.1 |
| *Stylonychia lemnae* | *Alveolata › Ciliophora› Sporadotrichida* | | **SlPIEZO2b** | **3089** | **33** | CDW86065.1 | NCBI | https://www.ncbi.nlm.nih.gov/protein/CDW86065.1 |
| *Symbiodinium microadriaticum* | *Alveolata › Dinophyceae › Suessiales* | | **SmPIEZO** | **1940** | **16** | OLQ07774.1 | NCBI | https://www.ncbi.nlm.nih.gov/protein/OLQ07774.1 |
| *Tetrahymena thermophila* | *Alveolata › Ciliophora › Hymenostomatida* | | **TtPIEZO1a** | **2828** | **33** | XP_001017682.2 | NCBI | https://www.ncbi.nlm.nih.gov/protein/XP_001017682.2 |
| *Tetrahymena thermophila* | *Alveolata › Ciliophora › Hymenostomatida* | | **TtPIEZO2a** | **4364** | **35** | XP_001021704.2 | NCBI | https://www.ncbi.nlm.nih.gov/protein/XP_001021704.2 |
| *Tetrahymena thermophila* | *Alveolata › Ciliophora › Hymenostomatida* | | **TtPIEZO2b** | **3287** | **39** | XP_976967.2 | NCBI | https://www.ncbi.nlm.nih.gov/protein/XP_976967.2 |
| *Vitrella brassicaformis* | *Alveolata › Colpodellida* | | **VbPIEZO** | **3315** | **37** | CEL94374.1 | NCBI | https://www.ncbi.nlm.nih.gov/protein/CEL94374.1 |
| ***Euglenozoa*** | | | | | | | | |
| *Euglena gracilis* | *Euglenozoa › Euglenoida › Euglenales* | | **EgPIEZO** | **2510** | **33** | AUG72368.1 | NCBI | https://www.ncbi.nlm.nih.gov/protein/AUG72368.1 |
| *Leishmania panamensis* | *Euglenozoa › Kinetoplastida › Trypanosomatida* | | **LpPIEZO1** | **1867** | **18** | XP_010698337.1 | NCBI | https://www.ncbi.nlm.nih.gov/protein/XP_010698337.1 |
| *Leishmania panamensis* | *Euglenozoa › Kinetoplastida › Trypanosomatida* | | **LpPIEZO2** | **2538** | **33** | XP_010703279.1 | NCBI | https://www.ncbi.nlm.nih.gov/protein/XP_010703279.1 |
| *Leptomonas pyrrhocoris* | *Euglenozoa › Kinetoplastida › Trypanosomatida* | | **LpPIEZO1** | **1842** | **13** | XP_015660634.1 | NCBI | https://www.ncbi.nlm.nih.gov/protein/XP_015660634.1 |
| *Leptomonas pyrrhocoris* | *Euglenozoa › Kinetoplastida › Trypanosomatida* | | **LpPIEZO2** | **2620** | **32** | XP_015659765.1 | NCBI | https://www.ncbi.nlm.nih.gov/protein/XP_015659765.1 |
| *Perkinsela sp.* | *Euglenozoa › Kinetoplastida* | | **PsPIEZO** | **2623** | **29** | KNH04719.1 | NCBI | https://www.ncbi.nlm.nih.gov/protein/KNH04719.1 |
| *Trypanosoma conorhini* | *Euglenozoa › Kinetoplastida › Trypanosomatida* | | **TcPIEZO1** | **2729** | **35** | XP_029232226.1 | NCBI | https://www.ncbi.nlm.nih.gov/protein/XP_029232226.1 |
| *Trypanosoma conorhini* | *Euglenozoa › Kinetoplastida › Trypanosomatida* | | **TcPIEZO2** | **2520** | **39** | XP_029227104.1 | NCBI | https://www.ncbi.nlm.nih.gov/protein/XP_029227104.1 |
| *Trypanosoma theileri* | *Euglenozoa › Kinetoplastida › Trypanosomatida* | | **TtPIEZO1** | **2795** | **37** | XP_028887613.1 | NCBI | https://www.ncbi.nlm.nih.gov/protein/XP_028887613.1 |
| *Trypanosoma theileri* | *Euglenozoa › Kinetoplastida › Trypanosomatida* | | **TtPIEZO2** | **2678** | **37** | XP_028885470.1 | NCBI | https://www.ncbi.nlm.nih.gov/protein/XP_028885470.1 |
| ***Stramenopiles/Heterokonts*** | | | | | | | | |
| *Achlya hypogyna* | *Stramenopiles › Oomycetes › Saprolegniales* | | **AhPIEZO1** | **2808** | **31** | OQR99194.1 | NCBI | https://www.ncbi.nlm.nih.gov/protein/OQR99194.1 |
| *Achlya hypogyna* | *Stramenopiles › Oomycetes › Saprolegniales* | | **AhPIEZO2** | **2774** | **30** | OQR98286.1 | NCBI | https://www.ncbi.nlm.nih.gov/protein/OQR98286.1 |
| *Albugo candida* | *Stramenopiles › Oomycetes › Albuginales* | | **AcPIEZO** | **1991** | **18** | CCI50278.1 | NCBI | https://www.ncbi.nlm.nih.gov/protein/CCI50278.1 |
| *Bremia lactucae* | *Stramenopiles › Oomycetes › Peronosporales* | | **BlPIEZO** | **2558** | **33** | TDH69418.1 | NCBI | https://www.ncbi.nlm.nih.gov/protein/TDH69418.1 |
| *Cafeteria roenbergensis* | *Stramenopiles › Bicosoecida › Bicosoecales* | | **CrPIEZO** | **5346** | **32** | KAA0169618.1 | NCBI | https://www.ncbi.nlm.nih.gov/protein/KAA0169618.1 |
| *Phytophthora rubi* | *Stramenopiles › Oomycetes › Peronosporales* | | **PrPIEZO** | **2863** | **32** | KAE9034249.1 | NCBI | https://www.ncbi.nlm.nih.gov/protein/KAE9034249.1 |
| *Plasmopara halstedii* | *Stramenopiles › Oomycetes › Peronosporales* | | **PhPIEZO** | **2831** | **34** | XP_024586519.1 | NCBI | https://www.ncbi.nlm.nih.gov/protein/XP_024586519.1 |
| *Pythium brassicum* | *Stramenopiles › Oomycetes › Pythiales* | | **PbPIEZO** | **2891** | **30** | TYZ68687.1 | NCBI | https://www.ncbi.nlm.nih.gov/protein/TYZ68687.1 |
| *Saprolegnia parasitica* | *Stramenopiles › Oomycetes › Saprolegniales* | | **SpPIEZO1** | **2773** | **31** | XP_012201597.1 | NCBI | https://www.ncbi.nlm.nih.gov/protein/XP_012201597.1 |
| *Saprolegnia parasitica* | *Stramenopiles › Oomycetes › Saprolegniales* | | **SpPIEZO2** | **2844** | **29** | XP_012197598.1 | NCBI | https://www.ncbi.nlm.nih.gov/protein/XP_012197598.1 |
| *Thraustotheca clavata* | *Stramenopiles › Oomycetes › Saprolegniales* | | **TcPIEZO1** | **3028** | **38** | OQS03242.1 | NCBI | https://www.ncbi.nlm.nih.gov/protein/OQS03242.1 |
| *Thraustotheca clavata* | *Stramenopiles › Oomycetes › Saprolegniales* | | **TcPIEZO2** | **3119** | **30** | OQS07900.1 | NCBI | https://www.ncbi.nlm.nih.gov/protein/OQS07900.1 |

*By TMHMM (http://www.cbs.dtu.dk/services/TMHMM/)
